# A percolation-type criticality threshold controls immune protein coating of surfaces

**DOI:** 10.1101/2024.10.15.618530

**Authors:** Zhicheng Wang, Sahil Kulkarni, Jia Nong, Marco Zamora, Alireza Ebrahimimojarad, Elizabeth Hood, Tea Shuvaeva, Michael Zaleski, Damodar Gullipalli, Emily Wolfe, Carolann Espy, Evguenia Arguiri, Yufei Wang, Oscar A Marcos-Contreras, Wenchao Song, Vladimir R Muzykantov, Jinglin Fu, Ravi Radhakrishnan, Jacob W Myerson, Jacob S Brenner

**Affiliations:** Department of Systems Pharmacology and Translational Therapeutics, Perelman School of Medicine, University of Pennsylvania; Philadelphia, PA, 19104, USA; School of Engineering and Applied Sciences, University of Pennsylvania; Philadelphia, PA, 19104, USA; Center for Computational and Integrative Biology, Rutgers University-Camden; Camden, New Jersey, 08103, United States; School of Biomedical Engineering, Science and Health Systems, Drexel University; Philadelphia, Pennsylvania, 19104, United States; Department of Neurology, Perelman School of Medicine, University of Pennsylvania; Philadelphia, Pennsylvania, 19104, United States; Department of Bioengineering, University of Pennsylvania; Philadelphia, Pennsylvania 19104, United States; Department of Chemical and Biomolecular Engineering, University of Pennsylvania; Philadelphia, Pennsylvania 19104, United States; Penn Institute for Computational Science, University of Pennsylvania; Philadelphia, Pennsylvania 19104, United States; Pulmonary, Allergy, and Critical Care Division, Department of Medicine, Perelman School of Medicine, University of Pennsylvania; Pennsylvania, 19104, United States

## Abstract

When a material enters the body, it is immediately attacked by hundreds of proteins, organized into complex networks of binding interactions and reactions. How do such complex systems interact with a material, “deciding” whether to attack? We focus on the “complement” system of ∼40 blood proteins that bind microbes, nanoparticles, and medical devices, initiating inflammation. We show a sharp threshold for complement activation upon varying a fundamental material parameter, the surface density of potential complement attachment points. This sharp threshold manifests at scales spanning single nanoparticles to macroscale pathologies, shown here for diverse engineered and living materials. Computational models show these behaviors arise from a minimal subnetwork of complement, manifesting percolation-type critical transitions in the complement response. This criticality switch explains the “decision” of a complex signaling network to interact with a material, and elucidates the evolution and engineering of materials interacting with the body.

## Introduction

All signaling networks are complex^1^, but that complexity is greatly amplified when a signaling network interacts with a material from outside the system. As a result of this added complexity, the topic of how biological complex systems interact with materials has been greatly under-studied. Now more than ever, we must understand how complex biological systems interact with materials, due to the accelerating use of advanced materials in clinical practice, ranging from novel coatings of medical devices to the lipid nanoparticles of the COVID-19 vaccines^2–4^. Understanding interactions between materials surfaces and biological complex systems requires measurements ranging from the nanoscale (where physical interactions occur) to the macroscale (where clinically-relevant emergent properties manifest), and combining such experiments with the computational tools of complexity science.

Here we begin such an investigation, focusing on one of the first-evolved and most clinically important biological complex systems: the “complement” system of ∼40 plasma proteins. The first complement proteins emerged >500 million years ago, and complement is still central in most animals, from sea sponges to humans^5^. Complement proteins bind to microbial surfaces, thus marking pathogens for clearance by phagocytes, while also setting off inflammatory signaling networks^6^. This immune mechanism defends against pathogens, but it can also attack nanomedicines, gene therapy vectors, medical devices, and our own cells^7–9^. Additionally, complement causes damage in nearly every inflammatory disease, including stroke, trauma, and acute lung injury, as in COVID-19^10^.

Complement was first discovered in the 1890s, so the biochemical signaling reactions defining the pathway have been characterized in great depth (Supp. Fig C1 illustrates the complex network diagram of complement)^6^. However, in many cases, we still do not understand why complement coats some surfaces but not others^9,11–14^. Well-defined models for how complement distinguishes one surface from another could advance our understanding of immunology and microbiology and guide new approaches to medical needs ranging from gene therapies to dialysis^15^.

In this study, we examine how complement’s interaction with a material changes upon varying a fundamental control parameter of a material, the surface density of potential complement attachment points. Such information could elucidate evolutionary constraints that complement places on the surface density of proteins on microbial and mammalian cells^5^, and could guide design of molecularly engineered surfaces, like nanoparticles and blood-exposed scaffolds and devices, to avoid complement activation^9^. Beyond practical implications, such studies address the general scientific question: How does a complex molecular network (∼40 proteins in the case of complement) respond to the properties of surfaces, in this case making a “decision” of whether or not to attack the material?

To investigate this question, we combine three approaches: 1) We use nanoscale engineering to vary the distance between potential complement attachment points, then measure the effect on complement activation, both *in vitro* and *in vivo*, at multiple levels of the pathway; 2) We develop new single-particle measurements to reveal how complement coats individual nanoparticles featuring surfaces with different spacings between potential complement attachment points; 3) We synthesize our experimental data with computational and mathematical tools from the science of complex systems (complexity science), showing how a subset of the complement network can explain the entire system’s behavior. Collectively, these approaches lead to a central finding: For a given surface, there is a percolation-type criticality threshold of spacing between potential complement attachment points, below which complement densely coats the surface. Thus, these studies address the question of how the complex network of complement “decides” whether to attack a material, and provide a surprising answer: A small subnetwork of complement makes the decision by undergoing a specific type of phase transition in response to the material properties.

## Results

### A threshold distance between potential complement attachment points determines nanoparticle effects in vivo

We started our investigations by examining whether changes in a nanoscale material property could affect *in vivo* macroscale phenomena known to involve the complement system. The nanoscale property we focused on was the surface density of potential complement attachment points. To control the surface density of potential complement attachment sites, we varied the number of proteins conjugated to the surface of nanoparticles. We hypothesized that each surface protein could serve as only a single attachment point for the primary complement opsonin, C3, since C3 is larger (∼190 kDa) than all surface proteins studied here, and thus steric hindrance should prevent more than one C3 from binding to each surface-anchored protein.

As test particles, we initially used liposomes, a clinically tested nanoparticle with lipid membranes resembling those of cells^16,17^. For our initial studies, we chose IgG as the surface-conjugated protein, modeling targeting antibodies used to functionalize nanomedicines in numerous studies^18^. Antibody-liposomes are a broadly interesting test case where we have precise control over surface properties, allowing us to carefully consider relationships between surface properties, broad biological reactions to those properties, and specific complement reactions to those properties. Antibodies such as IgG have unique functions in complement reactions, so we also tested other surface proteins (e.g., Fab fragments in Supplementary Figure 27), and tested conditions that block IgG-specific effects on the complement pathway (e.g., Mg++ EGTA-GVB buffer prevents an IgG’s Fc from activating the classical pathway of complement; Figs 2-5), to isolate general from IgG-specific effects.

We conjugated different numbers of proteins to liposomes to test how intermolecular spacing of surface proteins changes *in vivo* behavior of intravenous (IV) nanoparticles (Supplementary Fig. 1). Liposome size and surface charge did not vary significantly with respect to the tested densities of surface protein (Supplementary Figs. 2-4). The protein-coated liposomes were tested in mice following IV bacterial lipopolysaccharide (LPS) insult. We previously showed that nanoparticles with exposed surface proteins can be taken up in leukocytes that localize inside the lungs’ capillaries following inflammatory insults like IV LPS administration^19^. We found that these nanoparticle-leukocyte interactions were mediated by complement^19^.

Fitting our previous observations, liposomes with 200 IgG per nanoparticle accumulate in the inflamed lungs at a high concentration. However, lung uptake is uniformly low for <100 IgG per liposome (Fig. 1a). Converting the protein density on the liposome surfaces to average intermolecular protein spacing (Supplementary Fig. 5), we find a threshold in lung uptake vs. protein-protein spacing, with lung uptake increasing steeply for spacing ≤ 25 nm (Fig. 1b). Uptake in other organs did not have this same threshold (Supplementary Fig. 6). We used flow cytometry to verify that spacing-dependent lung uptake was the result of nanoparticle uptake in leukocytes (Fig. 1c-d, Supplementary Figs. 7-9). Changing surface protein spacing from 36 nm to 18 nm, we find an order of magnitude increase in nanoparticle uptake in lung leukocytes.

**Figure 1.**
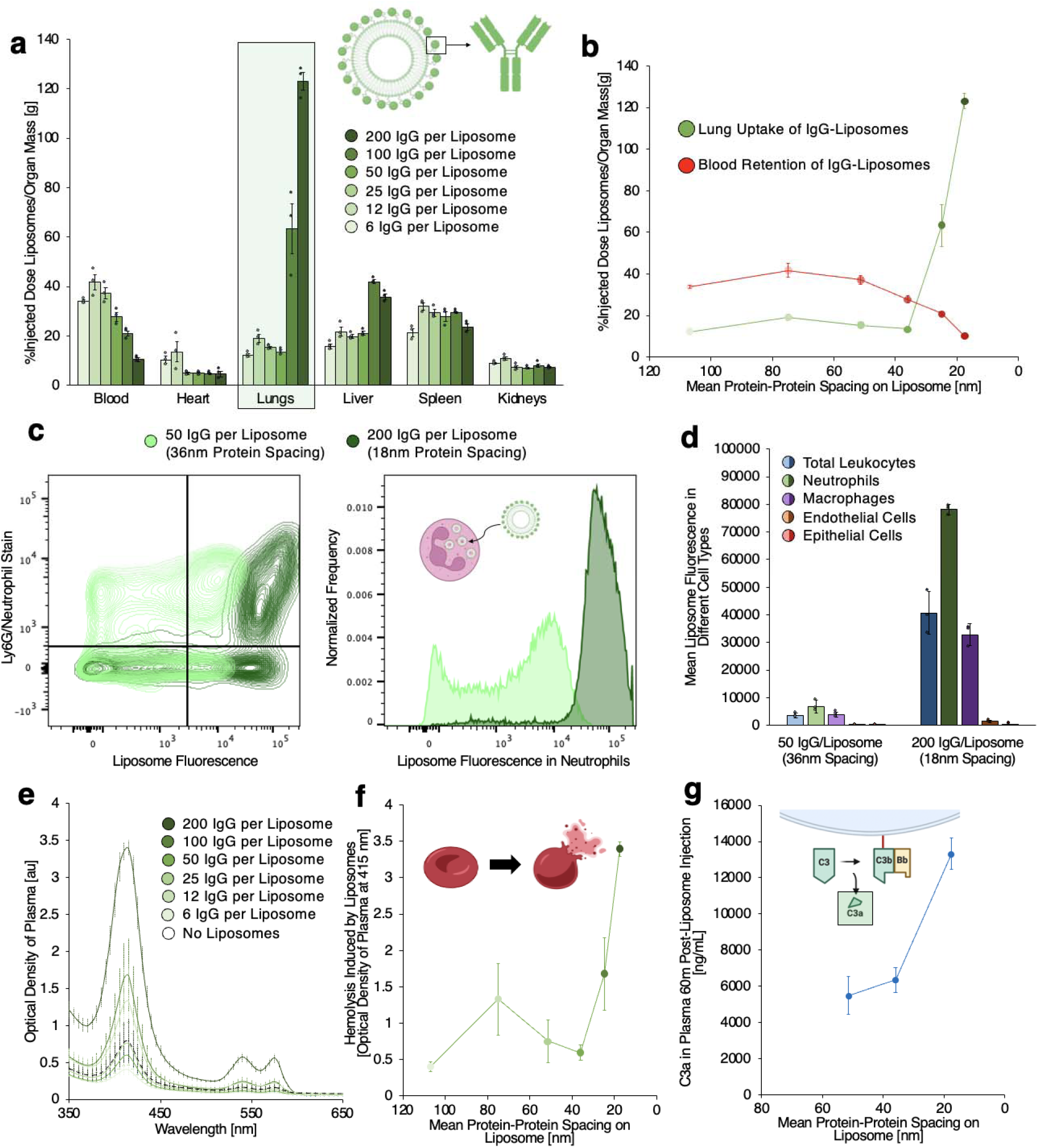
Nanoparticle biodistribution and side effects have thresholds with respect to protein-protein spacing on the nanoparticle surface. **a,** Biodistributions of IgG-coated liposomes show uptake of nanoparticles in the lungs and liver, and retention in the blood, depending on the quantity of IgG on the nanoparticle surfaces. **b,** Lung uptake and blood retention of nanoparticles from **a** as a function of average protein-protein spacing on the nanoparticle surface, showing lung uptake steeply increasing for spacing < 36 nm. **c,** Flow cytometry profiling uptake of protein-liposomes in lung neutrophils. Shifting surface protein spacing from 36 nm to 18 nm results in >10X increase in nanoparticle uptake in neutrophils. **d,** Summarized flow cytometry data shows shifting surface protein spacing from 36 nm to 18 nm significantly increases nanoparticle uptake in total leukocytes, neutrophils, and macrophages. **e,** Optical absorbance spectra of plasma taken from mice receiving intravenous protein-liposomes. Increasing surface protein density on nanoparticles leads to greater optical density at 415 nm, 545 nm, and 575 nm. **f,** Hemolysis, as indicated by optical density of plasma at 415 nm in data from **e**, vs. protein-protein spacing on dosed liposomes. Surface protein spacing < 36 nm leads to a steep increase in hemolysis. **g,** C3a levels, indicating generation of complement complexes that can adhere to nanoparticles, in plasma 60 minutes after injection of protein liposomes, showing increased C3a as protein-protein spacing on the liposomes decreases.

We probed for side effects of our protein-coated liposomes and again found thresholds with respect to surface protein spacing. Hemolysis, an established side effect of complement activation^20^, had a threshold with respect to surface protein spacing matching that of nanoparticle lung uptake (Fig. 1e-f, Supplementary Fig. 10). Nanoparticle effects on hemoconcentration (a proxy for systemic capillary leak) and platelet count had similar threshold responses (Supplementary Figs. 11-13). Finally, a direct measurement of complement activation, serum C3a, also showed a similar threshold (Fig. 1g, Supplementary Fig. 14). Thus, varying a nanoscale parameter (surface density of potential complement attachment points) over a small range produces a sharp threshold for macroscale effects, including pathologies.

### Single-particle measurements show surface protein spacing is an on/off switch for complement coating of nanoparticles

We scaled our measurements from the macroscale down to the nanoscale, investigating whether a sharp threshold would be observed in complement deposition onto individual nanoparticles. We measured deposition of the core opsonin of the complement system, C3. C3 can be cleaved via three different routes (the classical, lectin, and alternative pathways), to produce C3b, which possesses an exposed reactive thioester (Fig. 2a)^21^. That thioester can adhere to nucleophiles on surfaces, forming a covalent C3b-surface adduct.

**Figure 2.**
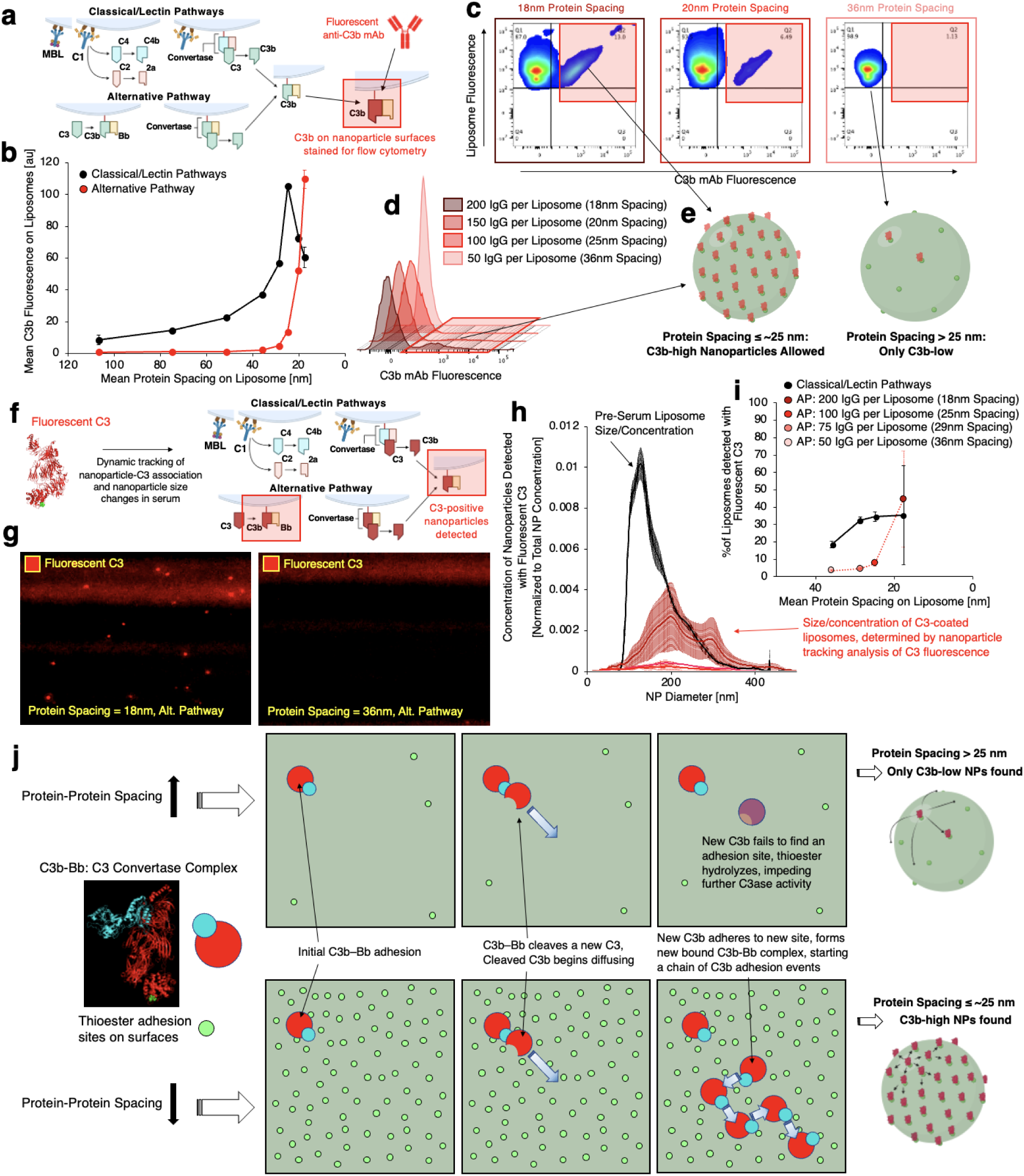
Protein-protein spacing on nanoparticles is an on/off switch for complement C3 deposition on nanoparticle surfaces. **a,** Schematic describing mechanisms of C3 deposition and flow cytometry method for detection on nanoparticle surfaces. **b,** C3b fluorescence on protein-nanoparticles vs. protein-protein spacing on the nanoparticles, showing a precipitous increase in alternative pathway C3b adhesion below a threshold spacing of ∼29 nm. **c,** Two-dimensional plots of nanoparticle fluorescence vs. alternative pathway C3b fluorescence for protein-nanoparticles with different surface protein spacing, with high-C3b populations detected for lower surface protein spacing (red boxes). **d,** Histograms of alternative pathway C3b fluorescence per nanoparticle for nanoparticles with different surface protein spacing, with high-C3b populations highlighted for low surface protein spacing. **e,** Schematics of C3b-high (low surface protein spacing) and C3b-low nanoparticles. **f,** Schematic, as in (a), showing method for nanoparticle tracking analysis of C3b deposition on nanoparticle surfaces. **g,** Still images of fluorescent C3 signal in NTA of serum+fluorescent C3+protein-nanoparticle samples for protein-nanoparticles with 18 (left) vs. 36 (right) nm surface protein spacing, showing C3-positive nanoparticles for 18 nm, but not 36 nm surface protein spacing. **h,** Size distributions showing total concentration vs. size of protein-nanoparticles (black curve) and concentration of C3-positive nanoparticles vs. size for serum+fluorescent C3+protein-nanoparticles with different surface protein spacing (red curves). **i,** % of protein-nanoparticles that were C3-positive vs. protein-protein spacing on the nanoparticles, indicating a threshold pattern curve shape for alternative pathway C3 deposition matching the curve shape in **b**. **j,** Schematic of proposed mechanism by which surface protein spacing regulates C3-surface interactions. Top panels: proposed dynamics of C3b generation and adhesion on surfaces with large protein-protein spacing. Leftmost top panel: Generation of an initial C3b-Bb complex on the surface via tickover. Middle top panel: Generation of a new C3b on the surface via autocatalytic cleavage of a fluid phase C3 by the initial C3b-Bb complex. Rightmost top panel: Proposed mechanism for failure of C3b to adhere to the surface with wide protein-protein spacing, wherein the newly generated C3b fails to diffuse and adhere to a new protein because its thioester is hydrolyzed before the C3b can colocalize with a new adhesion site. Lower panels: proposed dynamics of C3b generation and adhesion on surfaces with close protein-protein spacing. Rightmost lower panel: C3b generated on the surface readily finds a nearby adhesion site, before hydrolysis. The newly adhered C3b in turn forms new C3b-Bb complexes, in turn catalytically generating more C3b near the surface, which can again readily adhere to the surface at new sites, yielding a positive feedback loop of generation and adhesion of C3b on the surface with high protein density.

We developed single-particle measurement techniques to assess C3b adhesion to nanoparticle surfaces as a function of surface protein spacing. First, we stained for C3b on protein-liposome surfaces in flow cytometry measurements (Fig. 2a, Supplementary Figure 15). Protein-coated fluorescent liposomes were synthesized and incubated with serum *in vitro*. Protein-coated liposome + serum samples were stained with a fluorescent anti-C3b antibody and C3b fluorescence on the nanoparticles was quantified. C3b per liposome vs. intermolecular spacing on the liposomes shows increasing C3b deposition for spacing ≤ 28 nm (Supplementary Figs. 16-17). We separated C3b signals due to the alternative pathway vs. the classical/lectin pathways (Fig. 2a) by comparing experiments with a buffer that blocks the classical and lectin pathways (Mg++ EGTA-GVB) vs. a buffer that allows all 3 pathways of complement (PBS). These experiments show an alternative pathway-driven threshold in C3b deposition vs. intermolecular spacing. As intermolecular spacing decreases below ∼25 nm, alternative pathway C3b deposition precipitously increases (red trace in Fig. 2b), while C3b deposition due to the classical and lectin pathways levels off (black trace). The fact that the sharp threshold is driven by the alternative pathway shows that the threshold phenomenon is not IgG-specific, since the alternative pathway does not react to the Fc of IgG (only the classical pathway does). Thus, we found a threshold surface protein spacing of ∼25nm for C3b deposition on nanoparticles *in vitro*, similar to that for macroscale pathologies *in vivo* (Fig 1).

The threshold shown in Figure 2b (red trace) was found by taking a mean of the C3b signal across a population of nanoparticles. We initially assumed that the amount of C3b deposition would be evenly distributed across a population of nanoparticles. However, when we examined two-dimensional plots of nanoparticle fluorescence vs. alternative pathway C3b fluorescence, we found two divergent populations of nanoparticles (Figs. 2c-d). Among nanoparticles with a below-threshold spacing between surface proteins (< ∼25 nm), most particles have low C3b deposition, but a subpopulation has a high amount of C3b deposition (red boxes in Figs. 2c-d). Thus, for nanoparticles that have below-threshold spacing of their surface proteins (<25 nm), C3b does not deposit evenly across a population, but rather a fraction of particles gets heavily coated in C3b, while the majority have low C3b deposition. That is, sub-threshold nanoparticles can be C3b-high, but are not always C3b-high (Fig 2e, left panel). For a population of nanoparticles with above-threshold spacing between proteins (>25 nm), none of the nanoparticles get thoroughly coated in C3b (Fig 2e, right panel).

We verified our findings with a second technique for single-particle detection of C3b deposition. Protein-liposomes were incubated in serum doped with fluorophore-tagged recombinant C3. Nanoparticles with C3 fluorescence were quantified with nanoparticle tracking analysis (NTA) (protocol depicted in Fig. 2f)^22^. We confirmed that the nanoparticles themselves would not produce fluorescence signal interfering with C3 detection (Supplementary Movies 1-2), that fluorescent C3 would not react with endogenous vesicles in serum (Supplementary Movies 3-4), and that the concentration and size distribution of endogenous vesicles in serum were different from the concentration and size distribution of our protein-nanoparticles (Supplementary Fig. 18). Therefore, detection of particles with fluorescent C3 in our assay only indicated reaction of C3 with protein-liposome surfaces (Supplementary Movies 5-12). The quantity of protein-liposomes detected with fluorescent C3 varies with protein spacing on the liposomes (Figs. 2g-h, Supplementary Figs. 19-21). Focusing on C3 deposition via the alternative pathway, we observe a pattern precisely matching our findings with flow cytometry detection of C3b deposition, where decreasing surface protein spacing below 25 nm induces a precipitous increase in C3 addition to the nanoparticle surface (Fig 2i, Supplementary Fig. 22).

In summary, these single-particle measurements show two surprising phenomena for how the alternative pathway of complement works: 1) There is a threshold value of protein-protein spacing on nanoparticles, below which C3b deposition increases by orders of magnitude (Fig 2b & 2i). This holds for measurements averaged across a population of nanoparticles. 2) For nanoparticles with below-threshold protein-protein spacing, only a fraction of particles get heavily C3b-opsonized. To explain these two effects, we hypothesize the following mechanism (Fig. 2j): An initial C3b can adhere to a surface protein via the slow process of “tickover” (i.e., C3b generated by spontaneously generated C3(H_2_0)Bb). The resulting C3b-surface adduct can catalyze production of new C3b molecules, which can then diffuse to form adducts on their nearest-neighbor potential adhesion points (nearby surface proteins). But if the nearest-neighbor adhesion points are too far away (>25 nm), a soluble C3b’s thioester will hydrolyze before it can attach to a nearest-neighbor attachment point. On the other hand, if a C3b-surface adduct has very close (<25 nm) nearest-neighbor attachment points, the newly generated soluble C3b molecules can attach to nearby surface proteins, and those C3bs will in turn generate new C3b-surface adducts, that will in turn release more soluble C3b molecules that diffuse and attach to their nearest-neighbors, and so on. This process of “C3b spreading” across a surface should allow for “runaway events,” where C3b spreads from adhesion site to adhesion site to coat an entire nanoparticle (Fig 2j, bottom right). The runaway events of C3 spreading should only occur in a population of nanoparticles with below-threshold protein-protein spacing (< 25 nm), and such events are rare, due to the probabilistic nature of tickover. Such a mechanism of runaway C3 spreading events (Fig 2j) might explain all the phenomena seen in our single nanoparticle measurements.

### Sharp thresholds are present at multiple levels of the signaling cascade and they are invariant upon normalizing to the number of proteins on the nanoparticles

We determined whether the sharp thresholds observed with C3b deposition are present at other levels in the complement signaling cascade (Fig 3a, f, i). To do so, we performed *in vitro* complement activation experiments on protein-liposomes as in Figure 2, and then measured the soluble (solution-phase) products C3a and C5a by ELISA, and the surface-bound protein properdin using flow cytometry (as done for C3b).

**Figure 3.**
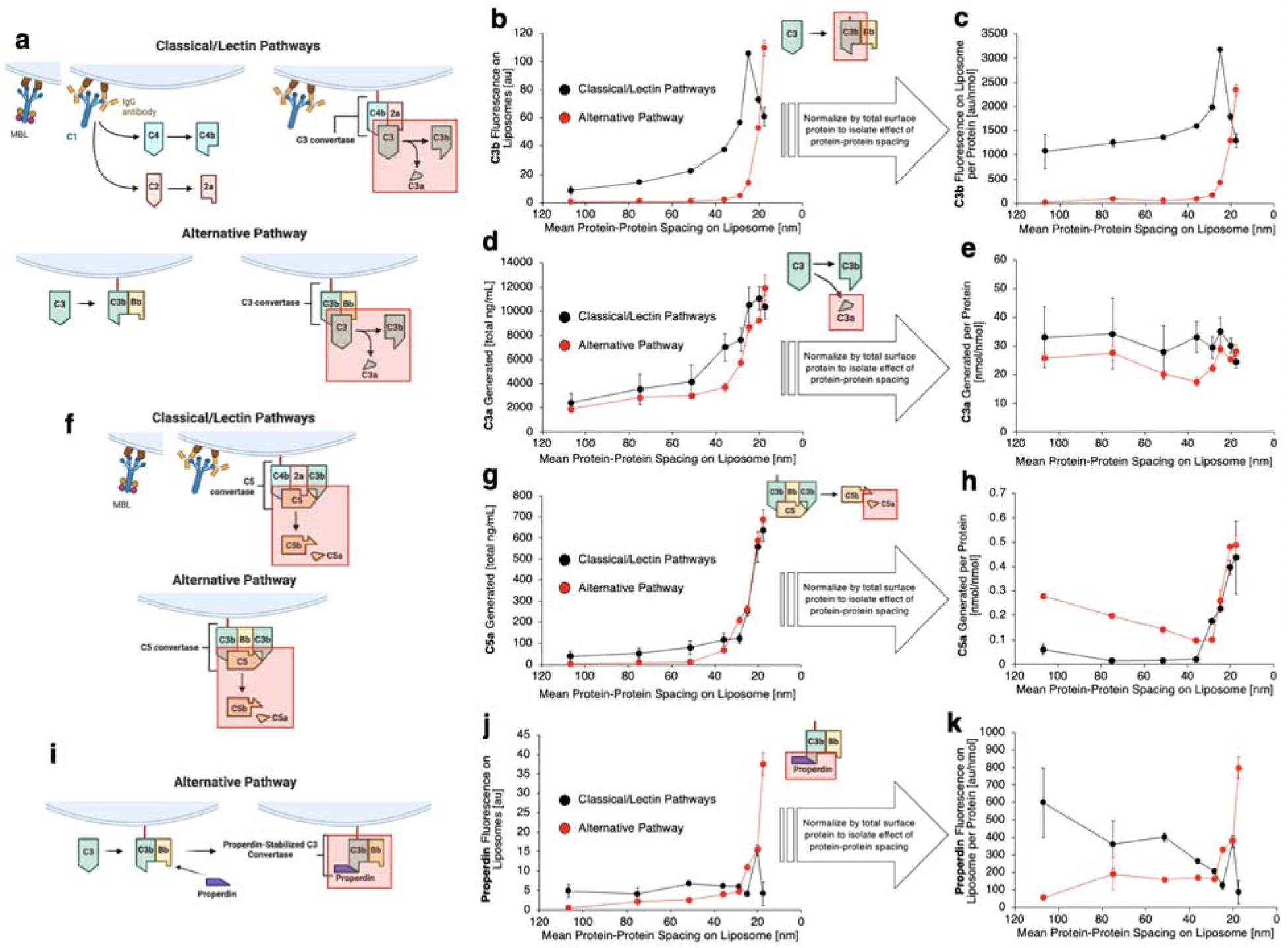
Sharp thresholds are present at multiple levels of the complement cascade, and they are invariant upon normalization to the number of proteins on the liposomes. **a,** Schematics of C3 convertase assembly for the different complement pathways, with C3 cleavage highlighted. **b-c,** C3b fluorescence on liposomes vs. protein-protein spacing on the liposomes (as in Figure 2b), showing invariance of the threshold with normalization to number of proteins per liposome. **d-e,** C3a production in serum doped with protein-liposomes vs. protein spacing on the liposomes, showing C3a production, a process that does not require C3-surface conjugation, has no threshold when normalized to the number of surface proteins per liposome. Note that C3a can be formed by cleavage of C3, even if the resulting C3b never forms a surface adduct. **f,** Schematics of C5 convertase assembly for the different complement pathways, with C5 cleavage highlighted. **g-h,** Data as in (d-e), but for C5a production vs. protein spacing, showing a threshold that is invariant with normalization to protein concentration. **i,** Schematic of properdin’s role of stabilization of the alternative pathway C3 convertase. **j-k,** Data as in **b-c**, but for detection of properdin on nanoparticle surfaces, showing a threshold that is invariant with normalization to protein concentration.

In our analysis, we normalized these results to the number of surface proteins on the liposomes This normalization is especially informative, because protein-liposomes with lower distances between surface proteins also have more proteins on their surface than liposomes with a higher distance between proteins (the number of liposomal surface proteins scales with 1/x^2^, where x is the mean distance between proteins). Thus, we normalized to calculate the number of active complement molecules (C3a, C5a, etc) produced per surface protein on the liposomes. This normalized metric has the clearest meaning for activity of the complement pathway compared to a surface property.

For C3b-surface adducts, upon taking the raw data (Fig 3b) and normalizing it to the number of surface-proteins on the liposomes, the alternative pathway threshold is unchanged (Fig 3c, red, Supplementary Fig. 25). Thus, the C3b threshold is not simply a product of having more surface-proteins on the liposomes. For C3a, the results are quite different. While there appears to be a threshold in the un-normalized data (Fig 3d, red), that threshold disappears upon normalization (Fig 3e, red; Supplementary Fig. 26a; Supplementary Fig. 27). At first, this difference between C3b and C3a may seem surprising, because every C3 cleavage results in one C3a and one C3b (Fig 3a). However, as noted in Figure 2j, many C3b molecules never adhere to a surface, as the C3b thioester may hydrolyze before encountering a surface nucleophile. Our C3b measurements (by flow cytometry) only examine C3b-surface adducts, not hydrolyzed C3b in the bulk solution. By contrast, all C3a molecules remain in solution and are measurable by ELISA. That C3b has a sharp threshold after normalization, while C3a does not, hints that the threshold phenomenon hinges on complement molecules binding to surfaces.

We therefore examined whether sharp thresholds exist for other complement products that depend on surface adducts. C5a is produced by complexes of surface-adhered C3b with other complement components (Fig 3f). As for C3b-surface adducts, we found that C5a has a sharp threshold before and after normalization to the number of surface proteins (Fig 3g, h, Supplementary Fig. 26b). C5b-9 is a byproduct of C5a generation, and we also observe a normalization-invariant threshold when tracing C5b-9 (Supplementary Fig. 28).

Properdin is a protein that binds to and stabilizes surface C3b complexes (Fig 3i). As with C3b and C5a, properdin has a sharp threshold before and after normalization (Figs. 3j-k, Supplementary Figs. 29-30). Since properdin is important to stabilizing the C3b-surface adducts, we inhibited properdin and examined effects on the C3b and C3a data. We found that inhibiting properdin, and thus destabilizing C3b-surface adducts, removes the threshold in C3b surface adhesion (Supplementary Figs. 31-33). Thus, the sharp threshold appears in multiple levels of the complement cascade, but only for complement products that require surface-adducts for their formation and under conditions where C3b-surface adducts are stable.

### Complement responses to bacteria are modulated by surface nucleophile density

The above reactions of complement to surface protein spacing were demonstrated for synthesized liposomes. We looked for a similar pattern in complement reactions to bacteria with variable surface protein density.

Nucleophiles on E. coli were blocked with methyl-terminated poly(ethylene) glycol chains. Since C3b adheres to nucleophiles, blocking nucleophiles on bacteria corresponds to blocking C3b from adhering to sites on surface proteins. By limiting the likelihood of C3b adhesion to some surface proteins, blocking nucleophiles yields an effective increase in the spacing between surface proteins (Fig. 4a). Quantitative ninhydrin assay (Supplementary Fig. 34) measured methylation-induced blocking of surface nucleophiles. Based on the average number of nucleophiles per protein and the average E. coli surface area, we determined an “effective” surface protein spacing on the modified E. coli, reflecting the number of blocked nucleophiles (Fig. 4b, Supplementary Fig. 35).

**Figure 4.**
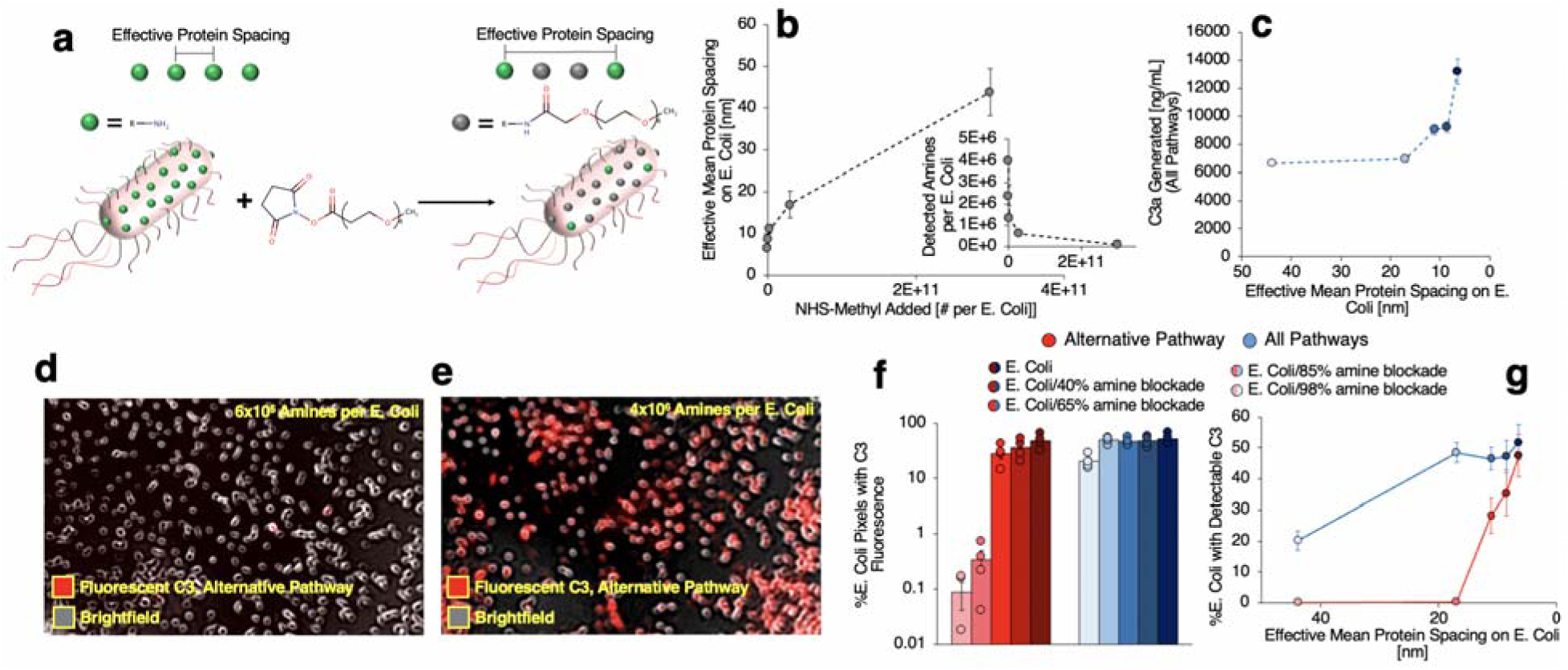
Modulation of effective surface protein spacing on E. coli produces a threshold effect on complement response and alternative pathway C3 adhesion to the microbe surfaces. **a,** Scheme for chemically blocking nucleophiles on E. coli with methyl-terminated poly(ethylene) glycol (PEG) groups, demonstrating how this strategy causes an effective increase in protein-protein spacing on the E. coli. **b,** Effective mean protein spacing on E. coli vs. methyl-PEG groups added to the E. coli, as determined by modified ninhydrin assay for amine detection. Inset: Number of amines detected per E. coli vs. methyl-PEG groups added to the E. coli. **c,** C3a production in serum+modified E. coli vs. effective protein spacing on E. coli, showing increased C3a production at lower surface protein spacing. **d-e,** Micrographs of modified E. coli after incubation with serum + fluorescent C3. Alternative pathway deposition of C3 on the E. coli is detectable for high amine concentration/low surface protein spacing (**e**), but not low amine concentration/high surface protein spacing (**d**). **f-g,** Quantification of C3 deposition on E. coli vs. effective protein spacing on E. coli, measured in images as in **d-e**, showing C3 deposition via the alternative pathway has a threshold dependence on effective protein spacing on the E. coli.

Modified E. coli were incubated with serum and C3a concentration was measured, as in our experiments with nanoparticles. C3a production induced by E. coli vs. effective protein-protein spacing on E. coli has a pattern resembling that for the analogous nanoparticle data (Fig. 4c). There are differences between the pattern for E. coli and the pattern for nanoparticles: 1) The lectin pathway responds to polysaccharides on bacterial surfaces^21^, so we expect complement to react to E. coli surfaces even without surface proteins. Indeed, there is a high level of C3a production even for wider protein-protein spacing on E. coli. 2) The protein spacing threshold for the C3a response to E. coli is < 10 nm, under half the protein-protein spacing threshold on nanoparticles. The difference in threshold spacing could result from the different properties of the E. coli surface proteins, from the different curvature of the E. coli surface, or from inaccuracies in our estimate of effective protein-protein spacing on the modified E. coli. However, the pattern remains: Below a specific protein-protein spacing, there is a precipitous increase in complement response due to serum-E. coli interactions.

Further paralleling our experiments with nanoparticles, we measured C3b adhesion to modified E. coli. Flow cytometry showed that more C3b accumulates on E. coli when the effective E. coli surface protein spacing is lower. Bound C3b vs. surface protein spacing varied more steeply when isolating the alternative pathway response to the modified E. coli (Supplementary Fig. 36). Similar trends were observed for properdin adhesion to E. coli (Supplementary Fig. 37). We also imaged modified E. coli after incubation with serum containing fluorescent C3 (Fig. 4d-e, Supplementary Fig. 38). When all complement pathways are active, microscopy detects robust amounts of C3 on E. coli, regardless of surface protein spacing (Fig. 4f-g, blue bars/curves). However, a threshold is evident for the alternative pathway, where C3 adhesion to E. coli is not detectable by microscopy for wider surface protein spacing, but C3 fluorescence on E. coli steeply increases for surface protein spacing < ∼17 nm (Fig. 4f-g, red bars/curves).

### Fixed protein spacing controls complement responses to planar protein arrangements

Our results thus far have considered complement reactions to particle surfaces where we control the average protein spacing, but do not precisely control the spacing of each protein pair on a particle. We used self-assembled DNA origami to precisely define the positions of four IgGs on 90 nm × 60 nm DNA planes (Fig. 5a, Supplementary Fig. 39)^23^. DNA strands for anchoring protein were spaced in rectangular conformations of 31 nm × 20 nm, 73 nm × 20 nm, or 73 nm × 40 nm. Placement of these strands and conjugation of protein to the anchors was confirmed by atomic force microscopy (Fig. 5b-d, Supplementary Figs. 40-42).

**Figure 5.**
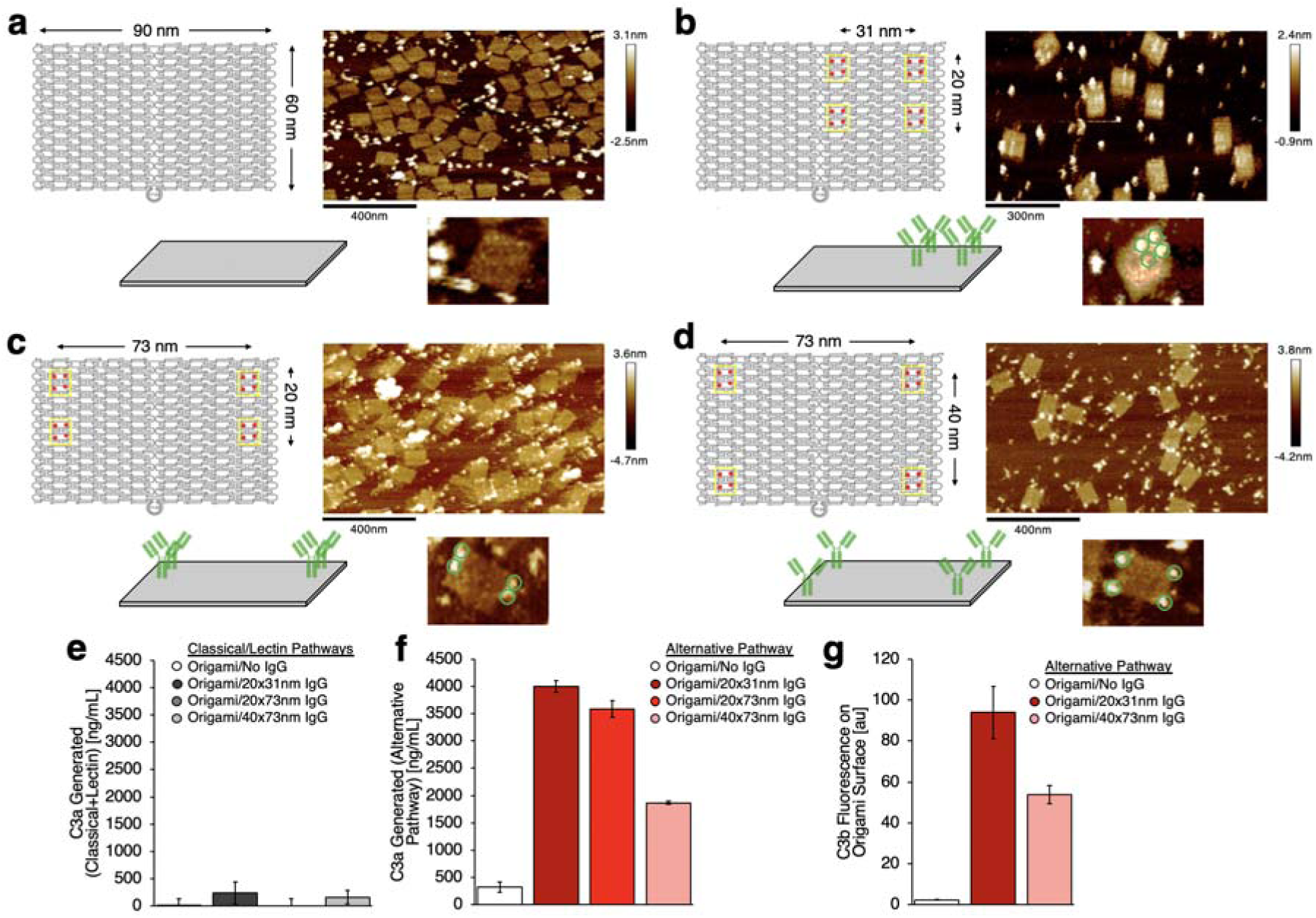
Engineering of protein arrangement on DNA origami surfaces modulates complement responses to those surfaces. **a,** DNA origami design for 90 nm × 60 nm planes (left panel), with geometry confirmed by atomic force microscopy (AFM, right panel). Origami planes were incubated with alkyne-functionalized IgG (bright puncta in the micrograph) to confirm lack of non-specific adhesion to the origami (lower panel). **b-d,** Left panels: DNA origami designs with four clusters (yellow boxes in schematics) of four azide-terminated anchor sequences (red points in schematics), with the clusters set at the corners of 31 nm × 20 nm (**b**), 73 nm × 20 nm (**c**), or 73 nm × 40 nm (**d**) rectangles. Right panels: AFM micrographs confirming geometry of origami planes after incubation with alkyne-functionalized IgG. Lower panels: conformations of IgG conjugated to origami surfaces. **e,** C3a production via the classical/lectin pathways after incubation of origami in serum. **f,** C3a production via the alternative pathway after incubation of origami in serum. **g,** C3b detected via fluorescent antibody stain after incubation of surface-bound origami with serum.

Origami planes with proteins were incubated with serum and C3a production was measured. For all protein arrangements, origami induced negligible C3a production via the classical and lectin pathways, so the complement response to the origami was dominated by the alternative pathway (Fig. 5e). 31 nm × 20 nm and 73 nm × 20 nm protein arrangements both induced high amounts of C3a production via the alternative pathway. But increasing the minimum protein spacing to 40 nm (73 nm × 40 nm conformation) halved the C3a production (Fig. 5f).

We also detected C3b adhesion on origami-IgG fixed to glass surfaces. After exposing the surface-bound origami to serum and washing, fluorescent C3b antibodies measure roughly double the amount of C3b on 31 nm × 20 nm IgG arrangements vs. 73 nm × 40 nm arrangements. Negligible quantities of C3b were detected on DNA origami without IgG (Fig. 5g). The clear result of our experiments with DNA origami: With exactly the same quantity of protein on exactly the same surfaces, changing *only* the intermolecular spacing of the surface proteins dramatically changes the complement response to and complement adhesion to the surface.

### Computational models show thresholding of complement-surface interactions emerge from percolation-type critical points

We used diverse computational models to determine if the known biochemical reactions of the complement network can produce the emergent properties^24^ found in experiments with nanoparticles: 1) thresholds in complement activation vs. surface protein spacing, and 2) approximately all-or-none C3b coating of surfaces with below-threshold surface protein spacing (divergent subpopulations). First, we adapted an ordinary differential equation (ODE) model of all reactions in, and impinging upon, the alternative pathway of complement^25^. The model uses published reaction parameters that were either experimentally measured or estimated based on prior experimental data sets (Supplementary Computational Model Details). The ODE model predicts curves of C3a production and C3b surface adhesion vs. protein-protein spacing that are strikingly similar to those measured experimentally, with similar protein-protein distance thresholds, even though no protein sizes were explicitly encoded into the model (Fig. 6a-b).

**Figure 6.**
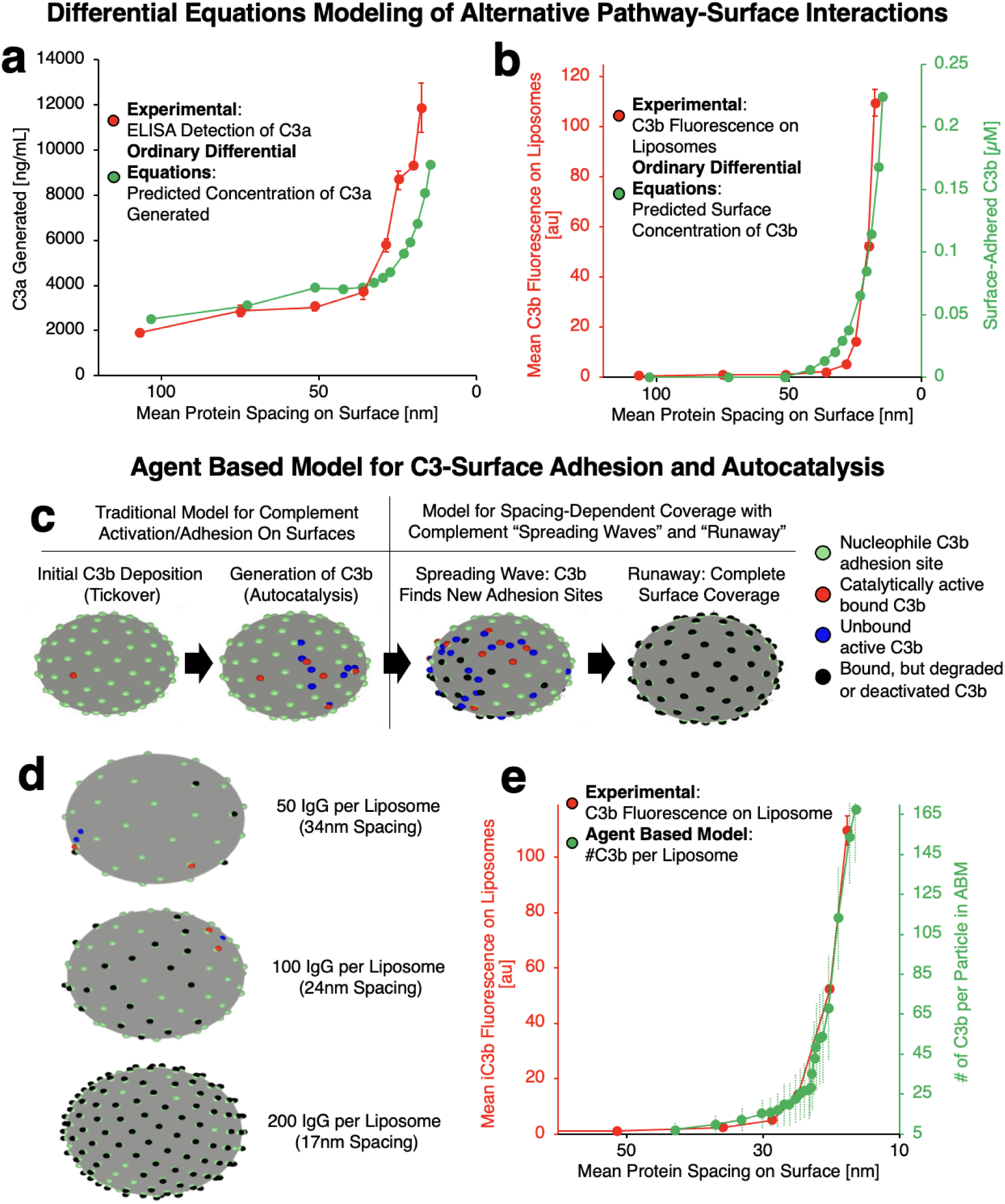
Effects of surface protein spacing in computational models of complement responses to surfaces match experimental findings. **a-b,** Comparisons between experimental findings and ordinary differential equations predictions of (**a**) alternative pathway C3a production in response to and (**b**) C3b adhesion to surfaces coated with IgG arranged with different protein-protein spacings. **c,** Schematic describing construction and qualitative findings of an agent-based model (ABM) for C3 interactions with protein-coated surfaces. **d,** Frames showing a single time point in ABM simulations of C3 interactions with particles coated with IgG arranged with 34 nm, 24 nm, and 17 nm spacing, indicating complete coverage of the particle surface with C3 (black dots) for the smallest IgG-IgG spacing. **e,** Comparison of experimental and ABM simulation data showing quantity of C3b adhered to IgG-coated nanoparticles vs. protein-protein spacing on the particles. Experimental data are reproduced from Fig. 2b and Fig. 3b.

The ODE model describes bulk-scale complement responses to surfaces, but experiments throughout this study indicate that single-particle and nanoscale C3 interactions with surfaces are a determinant of the complement vs. protein-protein spacing threshold. Therefore, to accompany the ODE model, we designed a lattice-based agent-based model (ABM), capturing aspects of single particle scale C3-surface interactions that we hypothesized might dictate the threshold behavior. In the ABM, the agents are C3bs. C3b can spontaneously deposit on a surface, simulating tickover^12,13^. The surface-bound C3b agent (Fig. 6c, red dots) can create new C3b agents, simulating autocatalysis^6,13^. Newly generated C3b agents (Fig. 6c, blue dots) will then diffuse near the surface and can either find a binding site on the surface or become ineligible to bind to the surface, simulating hydrolysis of the surface-binding thioester on C3b^21^. If the new C3b agent does adhere to the surface, it forms a new autocatalytic site that can in turn generate new C3b agents that undergo the same probabilistic diffusion/binding/decay processes. Over time, the autocatalytic agents can lose their ability to generate new C3b agents, simulating degradation of surface-bound C3 convertases (Fig. 6c, black dots). All these processes occur on a quasi-two-dimensional spherical surface covered with a lattice of complement attachment sites (Fig. 6c, green dots). The spacing of the complement attachment sites was varied, to simulate different densities of surface proteins on nanoparticles.

The ABM recapitulates both phenomena observed experimentally: A sharp threshold of complement activation (on a population level), and approximately “all-or-none” C3b coating of individual nanoparticles (divergent subpopulations). Additionally, ABM yields the “spreading C3b waves” we hypothesized based on the experimental results (Fig 2j). Below a threshold attachment site spacing, attachment of C3b agents to one site results in generation of new C3b agents that rapidly attach to neighboring sites (Fig. 6c, second from right panel). When these areas of C3b-occupied sites efficiently grow, we observe “runaway” events on individual particles: a C3b spreading wave continues until nearly every attachment site on a nanoparticle is occupied by a C3b (Fig. 6c, rightmost panel). Runaway events were stochastically observed, but only occurred when attachment site spacing was below a threshold value (Fig. 6d, Supplementary Movies 13-15). These runaway events appear to correspond to the population of heavily C3b-coated nanoparticles observed experimentally (Fig 2c, d).

The ODE and ABM models, while very different types of models, both clearly recapitulate our experimental data. Importantly, the parameters of these models were based on prior data in the literature, and they were not fit to our data. Thus, these models show that the complement network’s topology, reactions rates, and concentrations are sufficient to produce a sharp threshold in complement activation as we vary a surface property (distance between C3 attachment points). Additionally, these models recapitulate our newly described C3 spreading waves and runaway events. The fact that such phenomena had not been described before shows the power of studying a signaling pathway with the tools of surface science and complexity science (computational modeling of complex systems).

However, thus far, these models have only recapitulated our experimental findings, without answering the question: *Why* is there a threshold? Sharp thresholds for signaling pathway-surface interactions in biology have not been frequently described, so we seek to understand what properties of a complex signaling network would produce such a threshold.

To address the question, we take three steps below. First, from the topology of the biochemical reactions of the ODE model, we looked for a *minimal subnetwork* of chemical reactions that would produce a sharp threshold. Second, we compared that subnetwork to other complex systems known to have sharp thresholds upon varying a control parameter. Third, we checked if our minimal subnetwork and ABM would produce the same dynamical properties seen by other complex systems in their class. Together, these analyses lead to a singular conclusion: There is a minimal subnetwork within the complement cascade that has a percolation-type critical threshold in response to surface properties.

We examined the topology of the entire complement network to identify a minimal subnetwork of reactions that might manifest a sharp threshold. This is not a trivial task, as the ODE model used in Figure 6 includes 107 coupled ODEs with 74 parameters. As shown in Figure 7a, we grouped ODEs in the system to narrow our focus to autocatalytic C3b generation, C3b-surface attachment, and concentration of attachment sites on a surface. This reduced ODE model therefore hews to the physical model proposed in Figure 2j and tracks 3 species’ concentrations (Fig 7b): 1) [Sites] = the concentration of potential C3b attachment sites on a surface; 2) [C3b_bound_] = concentration of C3b-surface-site adducts (i.e., sites attached to a bound C3b, able to catalyze production of more C3bs); 3) [C3b] = soluble C3b with a reactive thioester. This organization makes clear the following analogy: Consider [Sites] to be like cells in an organism; [C3b_soluble_] is like a reservoir of active virus in the extracellular space; and [C3_bound_] is like a virus that has infected a cell, and is thus able to create more viruses (C3bs) that can infect more cells (Sites). With this analogy in mind, we compared our minimal subnetwork (leftmost equation in Fig 7b) to the well-studied TIV model of viral infection (rightmost equation in Fig 7b). Noting that the tickover rate (k_tick_) is orders of magnitude slower than the rate of C3 cleavage produced by the amplification loop, we can eliminate the term k_tick_[Sites] in the leftmost equation. That modification shows that our minimal subnetwork and the TIV model are equivalent in the following sense: They identical forms of their ODEs, only changing the names of the variables and parameters (see the color coding in Fig 7b). Thus, our minimal subnetwork should have the same dynamical properties as the classical TIV model.

**Figure 7.**
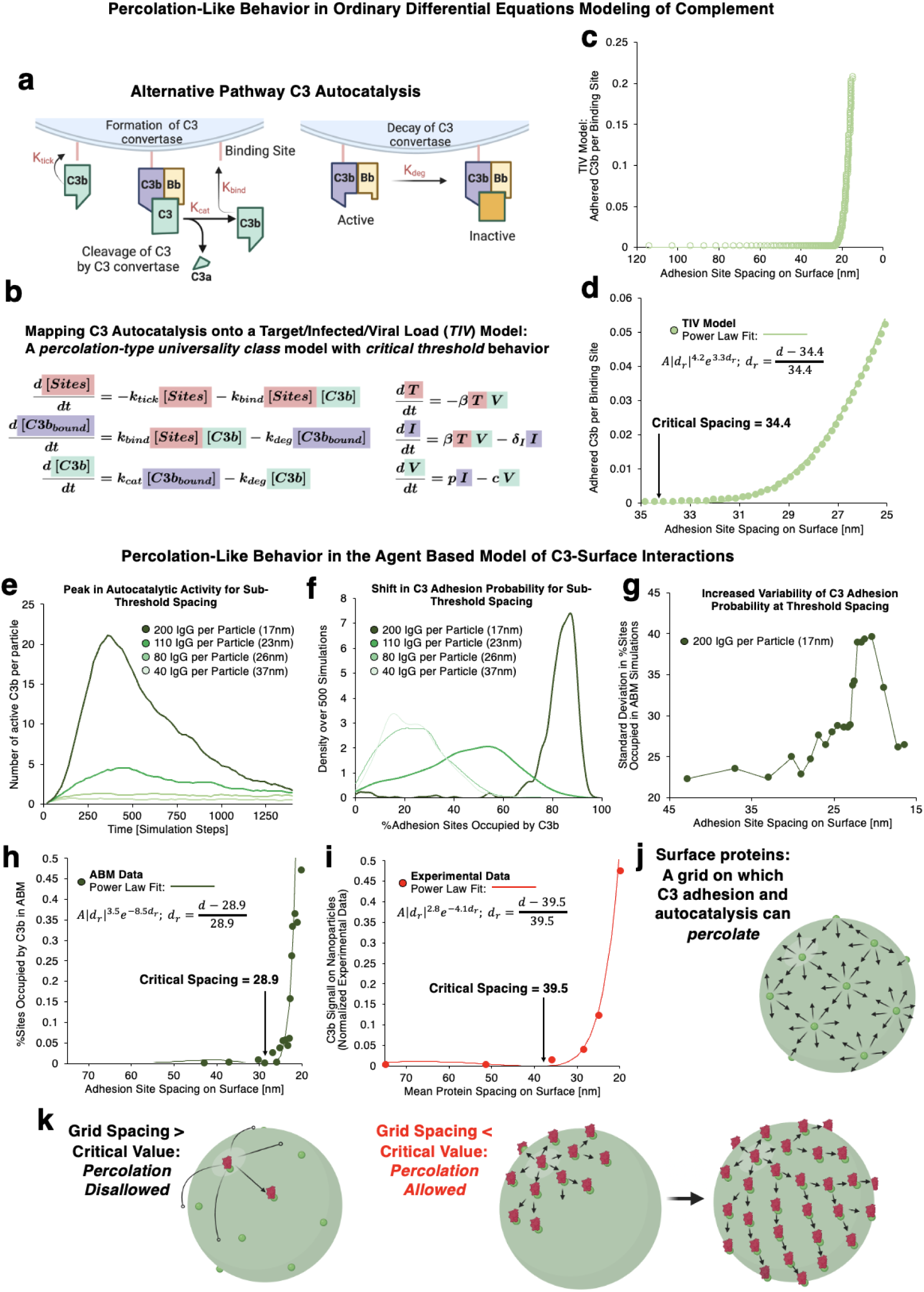
Modeling data show that the percolation universality class can explain the reaction of complement to surfaces. **a,** Simplified schematic of alternative pathway C3-surface adhesion and autocatalysis, with reaction constants highlighted in red text. **b,** Rate equations modeling the reactions depicted in **a**, showing that the system of differential equations describing the simplified model of C3-surface adhesion and autocatalysis matches the system of differential equations describing the TIV model for infection spreading, a percolation universality class phenomenon. **c,** Results showing that the TIV model recapitulates the threshold effect in C3-surface interactions. **d,** Magnified results from **c**, showing that TIV model findings near the threshold surface protein spacing fit a power law relationship, characteristic of the percolation universality class. **e-h,** Results from agent based model simulations (Fig. 6c-e) showing characteristics of the percolation universality class and critical threshold behavior. Agent based model results show; **e**, a peak in C3 autocatalytic activity only for sub-threshold surface protein spacing; **f,** a precipitous shift in C3-surface interaction probability for sub-threshold surface protein spacing; **g,** an increase in variability of C3-surface interaction probability around the threshold spacing; **h,** a close fit to a power law curve that matches the description of a percolation universality class phenomenon. **i,** Experimental measurements of C3b adhesion to nanoparticle surfaces (reproduced from Fig. 2a), showing a fit to a power law curve, as in **d** and **h**. **j**, Schematic describing a protein-coated surface as a grid for “complement percolation.” **k,** Schematics depicting models for how “complement percolation” behaves in response to above and below threshold surface protein spacing.

Thus, to more deeply understand complement’s minimal subnetwork, we can investigate the class of dynamical models into which TIV falls. Upon a quasi-steady-state assumption (QSSA), the TIV (and thus our minimal subnetwork) is equivalent to the more famous SIR (susceptible-infected-recovered) model of disease spread in a population^26^. Since we empirically showed that a QSSA holds for our minimal subnetwork (Supplementary Computational Model, section S3.3), we can therefore look to the well-studied dynamics of SIR to understand complement’s minimal subnetwork. The SIR model has been shown to display critical phenomena, including phase transitions (sharp transitions in a macroscale order parameter) upon varying control parameters like the distance between susceptible individuals^27–30^. Models displaying critical phenomena fall into particular universality classes, within which all the models display similar dynamics near their critical points, defined by critical exponents. SIR falls into the percolation universality class^30,31^, originally defined by the percolation model of hydrology: When a fluid contacts the surface of a porous rock, the probability of the fluid flowing to the other side of the rock is a function of the probability of connection between two adjacent pores, and as that latter probability increases, there is a phase transition in the probability of fluid flowing from one side to another. Other models falling into the percolation universality class include the forest-fire model, which also conceptually maps to our minimal network: Potential C3 attachment sites are like trees, and C3b is like fire. Thus, this analysis suggests that the minimal complement subnetwork (and the entire complement network) should display properties previously shown for critical points in the percolation universality class.

To see if the minimal subnetwork of complement actually has the properties of percolation, we analyzed results from the TIV-like model from Fig 7b (leftmost equations, using parameters created by lumping the parameters from the full 107-equation ODE model). The TIV-like model has a sharp threshold, as in our prior modeling and experimentation (Fig. 7c). Importantly, for models in the percolation universality class, near the critical point, we should observe a power-law fit for the number of C3b adhered per binding site, which we do (Fig 7d).

Because percolation is most studied within lattice networks, we next examined whether our ABM, which employs a lattice, also has percolation dynamics. Investigating the temporal dynamics of the ABM, we find that nanoparticles with below-threshold surface protein spacing have a transient peak in the number of active C3b-surface adducts, while above-threshold nanoparticles have no such transient peaks (Fig 7e). Such transient spikes only occurring above the critical point have been observed in other percolation models, such as for forest fires, where below the critical point of distance between trees, fire spreads rapidly but the number of engulfed trees peaks rapidly as trees are completely consumed.

Next, as is common in percolation theory, we examined the probability density function for the extent of coverage of lattice sites. In Figure 7f, we show that for nanoparticles with large distances between surface proteins, we have a right-tailed distribution, but that abruptly changes to a left-tailed distribution above the critical threshold. For above-threshold nanoparticles (e.g., 37 nm between attachment sites) there is a near-zero probability of a nanoparticle getting nearly totally coated by C3b, but for below-threshold nanoparticles (e.g., 17 nm distance), there is a >90% chance that >75% of the lattice sites (surface proteins) are conjugated to a C3b. This sharp transition is analogous to the phase change in “percolation probability” that is observed in the original percolation lattice model, and here corresponds to the probability of runaway events we posited based on experimental data (Fig 2j).

We investigated the ABM for three other hallmarks of critical points. As shown in Figure 7g (and more extensively in Supplemental Computational Figure SC7) and as expected of critical points, the variability in C3b deposition (standard deviation in percentage of C3b-occupied binding sites) peaks at the critical point. Further, as the physical size of the system increases, the standard deviation in the maximum number of active C3b-adducts also increases (Supplemental Computational Figure SC8). Finally, we checked the ABM for the classic fit of a power-law distribution near the critical point. We found such a power-law fit (Fig 7h) and showed a similar fit applies to our experimental data (7i).

To summarize how percolation explains our experimental results (Fig 7j & k): The complex web of complement protein reactions has a small subnetwork of reactions that have percolation-type criticality. The proteins on a surface function as points on a lattice/grid, and, via autocatalysis, C3b attachment to these points can spread to nearby lattice points in a process mapping to the mathematical description of fluid percolating through pores in a rock (Fig 7j). For grid spacing greater than the critical point, complete percolation is disallowed (no C3-spreading runaway events) (Fig 7k, left). But for grid spacing less than the critical value, percolation events can occur (C3 spreads across the surface in a runaway event; Fig 7k middle & right), producing a “sharp threshold” of complement activation at the population level, and subpopulations of particles that are completely coated with C3b.

## Discussion

This study illuminates the surprising and unintuitive ways in which biological complex systems interact with materials. While it is difficult enough to understand how complicated tangles of biological signaling networks make “decisions” on their own, it is even more complicated to elucidate such questions when the network is interacting with a complicated foreign material. To explore this under-studied area, we focused on the clinically important complement signaling network, which attacks most advanced therapeutics (nanoparticles, viral gene therapy, etc.) and attacks our own tissues in most inflammatory diseases. Understanding the principles upon which complement “decides” to attack a particular surface can thus guide our design of advanced therapeutics and therapies for inflammatory diseases. Decades of molecular biology studies have identified simple material features necessary for complement attack, such as an abundance of surface nucleophiles and an absence of recruited C3 inhibitors such as Factor I. However, there have been too few studies mechanistically probing the nanoscale features of a surface that are necessary for complement attack^15^, and no studies that have brought together nano-scale engineering with the tools of complexity science, which are necessary to understand complement’s “decisions” at their deepest levels.

We showed here that the complex system of complement proteins “decides” whether to attack a material’s surface by sensitively distinguishing if the surface density of potential complement attachment points exceeds a threshold. We showed that this switch-like sharp threshold behavior has manifestations that range over seven orders of magnitude on the length scale, from the nanoscale (C3b deposition on individual nanoparticles [Figs 2 and 5]) to the macroscale (pathologies in animals [Fig 1]). We found this switch-like behavior in complement interactions with very diverse surfaces, ranging from the fluid surfaces of clinically relevant liposomes (Fig 2, 3), to rigid nanoscale “sheets” whose proteins are fixed in place with single-digit-nanometer precision (Fig 5), and even to bacteria (Fig 4).

While investigating this threshold behavior, we serendipitously observed a second important phenomenon: Complement-coating of *individual* nanoparticles involves stochastic “runaway” events where some nanoparticles get completely coated with complement (Fig 2). The threshold behavior was first noted for *population-level* measurements of nanoparticles: A population of nanoparticles with below-threshold surface protein spacing heavily activates complement (measured by total molecules of C3b and properdin deposited, and C5a produced), while a population of nanoparticles with above-threshold spacing does not activate complement significantly. When we look at individual particles in a population of below-threshold nanoparticles, we find only a fraction of the nanoparticles is completely C3b-coated, while many individual particles have much lower levels of C3b. This second phenomenon of runaway C3b deposition on nanoparticles was only detectable because of recent innovations in single nanoparticle measurements, here using flow cytometry with nanoscale particle detection and fluorescent nanoparticle tracking analysis.

Having found the dual phenomena of switch-like behavior at a population level and runaway coating of individual nanoparticles, we had to understand how the complement cascade of protein interactions was producing these phenomena. At first glance, it may seem impossible to understand how these elegant phenomena arise out of complement’s complicated wiring diagram of ∼40 proteins, with their numerous and overlapping interconnections of chemical reactions (both enzymatic and binding reactions). To understand this complicated web’s decision-making, we took two steps: First, we recapitulated the two phenomena with computational models (Fig 6), and, second, we used the tools of complexity science to understand the network interactions within the computational models (Fig 7).

As all biological systems are dynamical systems, we employed the two most common frameworks to model dynamical systems: ODEs and ABMs. The coupled ODE model had the advantage of being able to use reaction rate constants and concentrations that were previously measured experimentally or estimated using prior modeling. However, the ODE system used here did not model individual nanoparticles, but rather implicitly modeled a single, infinite surface. By contrast, the ABM explicitly models individual nanoparticles on a quasi-2D surface, thus creating the type of lattice network that has worked well for understanding critical phenomena in statistical physics. Both models recapitulated our major observed experimental phenomena, doing so without parameter fitting. Further, they produced a critical threshold of surface protein spacing of ∼25 nm, which is what we found experimentally. These and several smaller results show that the complement cascade’s surprising behaviors are all derivable from the previously known network architecture.

But to truly understand complement’s behaviors and not simply recapitulate them with high-dimensional models, we needed to more fully apply the tools of complexity science. We were motivated by the fact that our observed phenomenology (sharp thresholds, runaway coating) was reminiscent of phase change behavior seen in statistical mechanics, the study of how large ensembles of microscopic objects produce macroscopic behavior. While phase changes are best known to the lay public in the form of the solid-liquid-gas transitions of states of matter, in statistical mechanics, phase transitions are generalized to occur when there is a singularity in the free energy or one of its derivatives^31^. This is mathematically generalized further in complexity science to include singularities occurring as complex systems’ control parameters are varied, producing an abrupt change in an order parameter (e.g., in a rock, changing the probability of connections between pores [a control parameter] can produce an abrupt change in the probability of a continuous path for fluid to flow through the pores from one side of the rock to the other [order parameter]). These generalized phase changes seemed to fit the behavior of complement, but to demonstrate this, we first sought to understand what was the minimal subnetwork of complement that could recapitulate the observed phenomena, as a smaller network might lend itself better to the analysis of phase changes.

Fortunately, we found a minimal subnetwork of complement that recapitulates the key phenomenology. By grouping reactions, the 107-equation ODE model was reduced to a 3-ODE model. This minimal subnetwork recapitulated the sharp threshold effect (Fig 7c). We then showed that this minimal subnetwork’s ODE model is equivalent to the famous SIR model from epidemiology. SIR is in the percolation universality class, and thus so is our subnetwork which captures threshold behavior of the larger network of complement. We went on to show that our minimal complement subnetwork’s critical point (the mathematical term for its “sharp transition”) has a key characteristic of a percolation-type critical point, power-law scaling near the critical point (Fig 7d). Further, our ABM of complement has numerous hallmarks of percolation near its critical point, including divergence of the variance (Fig 7e-h). Importantly, we would not have sought out these additional properties of the complement system without knowing that the complement cascade fits within the percolation universality class. Since percolation has been studied for decades in much simpler systems, those results can now guide further exploration of how complement and other signaling pathways work. Thus, we now have one deep answer to the original question of how complement “decides” which surfaces to attack: By using a percolation-type critical point.

This answer creates many new questions. First, the path to finding this critical point suggested a phenomenon that we have not yet directly observed: “C3 spreading” (Figs 2j and 6c). This hypothesized mechanism suggests that once one C3b deposits on a surface, a wave of C3b-adducts will spread out from it, created by the alternative pathway’s amplification loop. The experimental observation of runaway C3b coating of nanoparticles suggests C3 spreading must occur, as does our observation of C3 spreading in ABM simulations (Supplementary Movies 13-15). However, we need to develop new experimental tools to directly observe if C3 spreading waves actually occur. Second, we must understand the development of complement’s critical point in the context of evolution and human variation. Complement evolved 500 million years ago, but key components vary significantly between species, and even between individual humans. Do these variations affect criticality? What are the pathophysiological consequences of the runaway C3b coating on only a fraction of nanoparticles, rather than uniform coating? Lastly, it is important to expand the use of complexity science beyond the study of percolation. For example, we previously reported^19^ that among nanoparticles with surface proteins, a near-crystalline (low entropy) organization of the proteins (as in viral capsids or ferritin) produces far less complement activation than an amorphous (high entropy) supramolecular organization (as in nanoparticles formed by hydrophobic protein-protein interactions or in many types of protein coatings on liposomes). This suggests that configurational entropy, a concept from statistical mechanics, might be another surface property with a role in C3 spreading.

To answer these and other questions, we must continue the approach of combining the experimental tools of nano-engineering with the computational tools of complexity science. Here, that approach provided a new way to understand the interactions between complex systems and materials, finding elegance hiding within complexity.

## Methods

### Liposome preparation

Liposomes were prepared with azide-functionalized lipids to enable controlled bio-orthogonal conjugation to alkyne-functionalized proteins. Techniques for liposome preparation were described and validated in previous work^32^. Lipids were prepared with 54 mol% DPPC (Avanti), 40 mol% cholesterol (Avanti), and 6 mol% azide-PEG_2000_-DSPE (Avanti). For experiments requiring fluorescent liposomes, including flow cytometry and microscopy work, 0.5 mol% Top Fluor PC (1-palmitoyl-2-(dipyrrometheneboron difluoride) undecanoyl-sn-glycero-3-phosphocholine) was added to the lipid mixtures. To prepare lipid films from the mixtures, lipids in chloroform were dried with nitrogen or argon gas, then lyophilized overnight to remove residual solvent. Dulbecco’s phosphate buffered saline (PBS) was added to the dried lipids at a 20 mM final lipid concentration. The resultant lipid dispersions were frozen with liquid N_2_ and thawed in a 50°C water bath three times. After the three freeze-thaw cycles, the lipid suspensions were extruded 10 times through track-etched polycarbonate filters with 200 nm rated pore size. Post-extrusion, size of the liposomes was assessed with dynamic light scattering (DLS, Malvern) and numerical concentration was determined by nanoparticle tracking analysis (NTA, Nanosight/Malvern). Zeta potential was determined by zetasizer analysis (Malvern). Liposome sizes, zeta potentials, and concentrations were also assessed after each liposome modification (*e.g.*, conjugations) described below.

### FAb preparation

FAb fragments were prepared from rat IgG with Pierce’s FAb preparation kit (Thermo Scientific). Briefly, Papain-loaded agarose resin was seeded in a spin column, IgG was exchanged into digestion buffer via Zeba spin desalting column, and the resulting IgG solution was added to the prepared Papain column. The IgG-loaded Papain column was incubated in accordance with manufacturer recommendations for incubation time as a function of loaded IgG concentration. The cleaved IgG was removed from the Papain column via centrifugation, then buffer exchanged into PBS. FAb fragments were separated from undigested IgG and Fc fragments using a protein G spin column. FAb concentration was determined by absorbance at 280 nm, with an estimated extinction coefficient of 1.4.

### Protein iodination

Protein (IgG or FAb, as described above) was prepared in PBS at concentrations between 1 and 2 mg/mL in volumes between 100 and 200 µL. To oxidize radiotracer iodide salt for labeling of the protein, one Iodobead (Perkin-Elmer) was added to a borosilicate reaction tube as an oxidizing surface. Protein solution was added to the tube alongside the Iodobead, followed by Na^125^I at 25 µCi per 100 µg of protein. The protein-iodine reaction was allowed to proceed for 5 minutes at room temperature. To shield against exothermic splattering with the reactive radiotracer iodine, the reaction vessel was covered with parafilm and kept in a ventilated hood. After 5 minutes, the radiolabeled protein was separated from the reactive iodine by passing through a 7 kDa cutoff centrifugal desalting column (Zeba). Additional passages through fresh desalting columns removed remaining free iodine. Radioactivity in the protein sample vs. radioactivity retained in the desalting columns quantified protein-bound vs. free iodine in the samples. Labeled protein was only used if >95% of radioactivity was associated with protein.

### Fluorophore labeling of proteins

Anti-C3b, anti-properdin, and recombinant C3 were labeled with NHS ester Alexa Fluor 488 or NHS ester Alexa Fluor 647 for flow cytometry and fluorescence-mode nanoparticle tracking analysis experiments. To label the proteins, NHS ester fluorophore (ThermoFisher) was added to protein in PBS at 5:1 mol fluorophore:mol protein ratio. The mixture was incubated for two hours on ice and unreacted fluorophore was removed by centrifugation against 10 kDa molecular weight cutoff filters (Amicon) in four centrifugation and rinsing cycles. After recovery of labeled protein, spectrophotometry (Nanodrop) measured optical density of the resultant solutions at 280 nm and either 490 nm or 647 nm. C3 or antibody concentration was determined by optical density at 280 nm and fluorophore concentration was determined by optical density at 490 nm or 647 nm, with extinction coefficients of 73,000 L*mol^−1^cm^−1^ and 260,000 L*mol^−1^cm^−1^ for Alexa Fluor 488 and Alexa Fluor 647, respectively. Density of fluorophore on proteins was determined by the ratio of the two concentrations and protein was used if the labeling achieved >2 fluorophores per protein.

### Liposome-protein conjugation

Rat IgG or FAb, prepared as above, was functionalized by linking to dibenzylcyclooctyne-PEG_4_-NHS ester (DBCO, Jena Bioscience). IgG solutions (PBS) were adjusted to pH 8.3 with 1 M NaHCO3 buffer and reacted with DBCO for 1 hour at room temperature at a molar ratio of 20:1 DBCO:IgG. Excess DBCO was separated from DBCO-protein conjugates with 10 kDa cutoff centrifugal filters (Amicon). DBCO-IgG reaction efficiency was determined by spectrophotometry. Optical density at 280nm was proportional to IgG concentration and optical density at 309 nm determined DBCO concentration. Overlap between the DBCO and IgG spectra was corrected for by adjusting the 280 nm value with *Abs_280,Corrected_ = Abs*_280_ − 1.089 x *Abs*_309_. IgG concentration was determined via 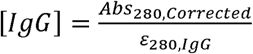, where *ε*_280,*IgG*_ is 204,000 L mol^−1^ cm^−1^, the IgG extinction coefficient at 280 nm. Molar DBCO concentration was determined via 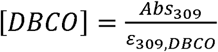, where *ε_280,DBCO_* is 12,000 L mol^−1^ cm^−1^, the DBCO extinction coefficient at 309 nm. Extent of DBCO-IgG coupling was determined as 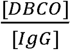. An identical analysis was used for FAb modification, but with *ε_280,FAB_* equal to 70,000 L mol^−1^ cm^−1^.

Protein loading on liposomes was planned based on liposome concentration, as determined by nanoparticle tracking analysis and DBCO-modified protein concentration, as determined above. At different protein:liposome ratios, DBCO-modified IgG or FAb was added to liposomes including azide-functionalized lipids. Liposome/protein mixtures were incubated overnight at room temperature. Protein was separated from protein-liposomes via Sepharose size exclusion chromatography. Isolated protein-liposomes were then concentrated against centrifugal filters (Amicon, 10 kDa cutoff). After recovery from the filters, protein-liposome concentrations were determined by nanoparticle tracking analysis.

For measurement of protein-liposome coupling efficiency or tracing of liposomes in biodistribution measurements (as described below), tracer amounts of radiolabeled protein (1:10 radiolabeled:non-labeled molar ratio) were added to protein-liposomes reaction. After protein-liposome conjugation reactions, when protein + protein-liposome mixtures were passed through Sepharose columns, 0.5 mL fractions were collected and ^125^I radioactivity was measured in each fraction. Radioactivity in fractions corresponding to free protein, *p*, was compared to radioactivity in fractions corresponding to liposomes, *l*, to determine the efficiency of protein-liposome coupling 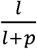. Comparing this efficiency to the quantity of protein added per liposome gives the yield number of proteins per liposome. The protein-liposome elution fractions, collected and concentrated as above, were reserved for *in vivo* radiotracing experiments to determine protein-liposome biodistributions.

### Biodistribution measurements

For this study, all biodistribution data were obtained in mice injured by intravenous bacterial lipopolysaccharides (LPS, from B4 strain *E. coli*) in advance of nanoparticle administration. To achieve this injury, mice were anesthetized with 3% isoflurane gas, then LPS in 100 µL PBS was injected at a dose of 2 mgkg^−1^ retro-orbitally. Liposomes were administered 5 h after LPS.

IgG-liposomes with tracer quantities of ^125^I-IgG were used in biodistribution experiments. Liposomes in PBS were injected intravenously into C57BL/6 male mice from Jackson Laboraties. The liposome dose was set at 2 mgkg^−1^ based on lipid+protein concentration in the liposome formulation) and a quantity of liposomes appropriate to that dose was brought to a final volume of 100 µL for bolus injection in the jugular vein, while mice were anesthetized with ketamine/xylazine (10 mgkg^−1^ ketamine, 100 mgkg^−1^ xylazine, intramuscular administration). Radioactivity in the injected dose was measured via gamma counter (Perkin-Elmer) immediately before injection.

Mice were sacrificed by vena cava blood draw into EDTA-coated syringe at 30 min after liposome injection. Drawn blood was analyzed with a complete blood count analyzer (Abaxis). Heart, lungs, liver, spleen, and kidneys were removed and rinsed in saline. Quantity of liposomes in each organ, along with a 100 µL blood sample, was determined by ^125^I radioactivity measured via gamma counter. Radioactivity in organs was compared to the radioactivity in the injected liposome dose to determine the fraction of dose in each organ and liposome concentration in each organ was calculated by normalizing the retained fraction of dose to the organ weight.

Blood samples from biodistribution experiments were also used to quantify hemolysis induced by liposomes over 30 min circulation in mice. Blood was centrifuged at 1500xg for 10 min and plasma was isolated from red blood cell and leukocyte layers. Spectra of optical density of plasma between 350 nm and 750 nm were obtained on a plate reader spectrophotometer. Peaks in optical density at 415 nm and 540 nm were used as markers of hemoglobin concentration in the plasma, providing a metric for red blood cell lysis in the blood samples.

### Flow cytometry tracing of liposome uptake in lungs

Single-cell suspensions were prepared from male C57BL/6 mouse lungs for flow cytometry. Fluorescent liposomes were administered at 2.5 mgkg^−1^ 30Dminutes before the animals were killed and their lungs extracted. Mice were anaesthetized with ketamine/xylazine (10 mgkg^−1^ ketamine, 100 mgkg^−1^ xylazine, intramuscular administration), killed by vena cava exsanguination, and lungs were perfused via the right ventricle with ∼10Dml of cold PBS. Lungs were excised and triturated on ice. The disaggregated lung tissue was then incubated in 3 mL of a digestive solution containing 2.5 mgmL^−1^ collagenase I and 1 mgmL^−1^ DNAse I for 45 minutes at 37°C, vortexing every 10 minutes. The digested lung tissue was passed through a 70 µm cell strainer and washed with PBS. The strained material was centrifuged at 500xg for 5 minutes at 4°C. The supernatant was removed and the pellet was resuspended in 10 mL cold ACK lysis buffer. After 5 minutes incubation on ice to lyse red blood cells and a second passage through a 70 µm strainer, the remaining cells were centrifuged again at 500xg for 5 minutes at 4°C. The supernatant was discarded and the cells were resuspended in PBS. Cell concentration was determined by hemocytometer and cells were diluted to achieve a concentration of 1×10^6^ cells/mL.

Cells were pelleted at 500xg, washed with FACS buffer (2% fetal calf serum and 1 mM EDTA in PBS), and incubated with Fc block for 15 minutes. The washed and blocked cells were incubated with staining antibodies for 30 minutes in the dark at room temperature. Excess antibody was removed by centrifugation at 500xg and washing with FACS buffer. The stained cells were then fixed in 4% PFA in a 10 minute room temperature incubation, followed by pelleting at 500xg and resuspension in FACS buffer prior to flow cytometry analysis. Gating strategies for analysis of flow cytometry data are depicted in Supplementary Figure 7. For leukocytes, cells were stained with anti-CD45 (Brilliant Violet 395), anti-Ly6G (Alexa Fluor 750), and anti-CD64 (PerCP Cy5). For non-leukocytes, cells were stained with anti-CD31 (Alexa Fluor 647) and anti-EpCAM (Brilliant Violet 711). Single-stain controls allowed automatic generation of compensation matrices in FlowJo software.

### ELISA methods

C3a and C5a concentrations were assessed by enzyme-linked immunosorbent assay (ELISA). For measurement of C3a and C5a generation *in vitro*, 20 µL fresh mouse serum was incubated with 20 µL protein-liposomes (numerical liposome concentration set to 2×10^12^ liposomes/mL) for 60 minutes. EDTA was added to the serum-nanoparticle mixture at 60 minutes to inhibit further complement activation. To inhibit the classical and lectin pathways of complement and isolate the alternative pathway response to liposomes, 10 µL of gelatin veronal buffer (GVB) was added to serum before adding 10 µL of liposomes (concentration set to 4×10^12^ liposomes/mL). In experiments where properdin was inhibited prior to liposome-serum incubation, 200 µL of serum was treated with 0.5, 1, 5, 10, or 50 µg of properdin-inhibiting antibody (prepared by Wenchao Song’s lab at the University of Pennsylvania) for 15 minutes. The above experimental protocol with 20 µL of serum and 20 µL of liposomes (2×10^12^ liposomes/mL) was repeated with the anti-properdin-treated serum. For measurement of C3a levels induced by liposomes *in vivo*, liposomes were administered to C57BL/6 mice at a dose of 3 mg/kg as an intravenous bolus (as in biodistribution experiments described above) and blood was drawn from the vena cava into EDTA at different time points after liposome injection. Plasma was separated from red blood cells and leukocytes by 1500xg centrifugation for 10 minutes. For all *in vitro* and *in vivo* conditions, after EDTA termination of the serum/plasma-liposome incubations, C3a or C5a levels were measured by using sandwich ELISA kits from BD Biosciences.

C5b-9 concentration was also determined by ELISA in *in vitro* experiments. Blood was collected from C57BL/6 mice via vena cava draw into heparin-coated tubes. 100 µL heparinized whole blood was incubated with 20 µL liposomes (2×10^12^ liposomes/mL) for 60 minutes, then EDTA was added to the mixture to inhibit further complement activation. Plasma was isolated from the whole blood by 2500xg centrifugation for 20 minutes, in accordance with methods recommended for the C5b-9 ELISA kit (AFG Biosciences). Capture, washing, and detection procedures were carried out as prescribed by the kit instructions.

### Flow cytometry measurements of complement adhesion to nanoparticles

20 µL fresh serum was incubated with 20 µL liposomes formulated with TopFluor PC (2×10^12^ liposomes/mL) for 60 minutes. EDTA was added to the mixture at 60 minutes incubation time to inhibit further complement-liposome interactions. Following ELISA protocols described above, GVB treatment of serum was used to isolate alternative pathway complement responses to liposomes and properdin-inhibiting antibody treatment of serum was used to limit properdin contributions to the complement response. Serum-nanoparticle mixtures were treated with fluorophore-labeled anti-c3b or anti-properdin antibodies (see “fluorophore labeling of proteins” above) for 30 minutes. The serum-nanoparticle-antibody mixtures were diluted 1:10000 before flow cytometry analysis. Flow cytometry data was obtained with BD FACS Canto or FACS Symphony flow cytometers, with the FACS Canto being adjusted with custom modifications designed to improve detection of extracellular vesicles. Specifically, detection threshold was determined based on side scatter with a 200 mW 488 nm laser and collected light was passed through a pinhole detector to reduce noise and enhance signal collected from the center stream. Gating strategies for single particle flow cytometry are depicted in Supplementary Figure 17.

### Nanoparticle tracking analysis measurements of complement adhesion to nanoparticles

Recombinant C3 (Complement Technologies) was labeled with Alexa Fluor 488 as described above. 30 µg fluorescent C3 was added to 20 µL of serum in 20 µL of PBS, then 4×10^10^ liposomes were added at a concentration set to achieve a final reaction volume of 60 µL. For experiments designed to isolate the alternative pathway of complement, serum was combined with 20 µL of GVB prior to addition of fluorescent C3 and liposomes. After 45 minutes incubation at room temperature, the 60 µL reaction volume was diluted with 940 µL of deionized and filtered water (MilliQ) and 100 µL of the diluted suspension was further diluted in 900 µL of water to prepare the dilute suspension analyzed with NTA. NTA analysis was completed in both fluorescent mode (488 nm laser with a 500 nm long pass filter) and scattering mode. Fluorescence detection identified nanoparticles with Alexa Fluor 488 labeled C3 on their surfaces. As controls, serum with fluorescent C3 and no nanoparticles or nanoparticles with no serum or fluorescent C3 were tested with identical reaction timing/volume and identical dilution to confirm specificity of the fluorescent C3 signal to C3-nanoparticle interactions.

### Microscopy characterization of complement adhesion to surface-bound nanoparticles

Glass cover slips were washed with deionized and filtered water (MilliQ) then ethanol, then dried under filtered nitrogen gas. Prepared cover slips were placed in a sealed glass container with an open container of 5 mL (3-aminopropyl)triethoxysilane (APTES) overnight at 70°C to permit vapor deposition of amine-terminated silanes. The amine-functionalized cover slips were rinsed in water for 30 minutes, then 100 µL of 10 µM NHS-PEG_4_-azide was added to the cover slips overnight at room temperature to functionalize the cover slips with azide-terminated moieties. Excess NHS-PEG_4_-azide was removed by rinsing with deionized and filtered water. The azide termini were used to immobilize fluorescent liposomes with azide-reactive DBCO-IgG on their surfaces. Liposomes were incubated with the azide-functionalized cover slips overnight at room temperature, then excess liposomes were removed by rinsing with deionized and filtered water. Cover slips with immobilized liposomes were incubated with 100 µL of PBS-or GVB-treated serum for 60 minutes at room temperature, then rinsed with PBS. Finally, the cover slips were stained with anti-C3b antibody labeled with Alexa Fluor 647 for one hour at room temperature, followed by rinsing with water. The cover slips were imaged with confocal microscopy to detect TopFluor PC fluorescence from liposomes and Alexa Fluor 647 fluorescence from anti-C3b staining. Co-registered images were recorded and the ratio of mean C3b fluorescence to mean liposome fluorescence was plotted.

### Preparation and characterization of E. coli

TOP10 E. coli (ThermoFisher #C404002) were grown in Terrific broth with ampicillin and an aliquot was heat-inactivated by 40□minute incubation at 60□°C. Bacteria were pelleted by centrifugation at 1000xg for 10 minutes, washed three times in PBS, then resuspended by pipetting to an estimated concentration of 2×10^8^ colony forming units (CFU)/mL. Bacterial concentration was verified by optical density at 600 nm, where linear correlation between optical density at 600 nm and bacteria concentration was confirmed by serial dilution of bacteria and an optical density of 1 arbitrary unit was determined to correspond to 8×10^8^ CFU/mL. To 50 µL of suspended bacteria, we added 0.0175 µmol, 0.05 µmol, 0.5 µmol, or 5 µmol of methyl-PEG_4_-NHS ester or a PBS sham at equivalent volume for overnight incubation while rotating at 4°C. Excess methyl-PEG_4_-NHS ester was removed by centrifuging the bacteria at 1000xg for 10 minutes then rinsing with PBS three times. Concentrations of bacteria after modification were confirmed by measurement of optical density at 600 nm.

Concentration of nucleophiles in E. coli suspensions was determined by modified ninhydrin assay. 0.16 g ninhydrin and 0.024 g hydrindantin were dissolved in 6 mL dimethyl sulfoxide. Lithium acetate was prepared at 4 M concentration in water and mixed with the ninhydrin solution at 1:3 vol:vol. 2,2’-(ethylenedioxy)bis(ethylamine) (EOD) standards were prepared at concentrations between 0 and 5 mM in 1:1 volume:volume ethanol:water. 50 µL of EOD standard solutions or E. coli suspensions (with and without methyl-PEG_4_-NHS treatment) and 50 µL of ninhydrin/hydrindantin/lithium acetate stock were mixed and heated at 40°C for 10 minutes. 100 µL of ethanol/water diluent were added at the end of the reaction and optical absorbance spectra were obtained for the resulting solutions, noting intensity of the Ruhemann’s Purple absorbance peak at 570 nm produced by reaction of ninhydrin with nucleophiles. Linearity of 570 nm optical density with respect to concentration of EOD was confirmed (Supplementary Figure 31) and unknown concentration of nucleophiles in E. coli samples was determined by comparison to abs(570) vs. EOD concentration curves. Average protein spacing on E. coli was estimated based on nucleophile concentration and the theoretical E. coli surface area. Details of the protein spacing calculation are included in “*Summary of determination of protein-protein spacing on liposomes and bacteria*” below.

20 µL of E. coli suspensions with different nucleophile densities were incubated with 20 µL of serum for 60 minutes at room temperature for C3a ELISA measurements of complement responses to the bacteria, following procedures detailed in “*ELISA methods*” above. Similarly, after incubation with serum, E. coli were stained with fluorescent anti-C3b or anti-properdin antibodies to detect C3b or properdin on E. coli surfaces in flow cytometry data, as described for liposomes in “*Flow cytometry measurements of complement adhesion to nanoparticles*” above. Methods in “*Nanoparticle tracking analysis measurements of complement adhesion to nanoparticles*” above were replicated in incubations of E. coli with serum doped with fluorophore-labeled C3. Instead of nanoparticle tracking analysis, to characterize fluorescent C3 adhesion to E. coli, widefield microscopy was used to image colocalization of E. coli with fluorescent C3. E. coli/serum/fluorescent C3 suspensions were pipetted onto glass slides. The slides were allowed to air dry, then the bacteria were fixed to the slide surfaces by briefly exposing the underside of the slides to flame. The slides were rinsed under running deionized and filtered water (MilliQ), dried under air, then imaged.

### Preparation of DNA origami

Rectangular DNA origami tiles were prepared in 1 × TAE-Mg^2+^ buffer using established protocols^33^. For each sample, 20 nM single-stranded M13mp18 DNA (7,249 nucleotides) was mixed with a 5-fold molar excess of staple stands and a 10-fold molar excess of probe strands. The mixture was annealed from 95 °C to 4 °C (see Supplementary Section D for detailed annealing program). The excess staple strands were removed by 3x repeated washing in 1×TBS Mg^2+^ buffer (pH 7.5) and filtered using 100 kDa cutoff 500 µL Amicon filters. The concentration of DNA origami tiles was quantified by absorbance at 260 nm, assuming an extinction coefficient of ∼109,119,009 M^−1^cm^−1^. Staple strand sequences were designed to allow IgG positioning at different locations on the origami. Detailed sequences, including positioning staples, are included in Supplementary Section D, tables 1-4. Origami structures were confirmed with atomic force microscopy.

### IgG-DNA conjugation

IgG was functionalized with DBCO as described above. IgG-DBCO was conjugated to azide-terminated strands (5-Azide-TTTCACACACACACACACACA-3) at a 1:2 mol:mol ratio overnight at 4 °C. Excess strands were removed by passing the solution through 10 kD-cutoff Amicon filters. DNA-conjugated IgG were then mixed with DNA origami tiles in 1 × TBS-Mg^2+^ buffer (pH 7.5) at a molar ratio of 2:1. The solution mixture was cooled from 37°C to 4°C (see Supplementary Section E for detailed annealing program). IgG-DNA origami were characterized with AFM and used in C3a ELISAs, with origami concentration set at 20 nM in 30 minute serum incubations at room temperature and other aspects of C3a detection conducted as described above. Origami were also immobilized on glass slides fitted for 96 well plates (Electron Microscopy Sciences, Inc.), incubated with serum, then stained with fluorescent anti-C3b antibody. 20 nM origami suspensions incubated with glass slides for 1 hour at room temperature, then slides were incubated with serum for 30 minutes at room temperature and rinsed with PBS three times. Fluorescent anti-c3b antibody (5 µL antibody stock diluted in 100 µL PBS) was added to the serum-treated origami-slides for 1 hour incubation at room temperature, followed by threefold rinsing with PBS. Anti-C3b fluorescence stain on glass slides was quantified by plate reader.

### Summary of determination of protein-protein spacing on liposomes and bacteria

For liposomes, we defined an average spacing of proteins on the nanoparticle surfaces. We measured the diameter of the liposomes before adding protein to the nanoparticles, using dynamic light scattering and nanoparticle tracking analysis. We used that diameter to determine the amount of liposome surface area on which the proteins will be grafted. As part of the nanoparticle tracking analysis, we also counted the numerical concentration of the liposomes to which we grafted the proteins. We added a quantity of protein such that, if 90% of the protein-DBCO successfully conjugated to the azide-liposomes, we would have 200/150/75/50/25/12/6 proteins per liposome, as determined by the numerical concentration of liposomes measured with nanoparticle tracking analysis. We radiolabeled a tracer portion of the proteins before conjugation. We ran the protein/liposomes reaction mixture through a size exclusion column to separate the unconjugated protein from the liposomes. After calculating liposome-associated protein signal/(liposome-associated protein signal + unconjugated protein signal) and checking that (liposome-associated signal + unconjugated signal) was roughly equal to the total added signal, we determined the efficiency of protein-liposome conjugation. We took this efficiency and determined the actual number of proteins added per liposome for each reaction condition. The final corrected protein per liposome value was then combined with the measured liposome surface area to determine the surface area footprint of each conjugated protein. Assuming a square footprint, we determined the average protein-protein spacing.

For E. coli, we used an analogous calculation to that used for the liposomes. The bacteria surface area was determined based on a theoretical shape: a 1 µm × 0.5 µm cylinder with 0.5 µm diameter hemispheres at either end. Bacterial concentration was determined by measurement of optical density at 600 nm. We used a ninhydrin assay to optically detect surface nucleophiles on the bacteria. Analogously to calculation of protein spacing on liposomes, we used the bacterial surface area and the ninhydrin-determined concentration of nucleophiles to determine average nucleophile density/spacing on the bacteria, assuming a square footprint on the bacteria surface for each nucleophile. Published proteomics data has characterized the amino acid sequences of proteins on E. coli surfaces, allowing us to determine an average number of nucleophiles per protein on E. coli. This allows conversion from nucleophile surface density to average protein surface density/spacing. To adjust this effective protein spacing, we adjusted the density of nucleophiles on the surface of the bacteria by conjugating methyl-terminated nucleophile-reactive compounds to the nucleophile groups. As with unmodified bacteria, ninhydrin assay determined nucleophile concentrations, in turn allowing determination of the adjusted protein density/spacing.

## Acknowledgements

We thank Penn Cytomics and Cell Sorting Shared Resource Laboratory at the University Pennsylvania, which is partially supported by Abramson Cancer Center NCI Grant (P30 016520) and the research identifier number is RRid: SCR_022376.

## Funding

National Institutes of Health grant R01-HL-153510 (JSB)

National Institutes of Health grant R01-HL160694 (JSB)

PhRMA Foundation Postdoctoral Fellowship in Drug Delivery 2023 PFDL 1008128 (ZW)

American Heart Association 916172 (JN)

National Institutes of Health grant R01-HL160694 (WS)

National Institutes of Health grant R01-HL-157189 (VRM)

Pennsylvania Department of Health Research Formula Fund (VRM)

National Institutes of Health grant 1R35GM136259 (RR)

National Institutes of Health grant 1R01CA244660 (RR)

PECASE award W911NF1910240 (JF)

Department of Defense DURIP W911NF2010107 (JF)

National Science Foundation MRI 2215917 (JF)

Chancellor’s Grant for Independent Student Research at Rutgers University–Camden (AE)

Equipment Leasing Funds from the State of New Jersey (JF, AE)

## Author contributions

Conceptualization: ZW, SK, RR, JWM, JSB

Methodology: ZW, SK, JN, MZ, AE, EH, TS, MZ, DG, EW, CE, EA, YW, OAM-C, JWM

Visualization: ZW, SK, JN, JWM

Funding acquisition: JF, VRM, RR, JSB

Supervision: RR, JWM, JSB,

Writing – original draft: ZW, SK, RR, JWM, JSB

Writing – review & editing: ZW, SK, JN, OAM-C, WS, VRM, JF, RR, JWM, JSB

## Competing interest

Authors declare that they have no competing interests.

## Data and materials availability

All data are available in the main text or the supplementary materials.

## Supplementary Materials

### SUPPLEMENTARY SECTION A: Supplementary Figures

**Supplementary Figure 1.**
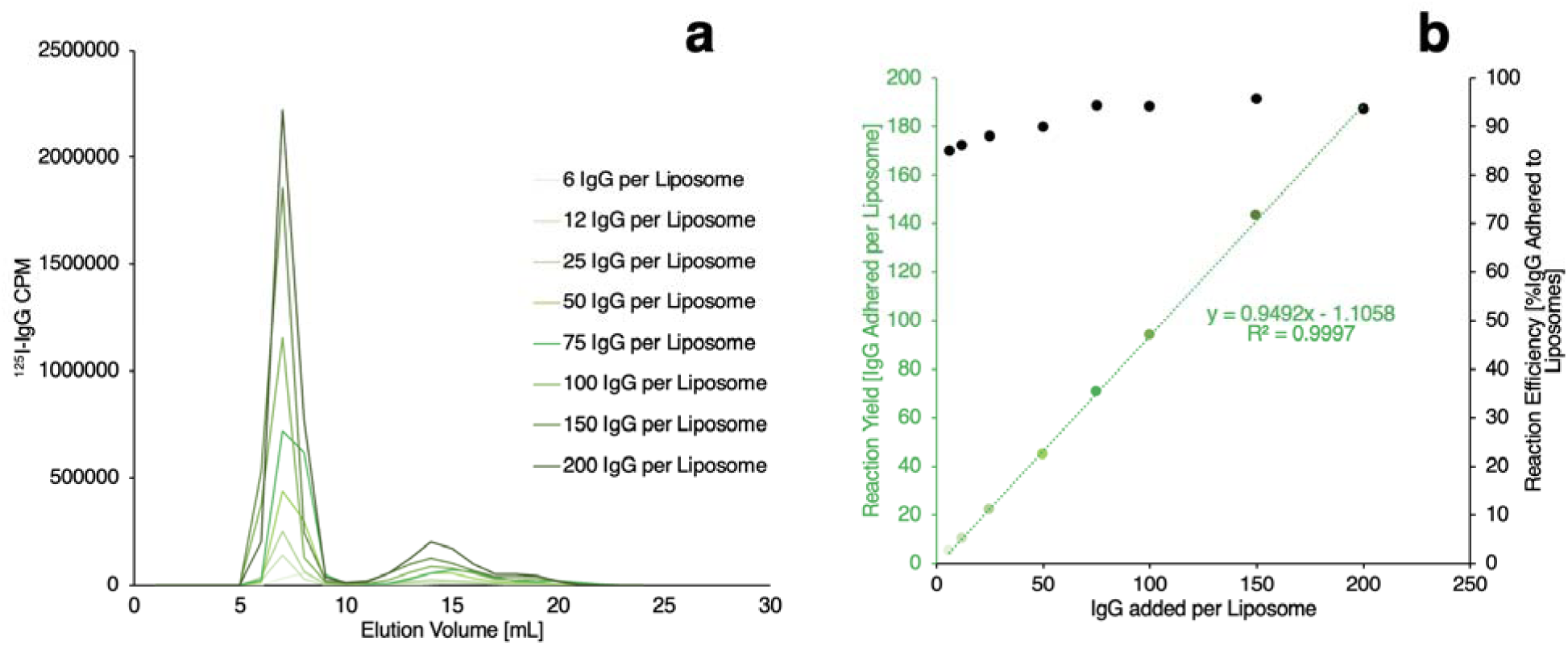
Characterization of IgG conjugation to liposome surfaces. (a) IgG was functionalized with dibenzocyclooctyne and was labeled with Iodine-125. Labeled and alkyne-functionalized IgG was reacted with azide-presenting liposomes and the mixture of IgG-liposomes and unreacted IgG was applied to a size-exclusion chromatography column. Liposomes eluted through the column over 5-10 minutes and IgG eluted through the column over 12-20 minutes. Radioactivity from Iodine-125-labeled IgG (Iodine-125 counts per minute/CPM) was traced in elution fractions to generate chromatographs with peaks for IgG-conjugated liposomes and unconjugated IgG. (b) Area under curve for the IgG-liposome peak and the unconjugated IgG peak was determined to calculate efficiency of IgG-liposome coupling (black data points, right axis). Based on calculated reaction efficiency and the numerical concentrations of IgG and liposomes added to the IgG-liposome reactions, the yield number of IgG conjugated per liposome was determined (green data points, left axis).

**Supplementary Figure 2.**
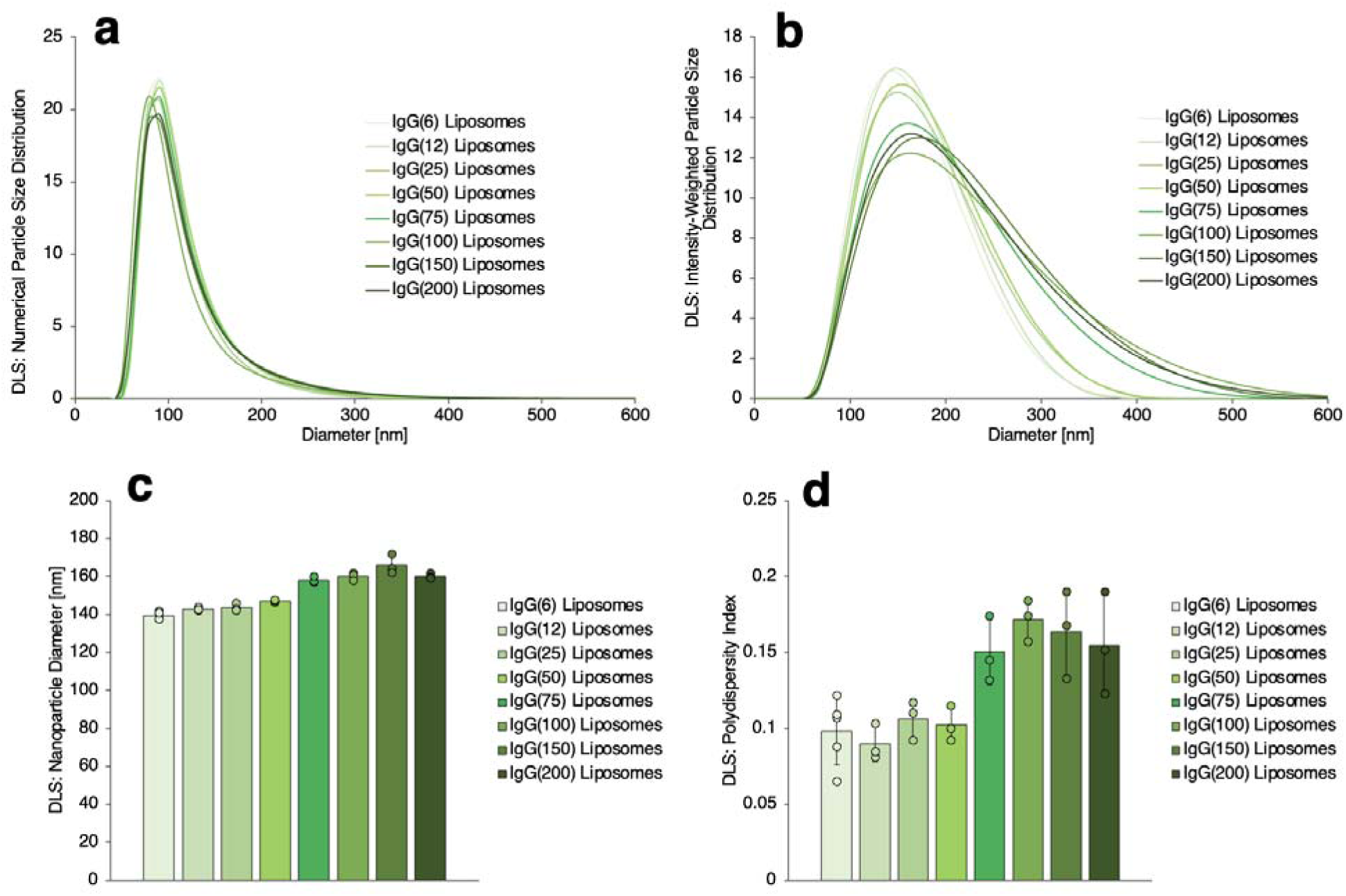
Dynamic light scattering sizing of IgG-liposomes. (a-b) Numerical (a) and intensity-weighted (b) size distributions of IgG-conjugated liposomes. (c) Mean ± standard error diameters of IgG-liposomes determined from size distributions in (a-b). (d) Polydispersity indices of IgG-liposomes determined from the size distributions in (a-b).

**Supplementary Figure 3.**
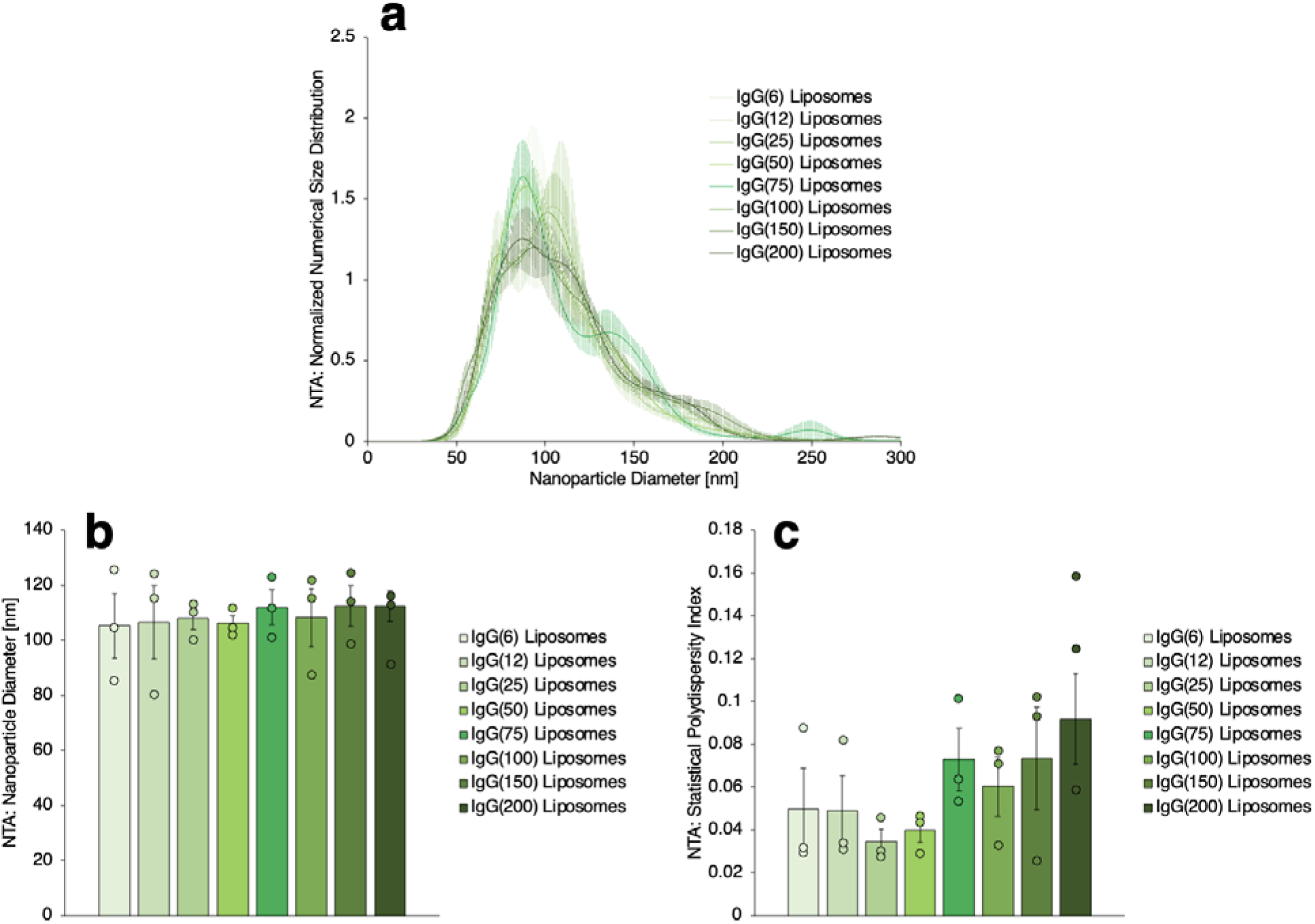
Nanoparticle tracking analysis (NTA) characterization of IgG-liposomes. (a) Numerical size distributions of IgG-liposomes with different numbers of IgG per liposome, showing good agreement with dynamic light scattering data in Supplementary Figure 2a. (b) Mean ± standard error nanoparticle diameters as determined by size distribution data in (a). (c) Statistical polydispersity indices as determined from the size distribution data in (a).

**Supplementary Figure 4.**
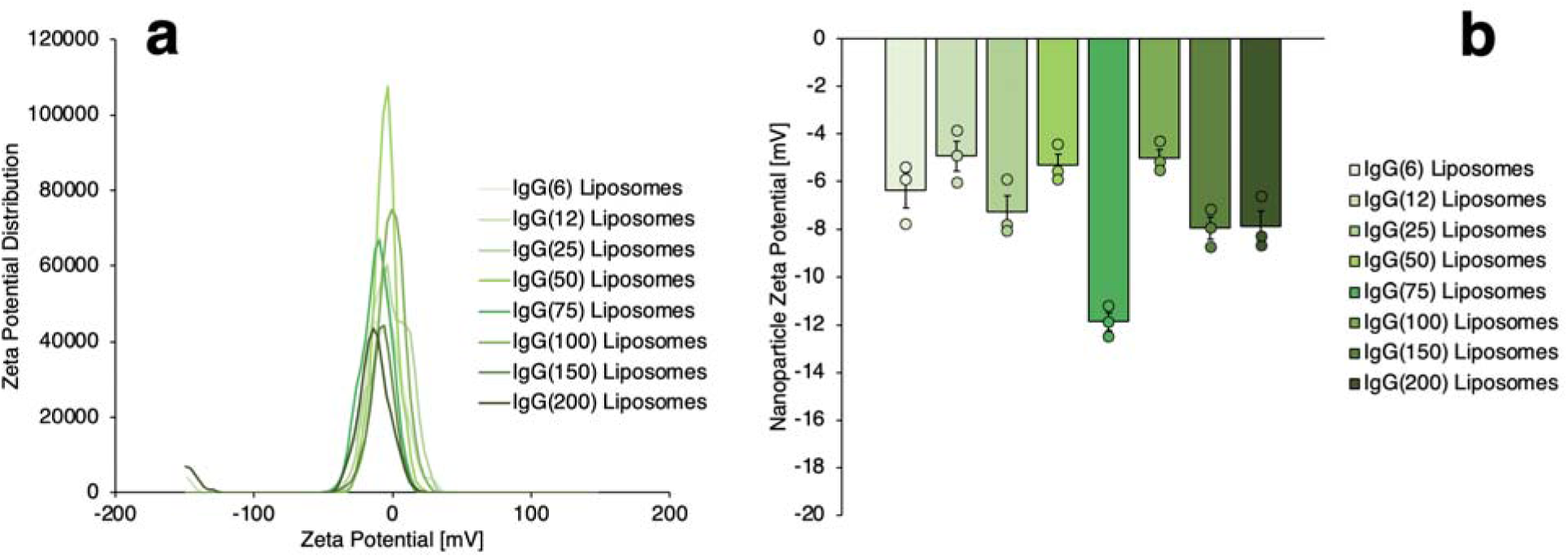
Zeta potential determination for IgG-liposomes. (a) Zeta potential frequency distributions for IgG-liposomes with different numbers of IgG per liposome. (b) Mean ± standard error zeta potentials as determined from distributions in (a).

**Supplementary Figure 5.**
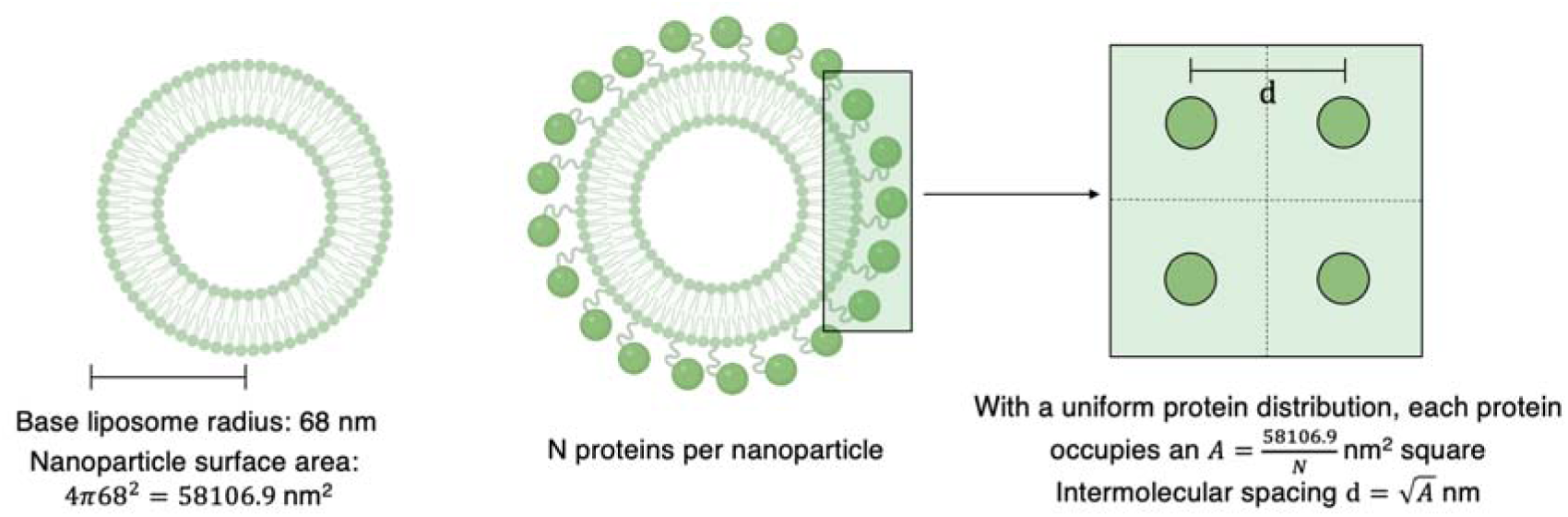
Schematic for calculation of surface protein spacing on liposomes. Based on data in Supplementary Figure 1, we determine the number of proteins per liposome. Based on data in Supplementary Figures 2-3, we determine the surface area of the liposomes to which the proteins are bound. Using these two values, we calculate the average protein-protein spacing as depicted.

**Supplementary Figure 6.**
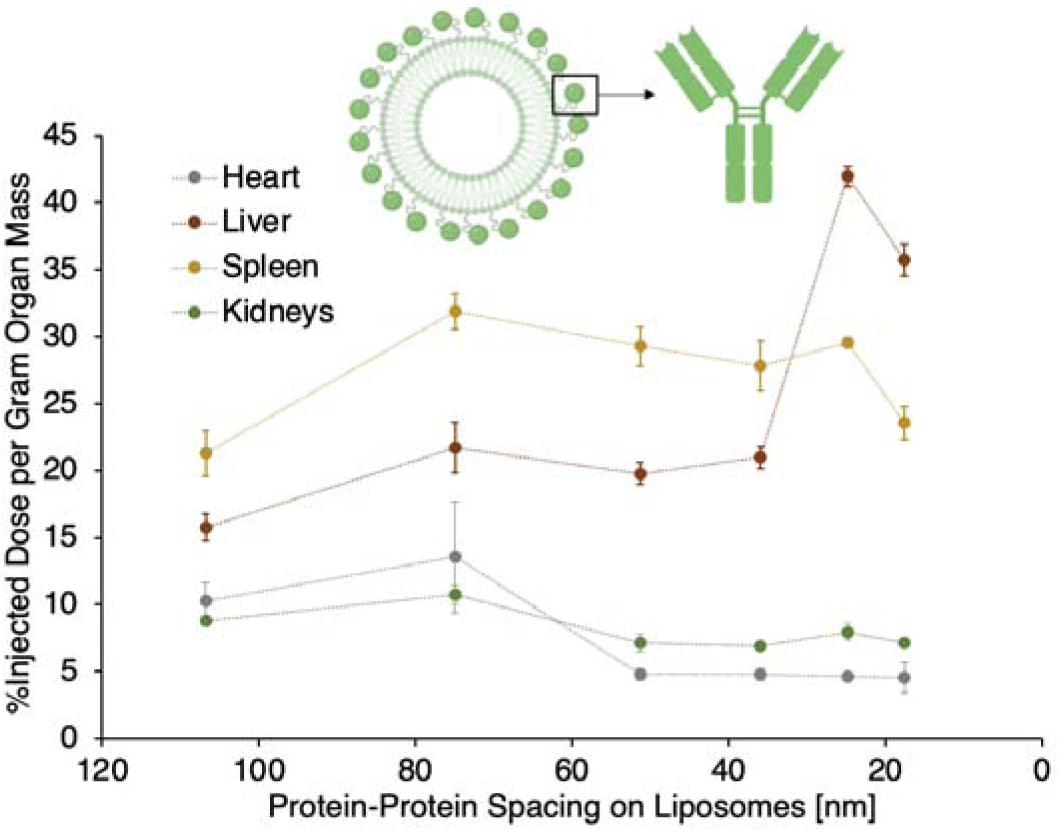
Biodistribution data for IgG-liposomes in mice with intravenous LPS-induced acute lung inflammation, as a function of protein-protein spacing on the liposomes. The data parallel that in Figure 1b in the main text, which show blood retention and lung uptake for the same nanoparticles in the same experiments.

**Supplementary Figure 7.**
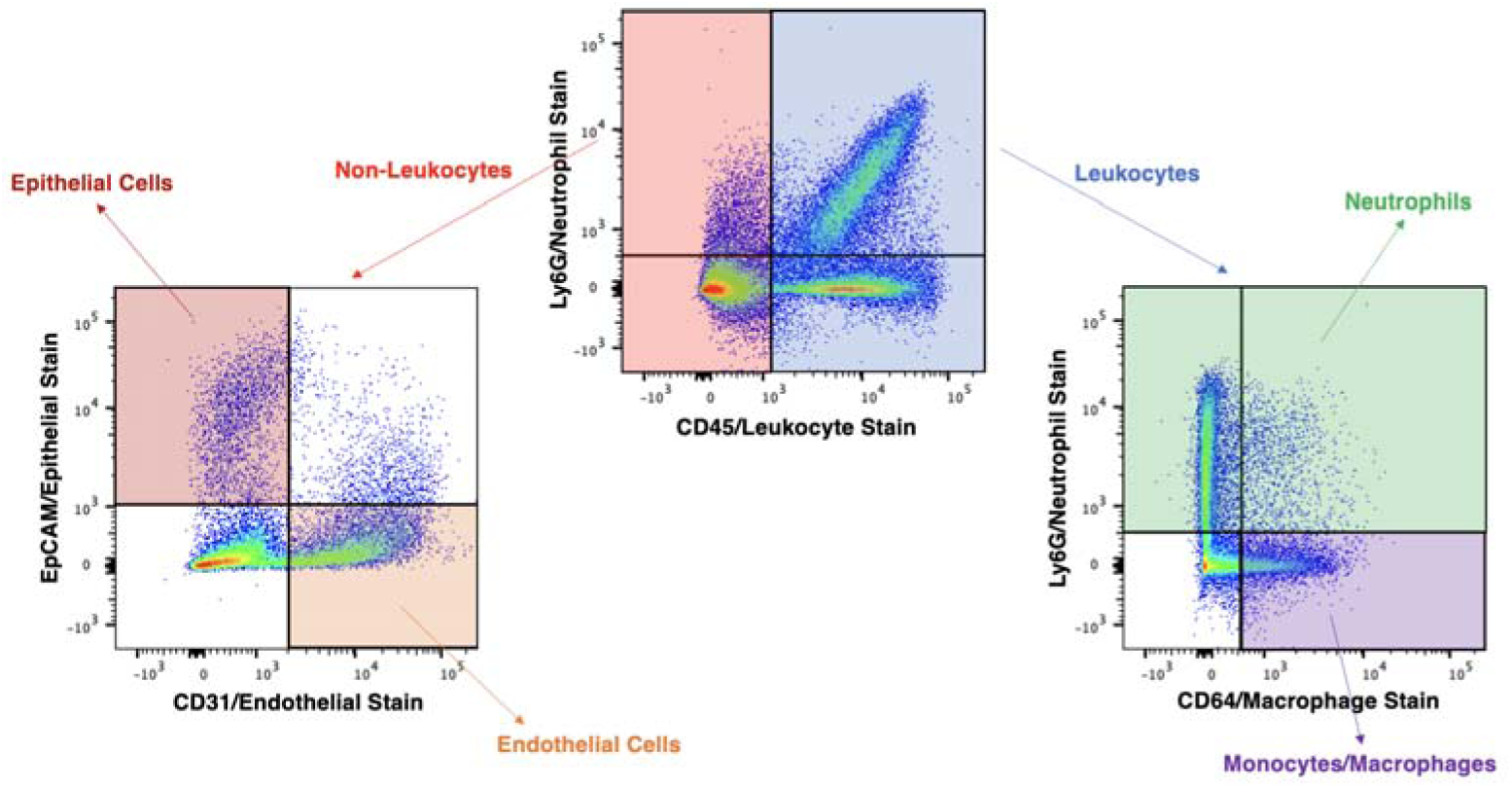
Diagram of gating strategies used to obtain lung flow cytometry data in Figure 1c-d in the main text and Supplementary Figures 8-9, for mice with intravenous LPS-induced acute lung inflammation. All identified cells were first gated as leukocytes and non-leukocytes by CD45 stain (top center panel). Non-leukocytes (left panel) were identified as endothelial cells by CD31 (PECAM) stain or epithelial cells by epithelial cell adhesion molecule (EpCAM) stain. Leukocytes (right panel) were identified as neutrophils by Ly6G stain or macrophages/monocytes by CD64 stain among Ly6G-negative leukocytes.

**Supplementary Figure 8.**
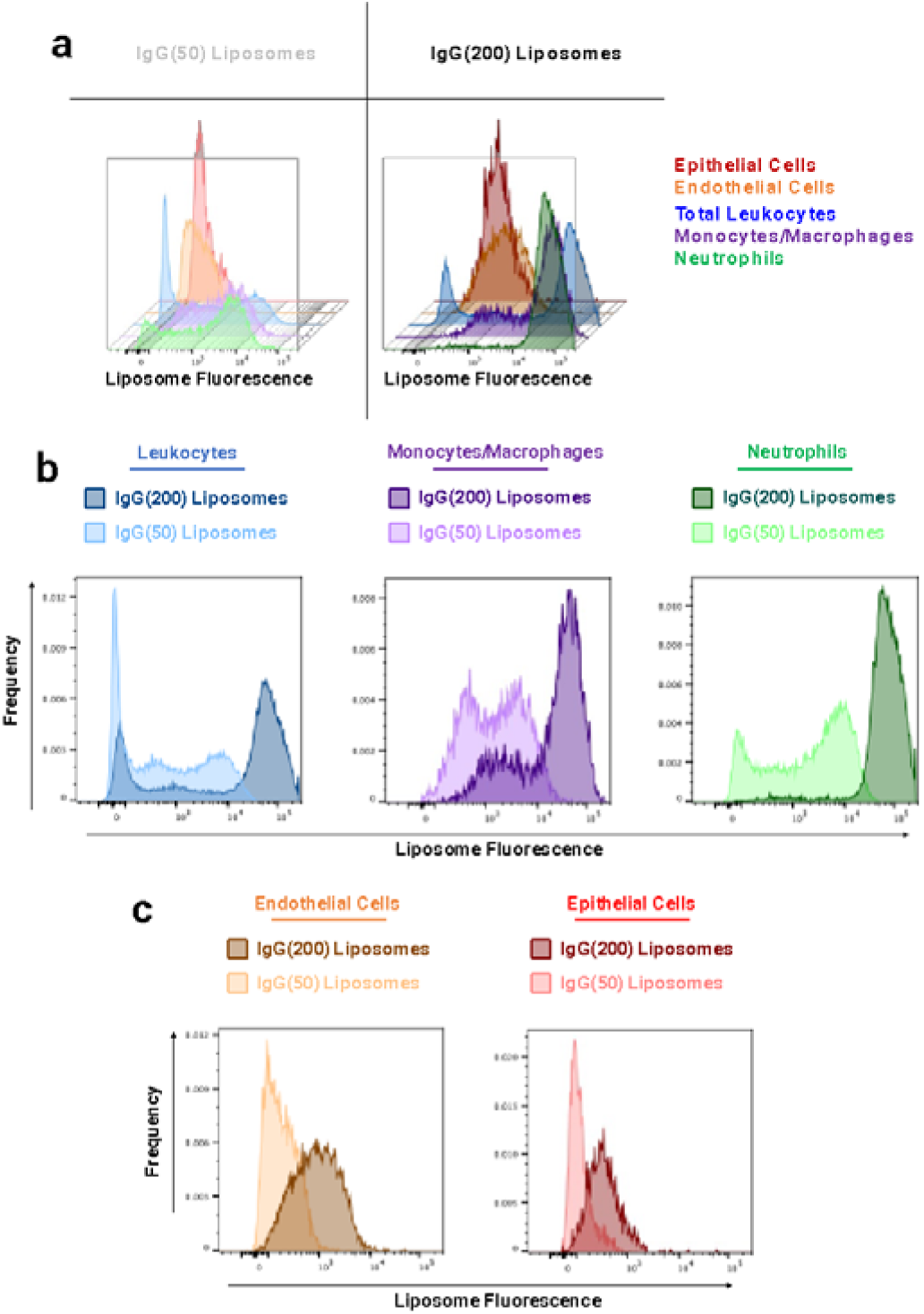
(a) Flow cytometry data indicating IgG-liposome fluorescence intensity associated with all tested cell types in the lungs of mice with intravenous LPS-induced acute lung inflammation for IgG(50) liposomes (36 nm protein-protein spacing, left panel) and IgG(200) liposomes (18 nm protein-protein spacing, right panel). (b) Overlaid histograms of IgG(50) (light colors) and IgG(200 (dark colors) fluorescence intensity in total lung leukocytes, lung monocytes/macrophages, and lung neutrophils. The neutrophil data are reproduced in Figure 1c. For all leukocytes, there is a large subpopulation of high-liposome fluorescence cells for IgG(200) liposomes that is not found with IgG(50) liposomes. (c) Overlaid histograms of IgG(50) (light colors) and IgG(200) (dark colors) fluorescence intensity in endothelial cells and epithelial cells. There is a small but significant shift to higher nanoparticle uptake for IgG(200) liposomes vs. IgG(50) liposomes.

**Supplementary Figure 9.**
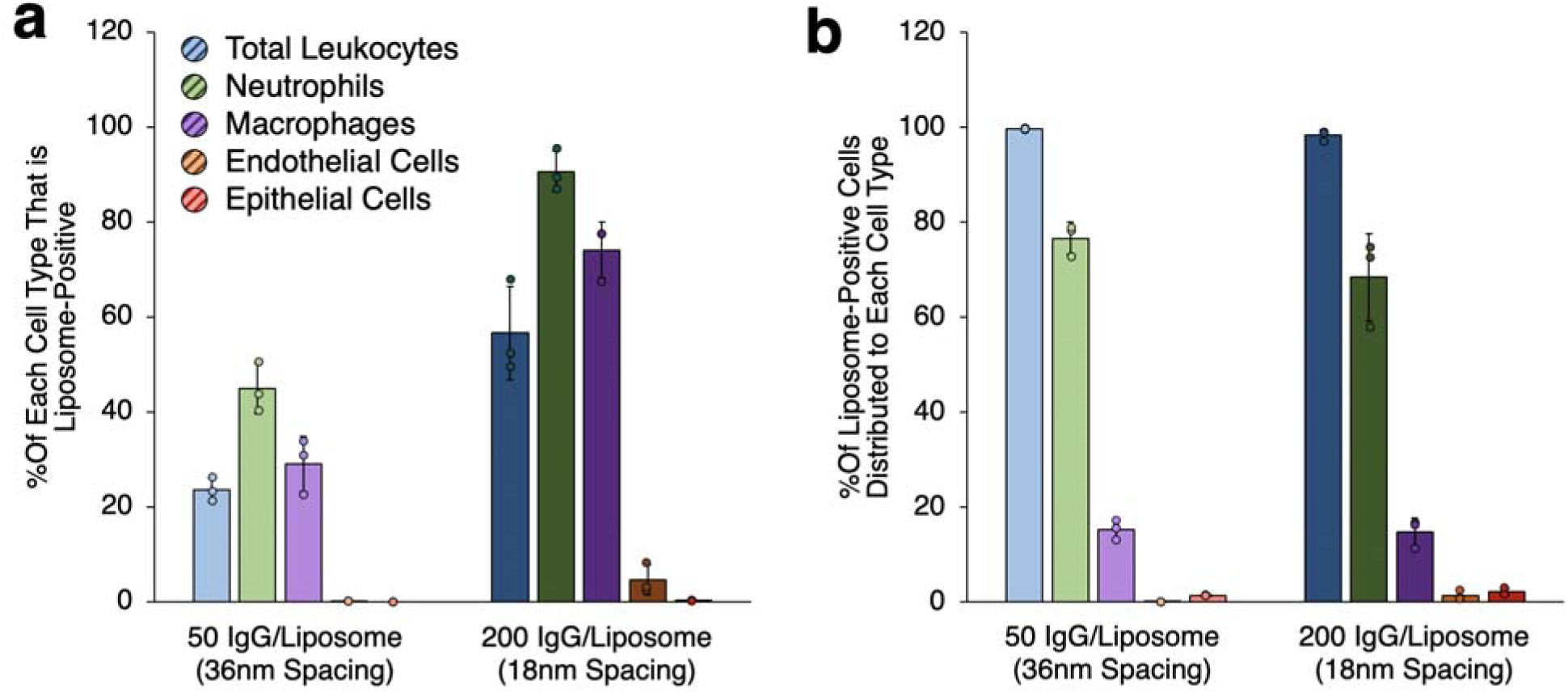
Summary flow cytometry data describing distribution of IgG-liposomes to different cell types in the lungs of mice with intravenous LPS-induced acute lung inflammation. (a) Percentage of each cell type positive for IgG(50)-liposomes (36 nm protein-protein spacing, light colors) and IgG(200) liposomes (18 nm protein-protein spacing, dark colors). Both types of liposomes distribute more effectively to leukocytes vs. epithelial or endothelial cells, but reducing the IgG-IgG spacing enhances liposome uptake in all cell types. (b) Percentage of liposome-positive cells that are each cell type for IgG(50)-liposomes (36 nm protein-protein spacing, light colors) and IgG(200) liposomes (18 nm protein-protein spacing, dark colors). Nearly all liposome-positive cells were leukocytes, and among leukocytes, ∼80% of liposome-positive cells were neutrophils. The distribution of liposome-positive cells among different cell types did not change significantly between IgG(50) and IgG(200) liposomes.

**Supplementary Figure 10.**
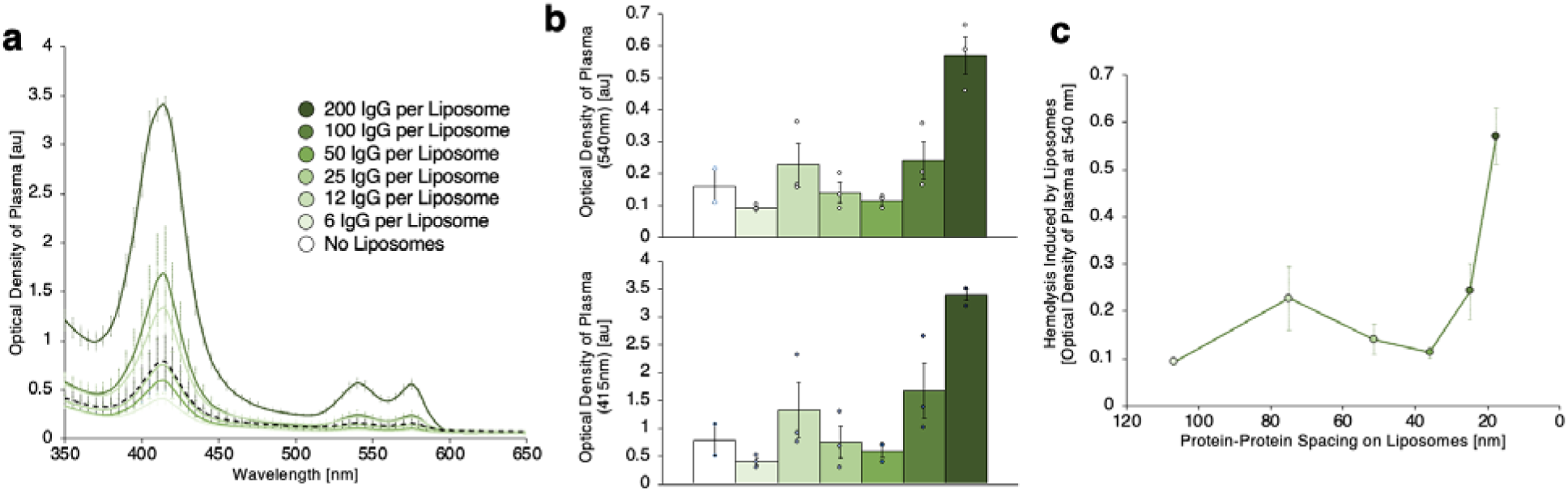
Optical density of plasma from mice with intravenous LPS-induced acute lung inflammation treated with IgG-liposomes with different IgG surface densities, indicating hemoglobin levels in the plasma and thus hemolysis induced by the nanoparticles. The spectra in (a) are reproduced in Figure 1e in the main text.The data in (b) present optical density of plasma at 540 nm and 415 nm. (c) Reducing IgG-IgG spacing on the administered liposomes below 36 nm results in a steep increase in optical density at 540 nm, identical to the threshold shown in Figure 1f in the main text.

**Supplementary Figure 11.**
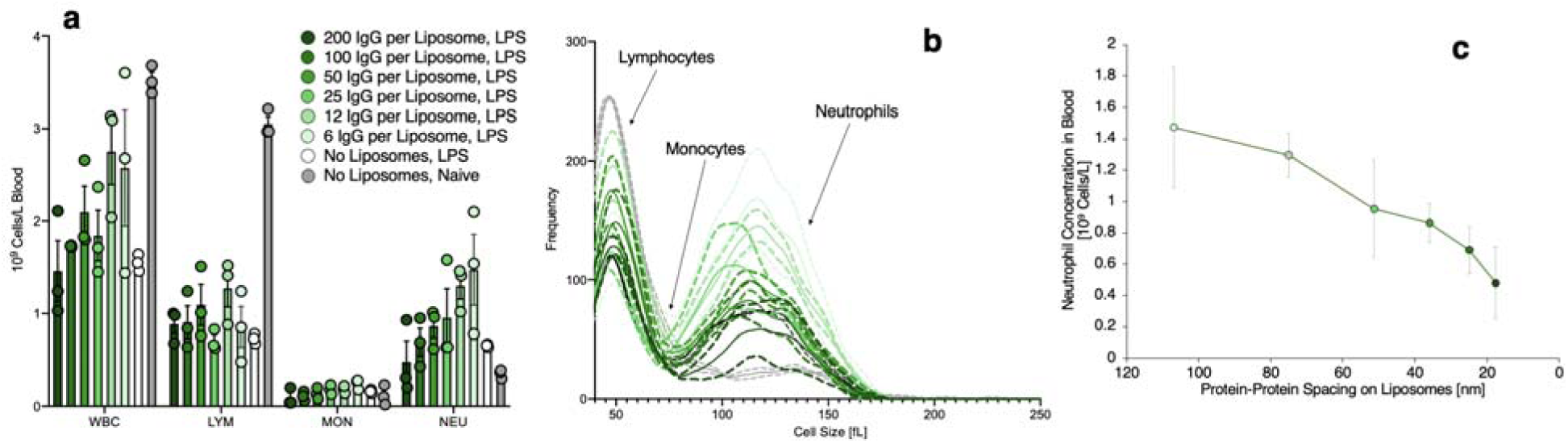
Complete blood count assessment of leukocytes in blood from mice with intravenous LPS-induced acute lung inflammation treated with IgG-liposomes. (a) Numerical concentrations of total white blood cells (WBC), lymphocytes (LYM), monocytes (MON), and neutrophils (NEU) in blood from mice treated with IgG-liposomes with different surface densities of IgG. Data for mice with intravenous LPS-induced acute lung inflammation that did not receive liposomes and data for naïve mice that received neither LPS nor liposomes are included for comparison. (b) Data for mice treated as in (a) showing leukocyte size distributions and indicating both concentration and size of each type of leukocyte evaluated in the assay. Solid lines indicate averages and dashed lines indicate ± standard error of the mean. (c) Neutrophil concentrations in blood from mice treated with IgG-liposomes as a function of IgG-IgG spacing on the liposomes.

**Supplementary Figure 12.**
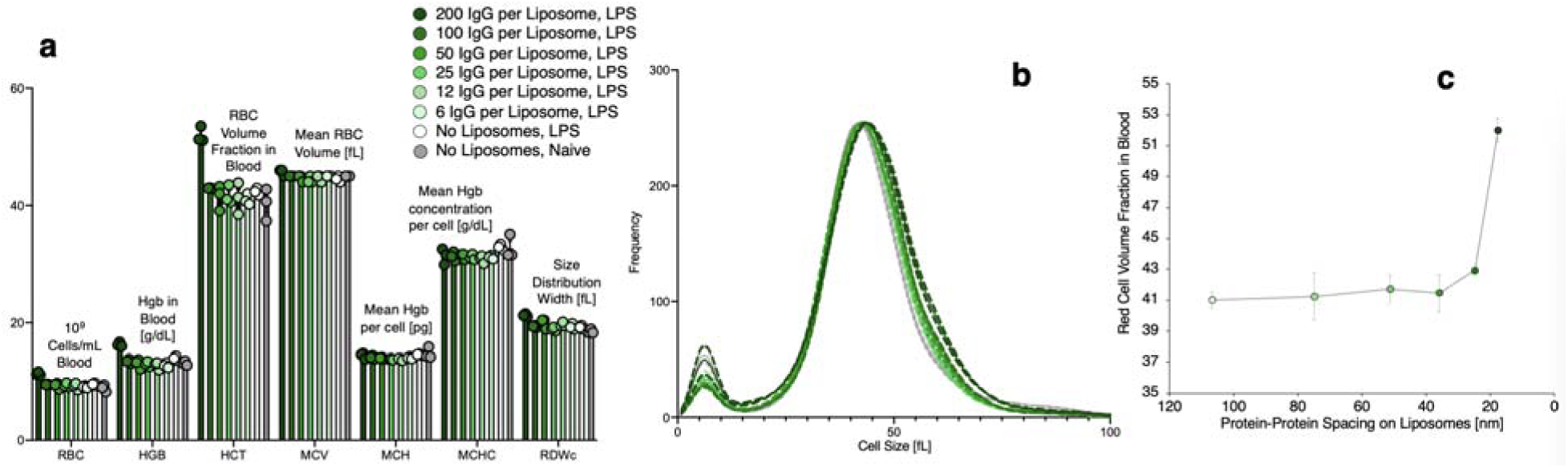
Complete blood count assessment of red blood cells in blood from mice with intravenous LPS-induced acute lung inflammation treated with IgG-liposomes. (a) Data indicating red blood cell concentrations, hemoglobin concentrations, hematocrit, mean red blood cell volume, mean hemoglobin per red blood cell, mean hemoglobin concentration per cell, and dispersity of the red blood cell size distributions for mice treated with IgG-liposomes with different surface densities of IgG. Data for naïve and IV-LPS-injured mice that did not receive liposomes are included for comparison. IgG-liposomes with 200 IgG per liposome are the only tested liposome that induced significant changes in red blood cell concentration and size. (b) Data for mice treated as in (a) showing red blood cell size distributions. For each distribution, the larger peak shows data for red blood cells and the smaller peak shows data for platelets. Solid lines indicate averages and dashed lines indicate ± standard error of the mean. (c) Hematocrit in IgG-liposomes-treated mice with acute lung inflammation as in (a), presented as a function of IgG-IgG spacing on liposomes, showing a steep increase in hematocrit for IgG-IgG spacing on liposomes lowered below 25 nm.

**Supplementary Figure 13.**
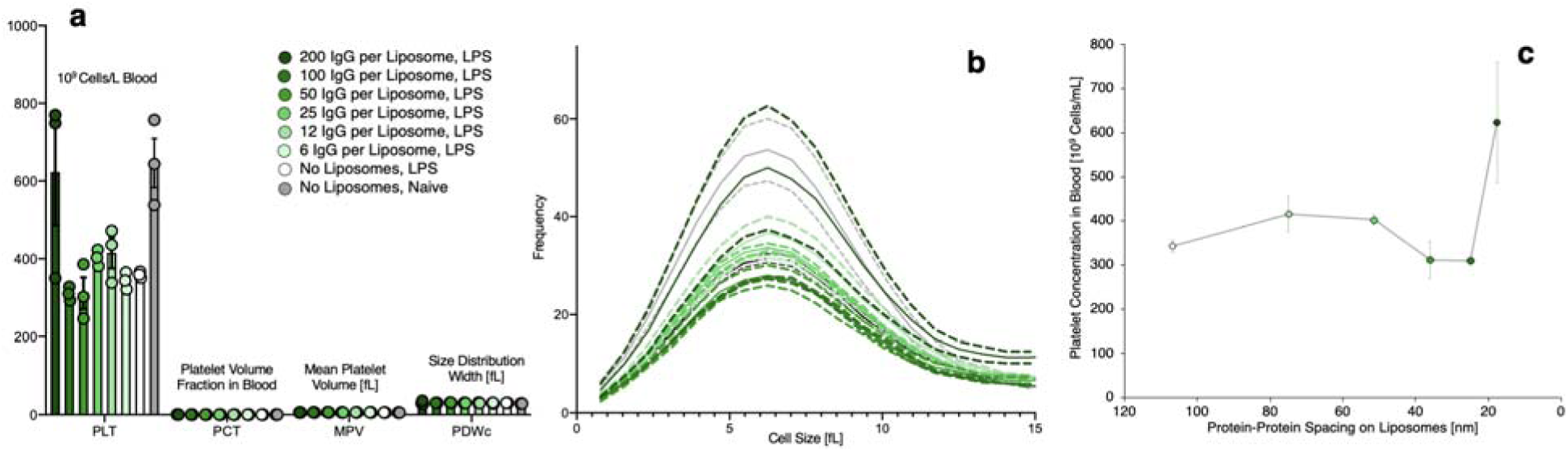
Complete blood count assessment of platelets in blood from mice with intravenous LPS-induced acute lung inflammation treated with IgG-liposomes. (a) Data indicating platelet concentrations, platelet volume fraction in blood, mean platelet volume, and dispersity of the platelet size distributions for mice treated with IgG-liposomes with different surface densities of IgG. Data for naïve and IV-LPS-injured mice that did not receive liposomes are included for comparison. (b) Data for mice treated as in (a) showing platelet size distributions. Solid lines indicate averages and dashed lines indicate ± standard error of the mean. (c) Platelet concentration in IgG-liposomes-treated mice with acute lung inflammation as in (a), presented as a function of IgG-IgG spacing on liposomes. A change in platelet concentration is noted only for IgG liposomes with the smallest IgG-IgG spacing.

**Supplementary Figure 14.**
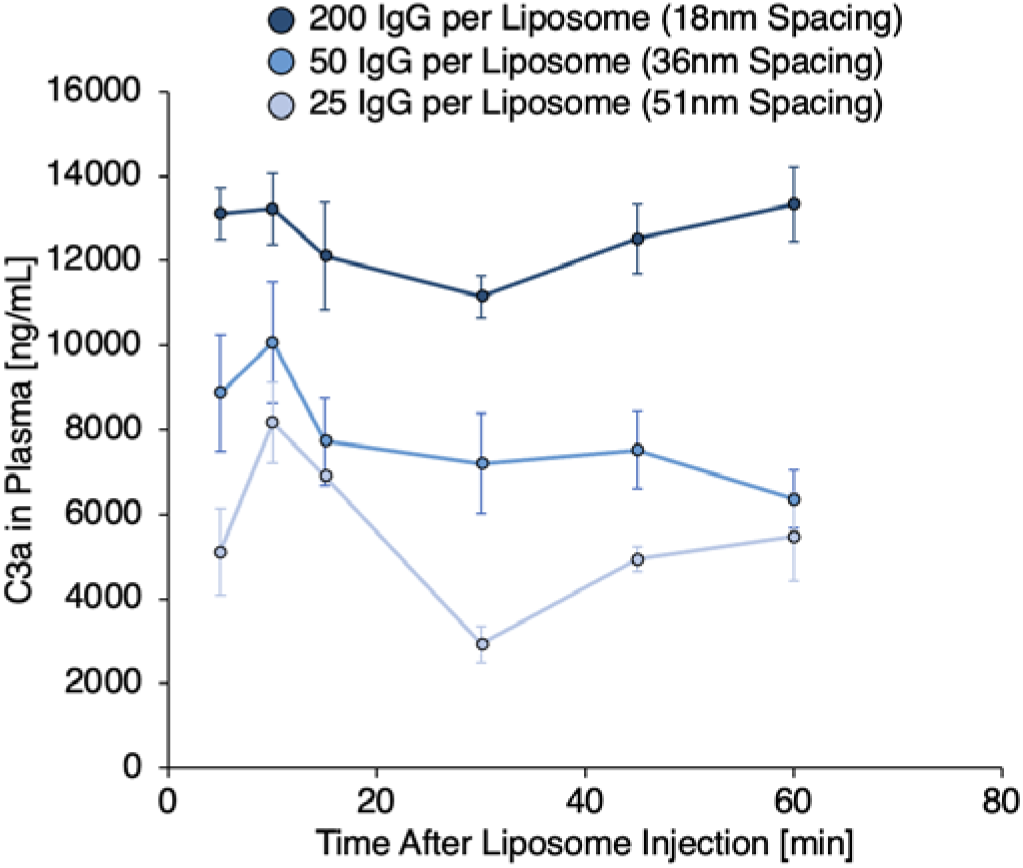
Plasma concentrations of complement fragment C3a in mice treated with IgG-liposomes with 200, 50, or 25 IgG per liposome, corresponding to surface IgG-IgG spacing of 18 nm, 36 nm, and 51 nm, respectively. Blood was obtained from mice at 5, 10, 15, 30, 45, and 60 minutes after liposome boluses. Data at 60 minutes are presented in Figure 1g in the main text as a function of IgG-IgG spacing on the liposomes.

**Supplementary Figure 15.**
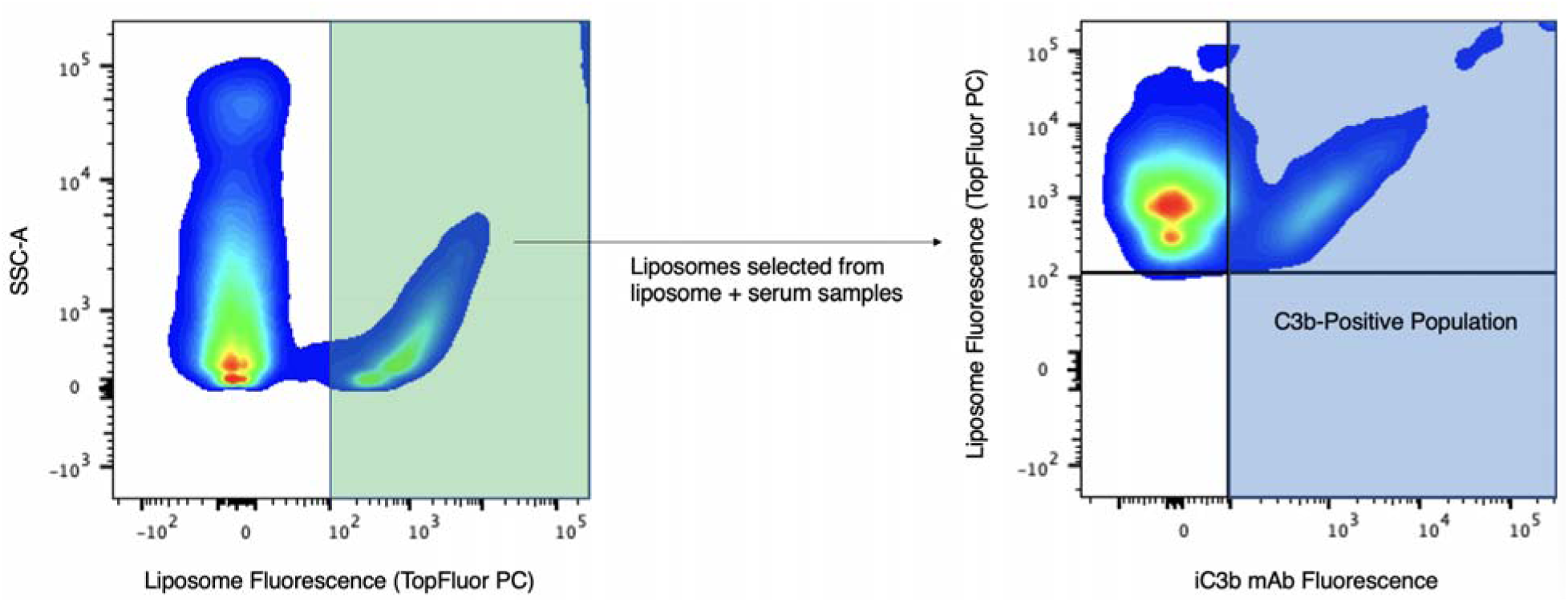
Diagram of gating strategies used to obtain single nanoparticle flow cytometry data in Figure 2a-c and 2h-j in the main text and in Supplementary Figures 16-17. Liposomes were incubated with serum then stained with fluorescent anti-C3b antibody. Left panel: Liposomes were distinguished from endogenous materials in serum by tracing fluorescent lipid (TopFluor PC) added to the liposome formulation. Among events identified as liposomes by the depicted gate (green), C3b fluorescence was quantified (right panel).

**Supplementary Figure 16.**
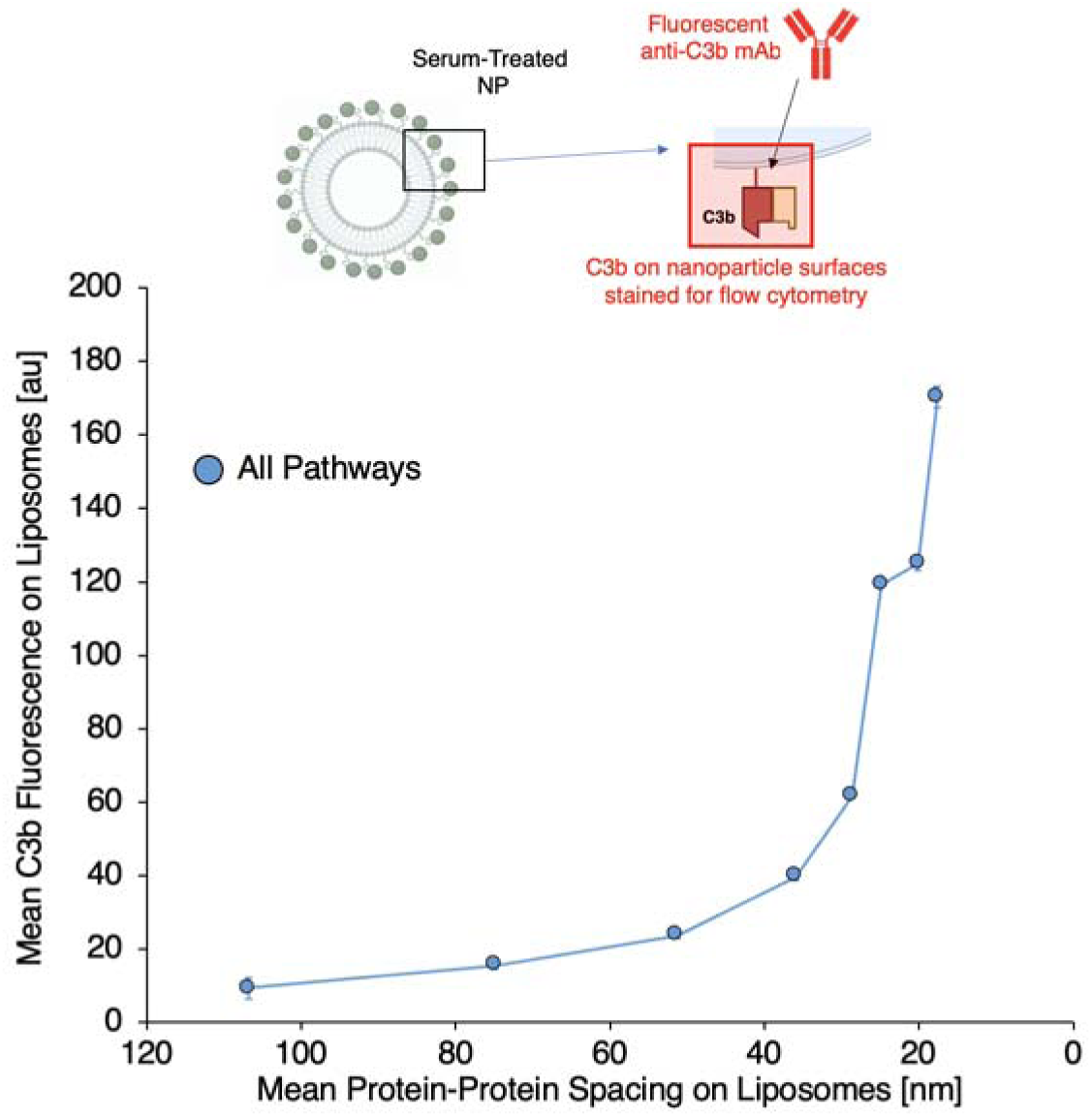
Flow cytometry data indicating fluorescent C3b stain on IgG-liposomes with different IgG-IgG intermolecular spacing after incubation with serum. Data are as in Figure 2b in the main text but show findings for cumulative C3b deposition on nanoparticles by all complement pathways, rather than the isolated alternative pathway or classical+lectin pathways. Data for the isolated classical+lectin pathways is determined as the difference between data for all pathways, as presented in this figure, and data for the alternative pathway, as presented by the red curve in Figure 2b. The data indicate a steep increase in C3b deposition on IgG-liposomes for IgG-IgG spacing < 36 nm, but also show, by comparison to Figure 2b, that alternative pathway-based C3b deposition is the portion of the complement response to the nanoparticle surface that is most sensitive to spacing of surface proteins on the nanoparticle.

**Supplementary Figure 17.**
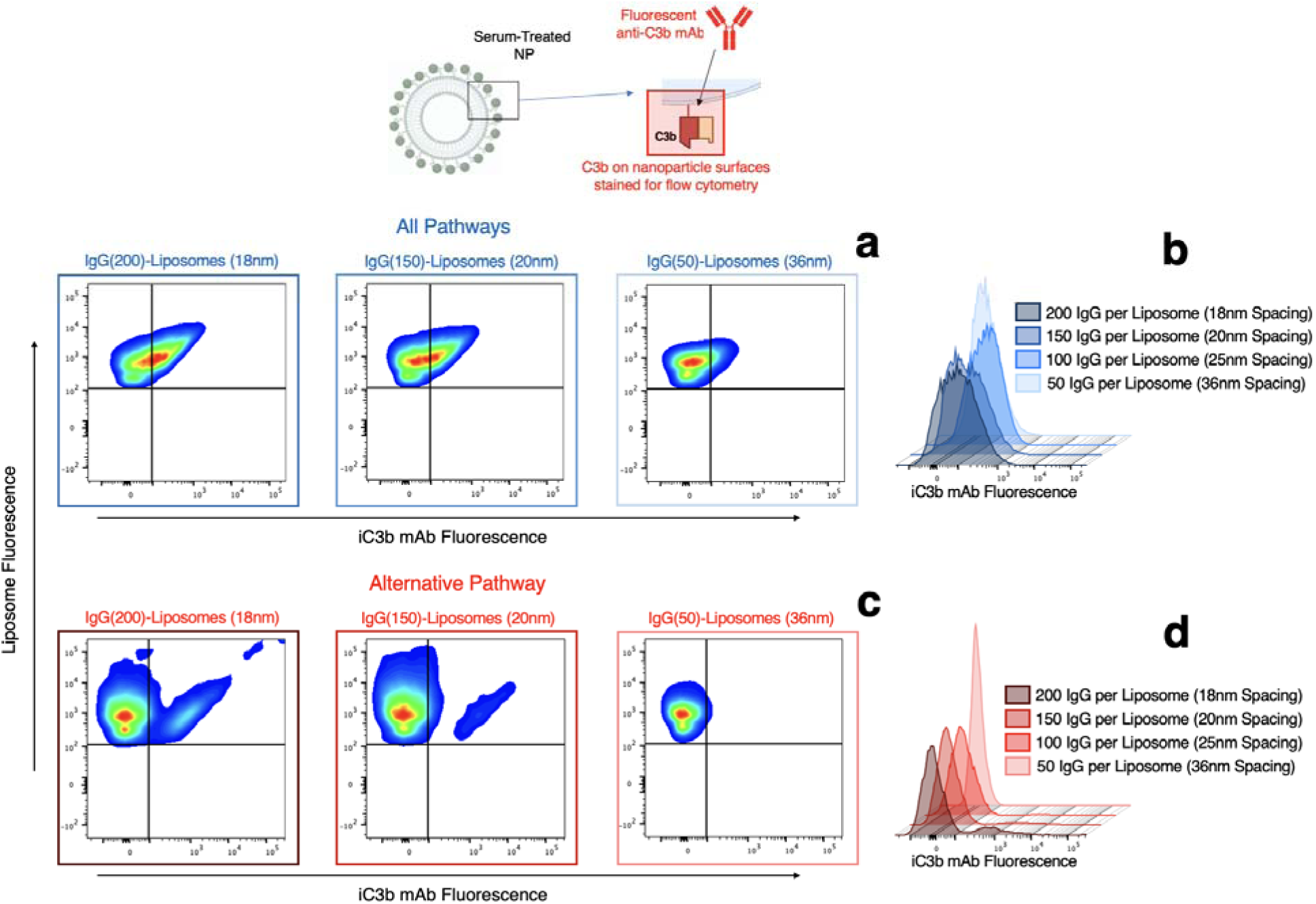
Flow cytometry data indicating fluorescent C3b stain on IgG-liposomes with different IgG-IgG intermolecular spacing after incubation with serum. Data show two-dimensional and histogram depictions of the fluorescence distributions that determined the plots in Supplementary Figure 16 and main text Figure 2b. Data in (a) and (b) show cumulative C3b deposition on liposomes by all complement pathways, as in Supplementary Figure 16. (a) Two-dimensional plots of liposome fluorescence vs. anti-C3b stain fluorescence. (b) Histograms of anti-C3b stain fluorescence on IgG-liposomes with different IgG-IgG surface spacings. Data in (c) and (d) are equivalent to data presented in Figure 2c and the Figure 2d inset in the main text, and are reproduced here for comparison between the alternative pathway response to the liposomes and the total complement response to the liposomes. While the cumulative complement response (a-b) causes more C3b deposition than the isolated alternative pathway response, the bimodal high-C3b populations for nanoparticles with higher surface protein densities are more readily apparent in data that isolates the alternative pathway response to the nanoparticles.

**Supplementary Figure 18.**
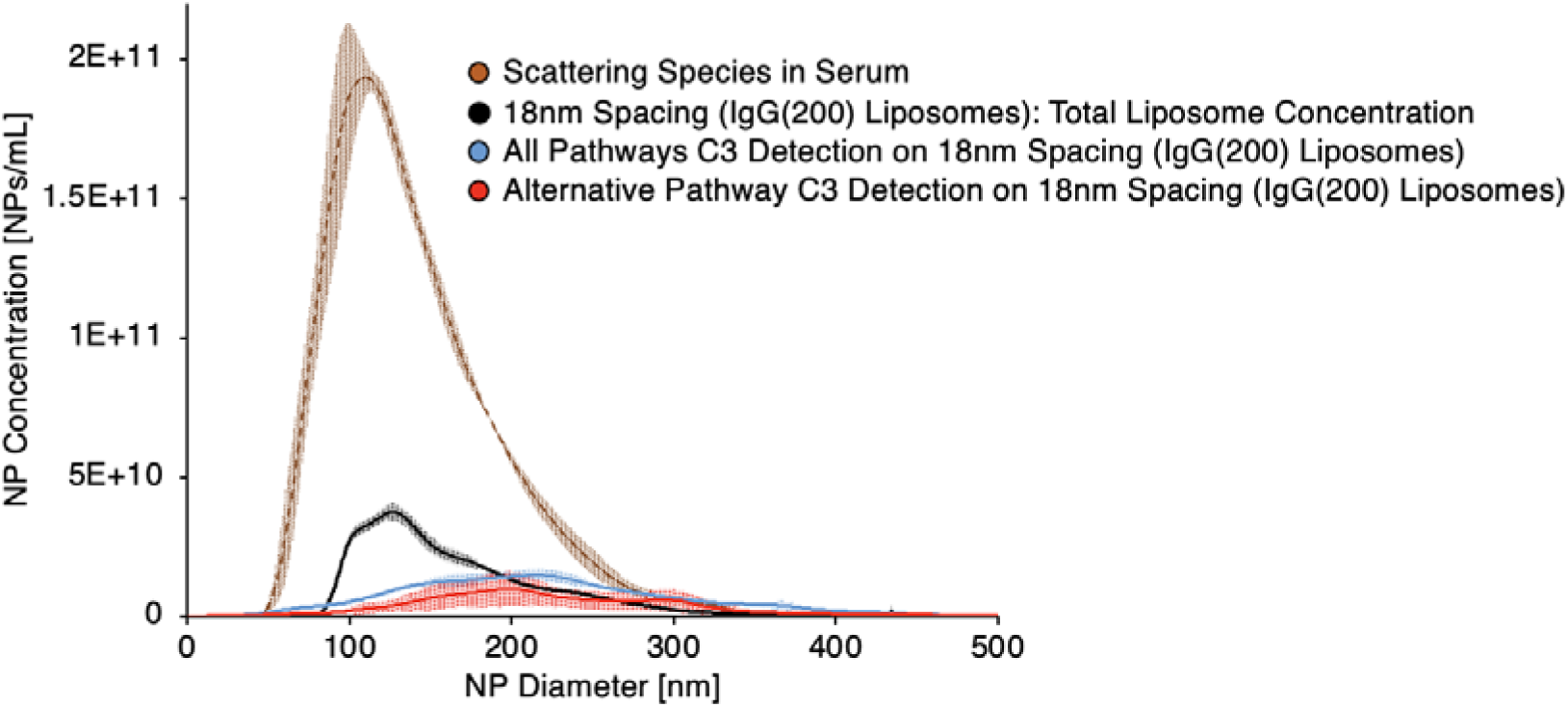
Nanoparticle tracking analysis (NTA) data indicating concentration vs. size distributions of endogenous light scattering species in serum (brown), IgG-liposomes with 200 IgG per liposome prior to being added to serum (black), the same IgG-liposomes as detected in serum via adhesion of fluorescent C3 to the liposomes (blue), and the same IgG-liposomes as detected in serum with gelatin veronal buffer (isolating the alternative pathway of complement) via adhesion of fluorescent C3 to the liposomes (red). Note that the concentration and size of the endogenous species is different from the concentration and size of the species detected in serum via fluorescent C3 interactions with liposomes in the serum. Liposomes detected via C3-liposome interactions are larger than the same liposomes before incubation with serum.

**Supplementary Figure 19.**
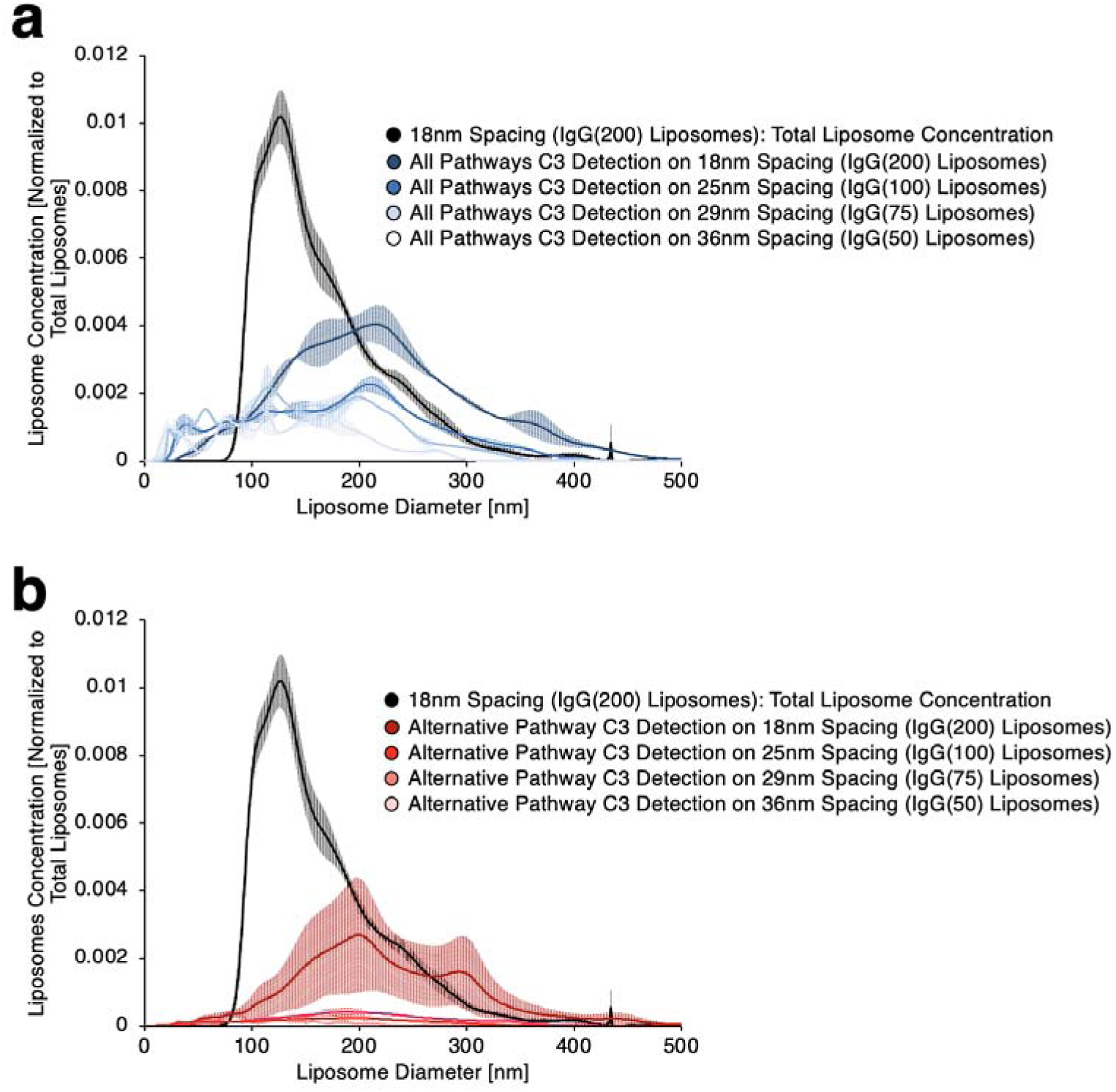
NTA data indicating concentration vs. size distributions of IgG-liposomes with 200 IgG per liposome prior to being added to serum (black) and of IgG-liposomes detected in serum via adhesion of fluorescent C3 to the liposomes, for liposomes with different surface protein densities. (a) Data for C3-liposome interactions in serum, indicating C3 deposition via all complement pathways. (b) Data as in (a), but for serum with gelatin veronal buffer, isolating C3 deposition via the alternative pathway. Data in (b) is reproduced from Figure 2h in the main text, shown here in juxtaposition to data in (a) for the cumulative effects of all complement pathways.

**Supplementary Figure 20.**
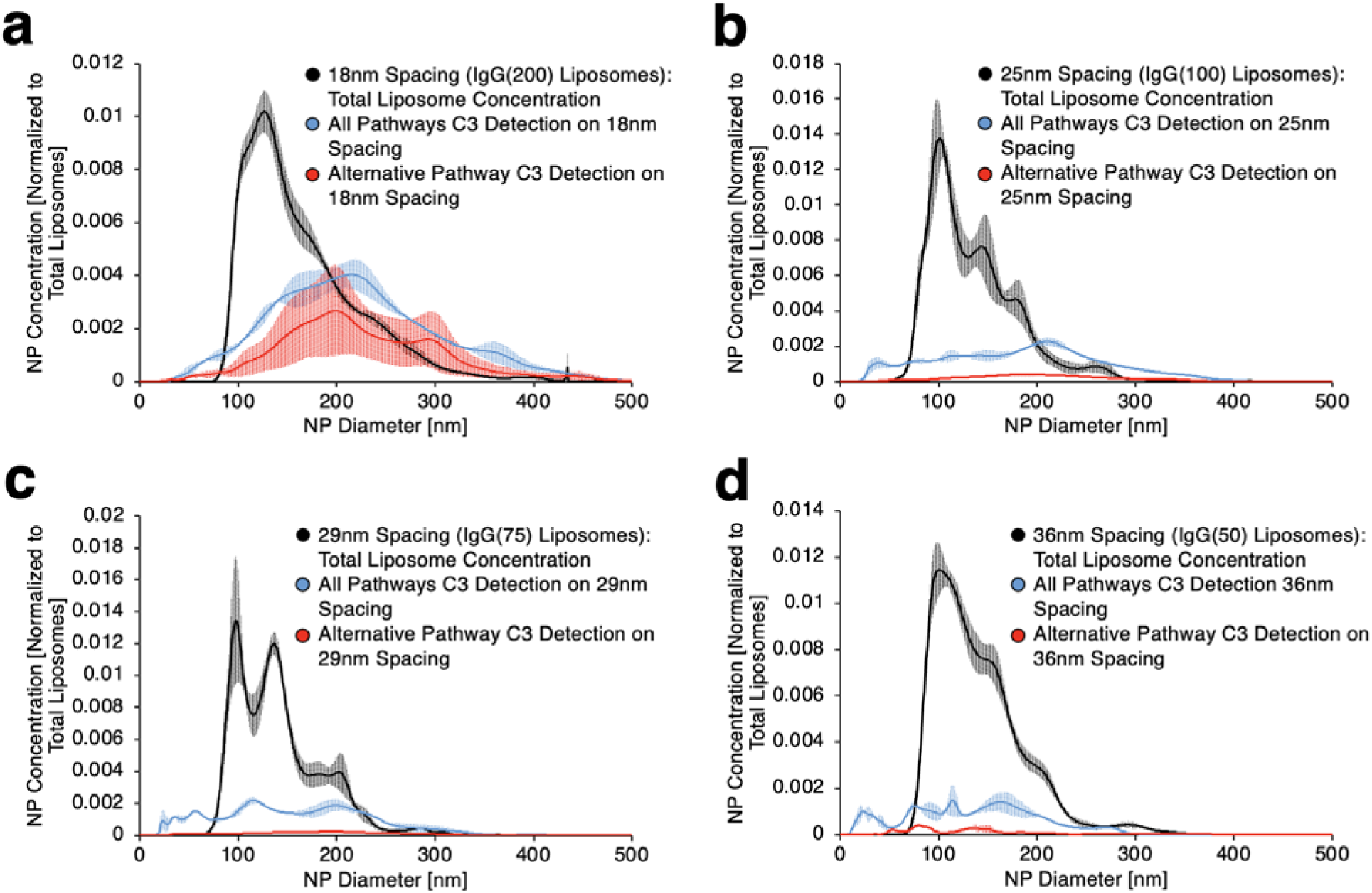
NTA data indicating size vs. concentration distributions of IgG-liposomes with different surface protein densities: (a) 18 nm protein-protein spacing (200 IgG per liposome); (b) 25 nm protein-protein spacing (100 IgG per liposome); (c) 29 nm protein-protein spacing (75 IgG per liposome); (d) 36 nm protein-protein spacing (50 IgG per liposome). Black curves indicate size distributions of each type of nanoparticle before introduction to serum. Blue curves indicate detection of liposomes via fluorescent C3 doped into serum and red curves indicate detection of liposomes via fluorescent C3 doped into serum with gelatin veronal buffer, isolating C3-liposome interactions via the alternative pathway. When isolating the alternative pathway, the number of C3-positive liposomes increases uniquely for the lowest surface protein spacings.

**Supplementary Figure 21.**
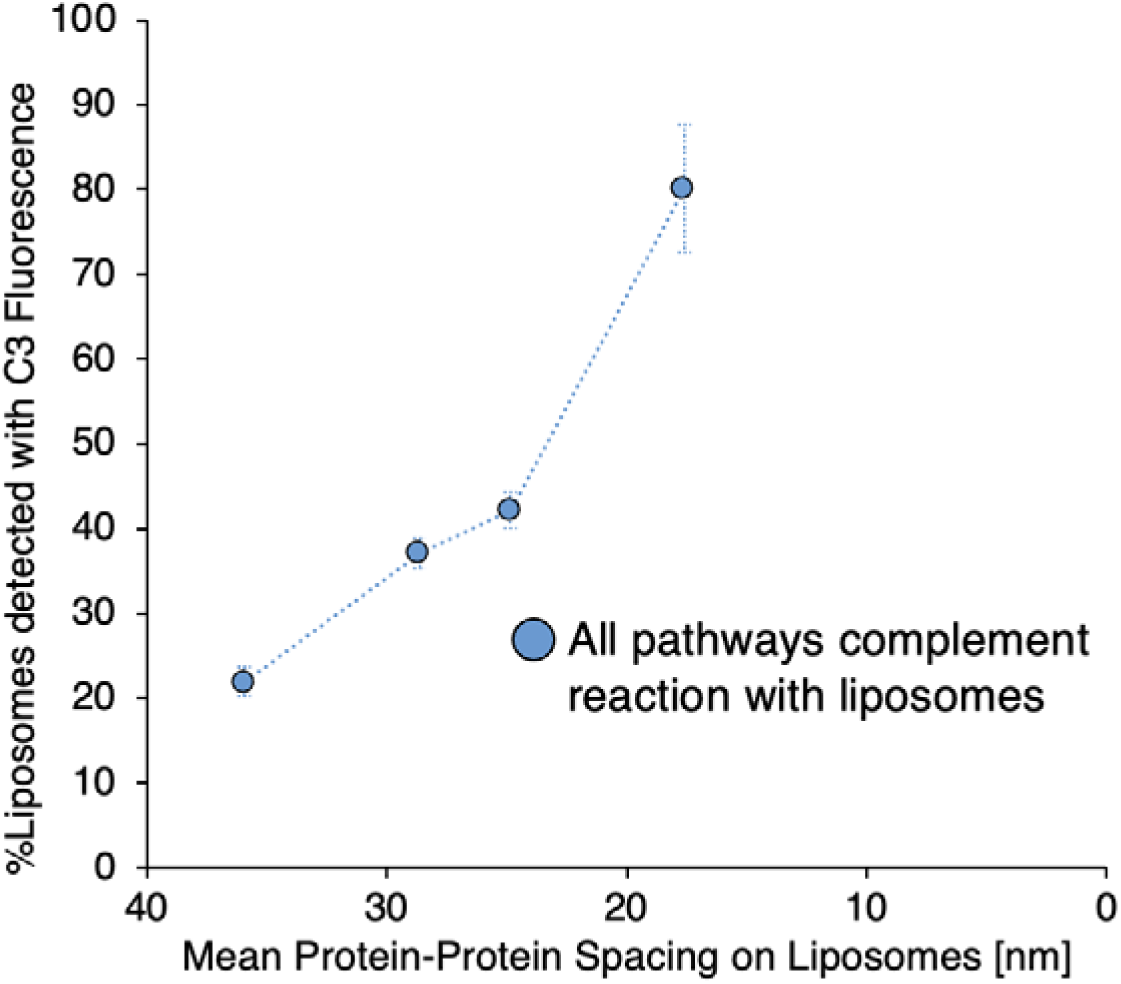
Summary of NTA data on detection of liposomes via interactions with fluorescent C3 in serum. Data shows the fraction of liposomes detected via fluorescent C3 with all complement pathways active, as determined by area under curve for the blue curves in Supplementary Figure 20.

**Supplementary Figure 22.**
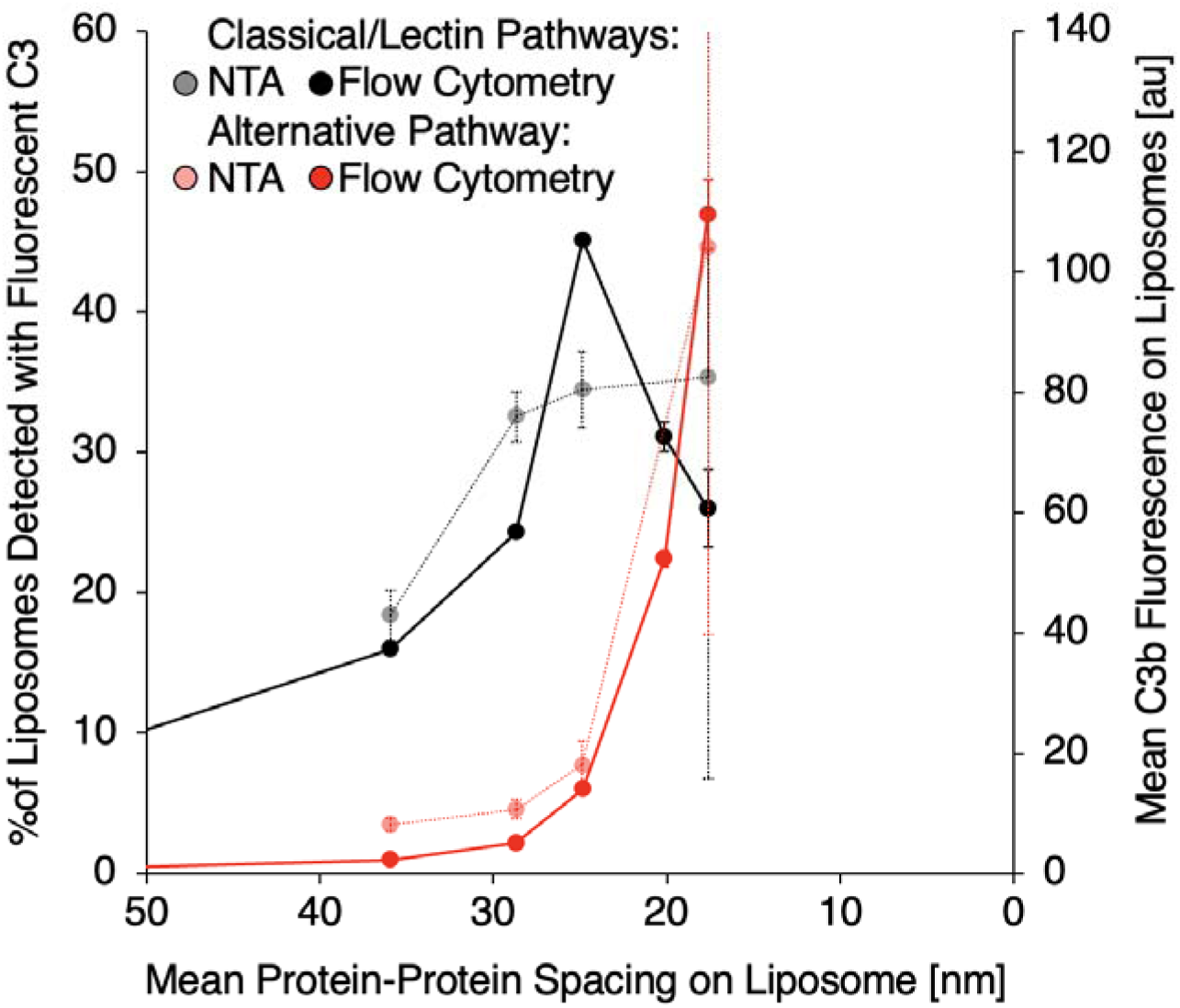
Alternative pathway- and classical/lectin pathway-driven C3b adhesion on liposomes vs. protein-protein spacing on liposome surfaces, as detected by NTA (light-colored curves) and flow cytometry (dark-colored curves). Noting that the flow cytometry data measures a arbitrary units value for quantity of complement on nanoparticles, the vertical axes have been scaled to show that the shape of the C3 deposition vs. surface protein spacing curves match between the two independently-developed assays, especially when isolating the alternative pathway contribution to C3 deposition on the liposomes. Data are reproduced from Figure 2b and Figure 2i in the main text, shown here in overlay to emphasize the agreement between the two measurement techniques.

**Supplementary Figure 23.**
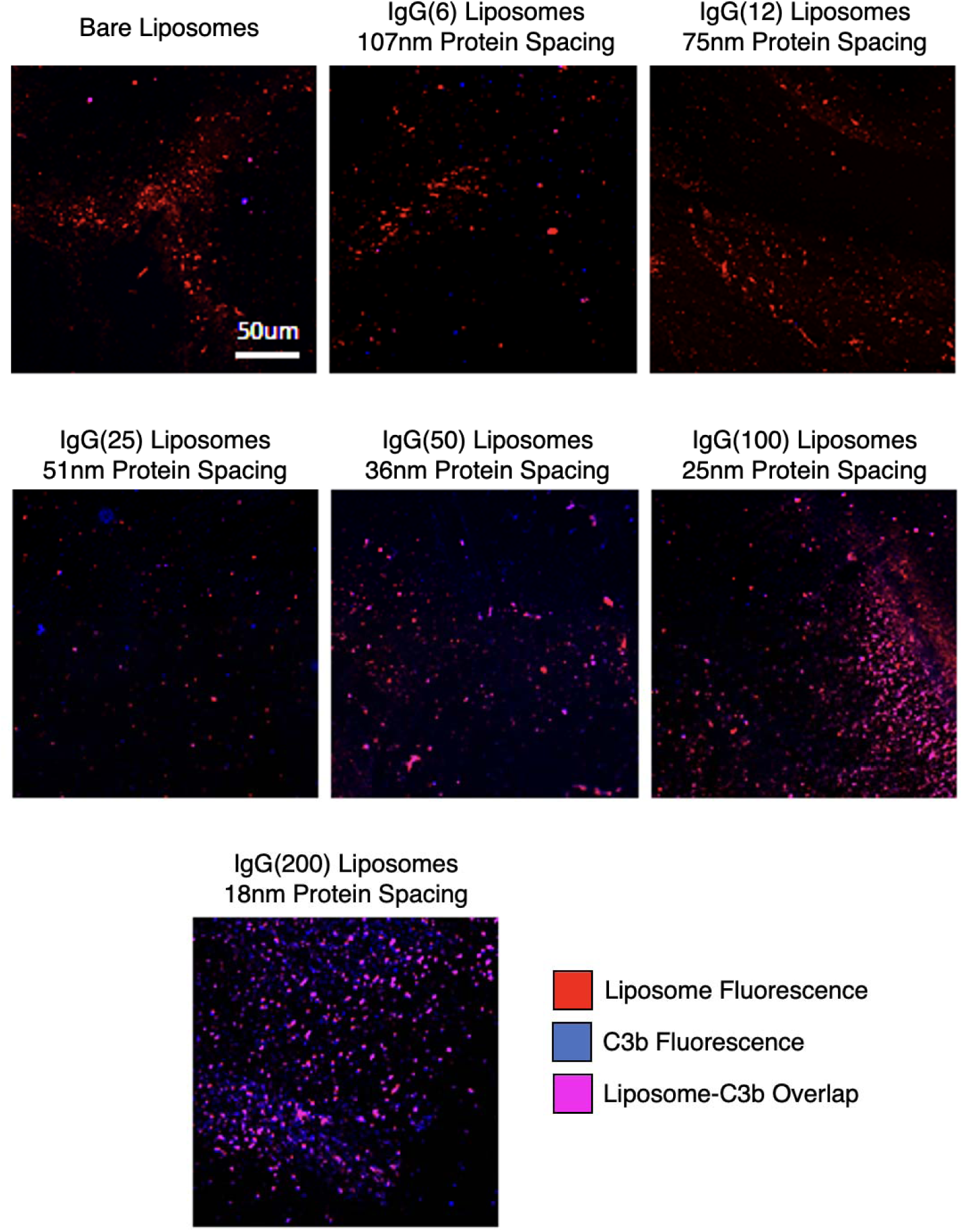
Confocal imaging of C3b deposition on surface-immobilized liposomes as a function of protein-protein spacing on the liposomes. Glass was coated with (3-aminopropyl)triethoxysilane via vapor deposition, followed by azide-terminated PEG-succinimide ester, allowing immobilization of liposomes coated with dibenzocyclooctyne-IgG (DBCO-IgG) via reactions between excess DBCO on the liposome-bound protein and azide groups presented on the functionalized glass surface. Liposomes included fluorescent lipid for tracing their deposition on the surfaces (red pseudocolor in the images). The surface-bound liposomes were exposed to mouse serum and, after washing with PBS, the serum-treated surface-bound liposomes were stained with fluorophore-tagged anti-C3b antibody (blue pseudocolor in the images). The presented images show results for incubation of liposomes with serum treated with gelatin veronal buffer, to isolate the alternative complement pathway.

**Supplementary Figure 24.**
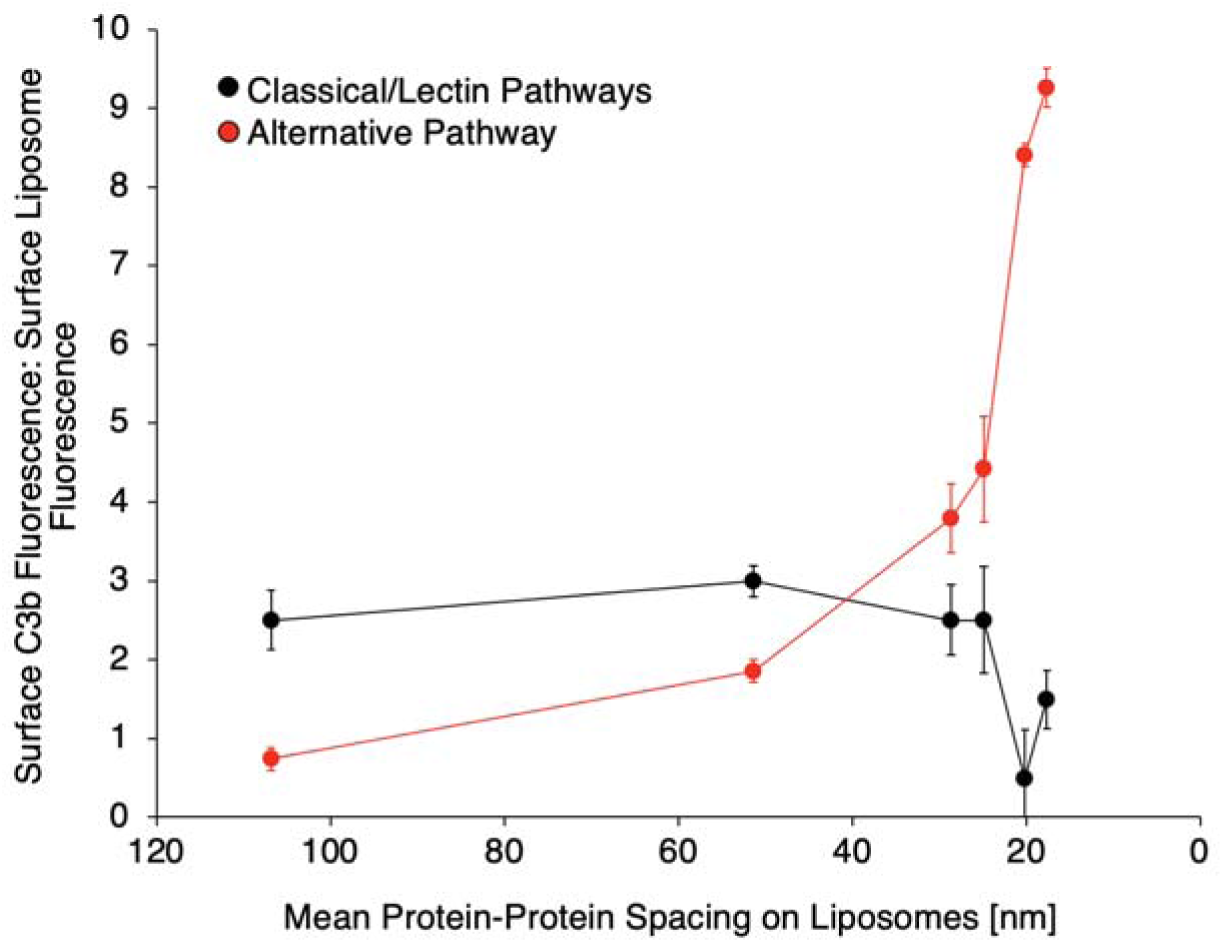
Quantification of C3b deposition on surface-immobilized liposomes as a function of protein-protein spacing on the liposomes. Data are based on confocal imaging, as depicted in Supplementary Figure 23 above. The ratio of C3b stain fluorescence intensity to bound liposome fluorescence intensity was determined in images of three separate glass slides for each condition. Images were obtained for serum treated with gelatin veronal buffer to isolate the alternative complement pathway and with PBS to permit all complement pathways.

**Supplementary Figure 25.**
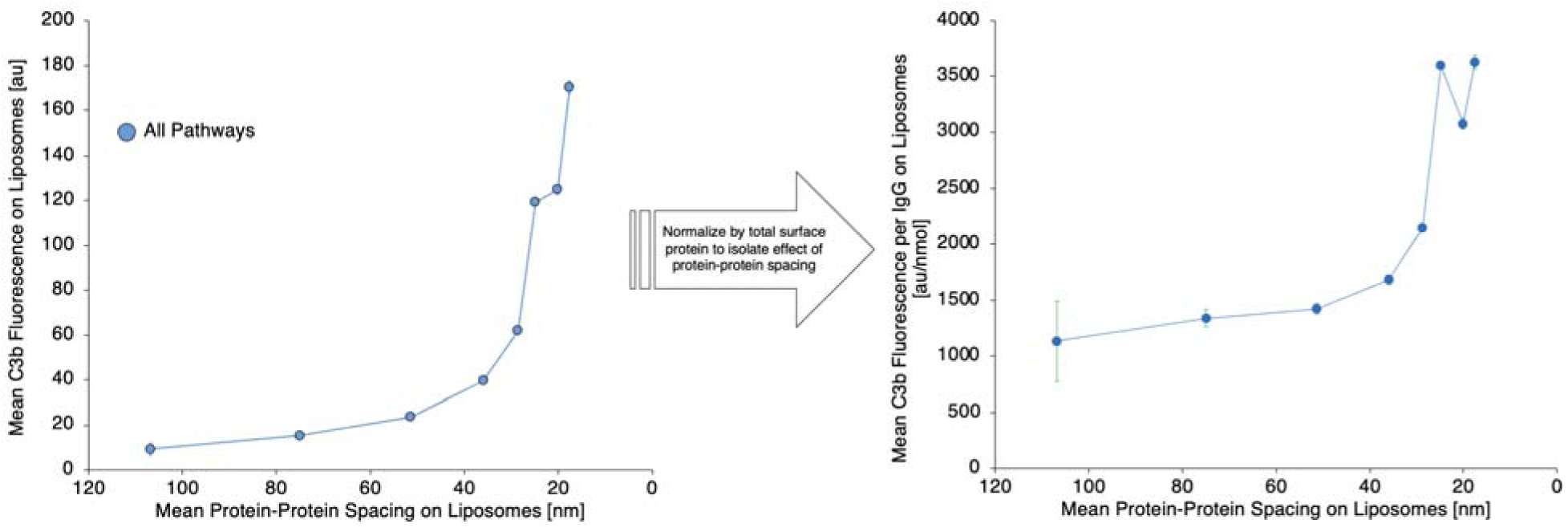
Flow cytometry data indicating fluorescent C3b stain on IgG-liposomes with different IgG-IgG intermolecular spacing after incubation with serum. Data are as in Figure 2b in the main text but show findings for cumulative C3b deposition on nanoparticles by all complement pathways, rather than the isolated alternative pathway or classical+lectin pathways. The data indicate a steep increase in C3b deposition on IgG-liposomes for IgG-IgG spacing < 36 nm that is invariant with normalization to protein concentration. By comparison to Figure 2b, there is a leveling off of “C3b per IgG” below the threshold protein-protein spacing, indicating that alternative pathway-based C3b deposition is the portion of the complement response to the nanoparticle surface that is most sensitive to spacing of surface proteins on the nanoparticle.

**Supplementary Figure 26.**
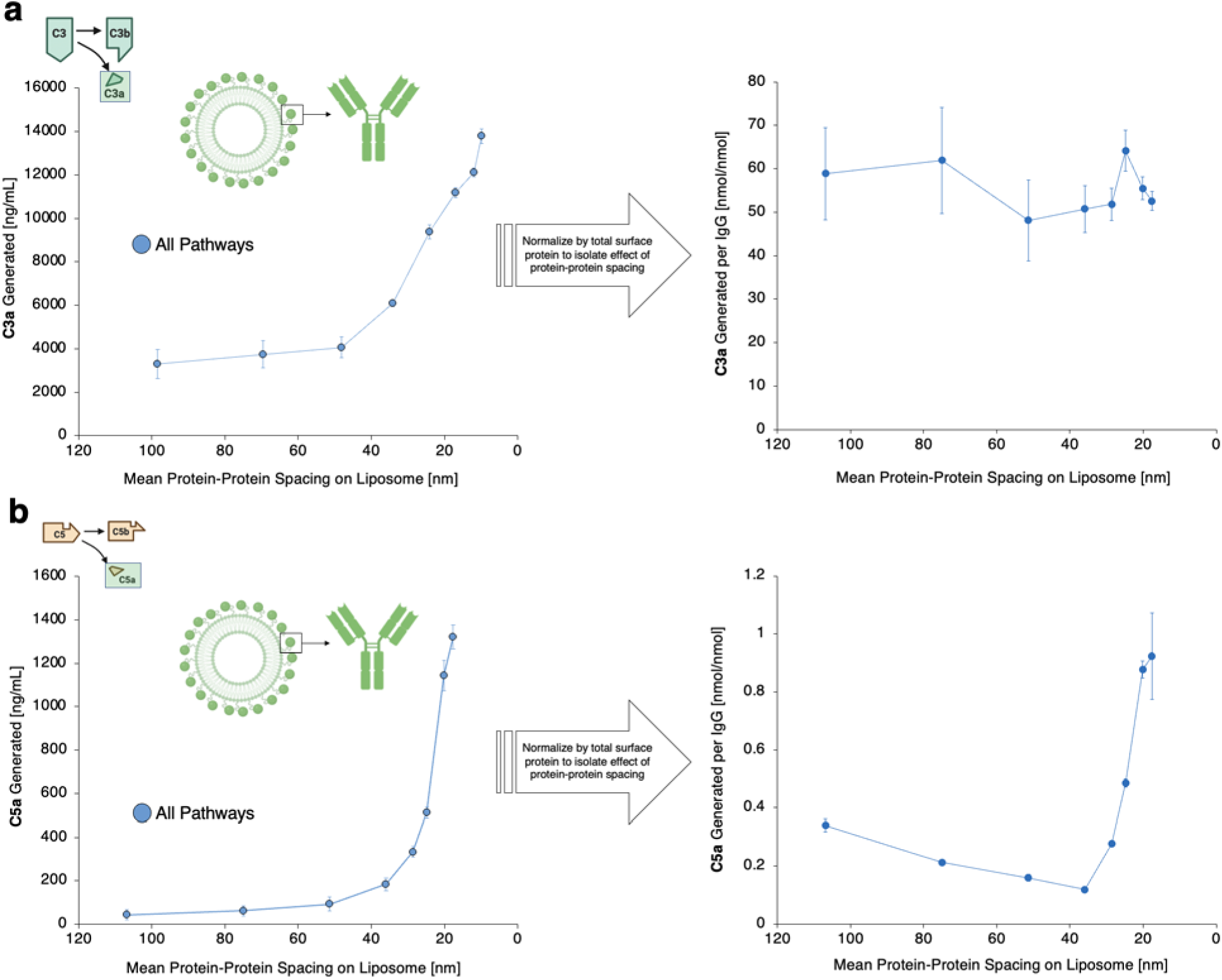
C3a or C5a concentrations in serum treated with IgG-liposomes coated with protein at different intermolecular spacings. Data are as in Figures 3d-e and 3g-h in the main text, but show findings for cumulative C3a generation by all complement pathways, rather than the isolated alternative pathway or classical+lectin pathways. (a) C3a response to IgG-liposomes, showing steeply increasing C3a production for IgG-IgG spacing < 36 nm, but no threshold effect when the data are normalized to protein concentration to obtain “C3a per IgG”. (b) C5a response to IgG-liposomes, showing steeply increasing C5a production for IgG-IgG spacing < 36 nm, with a threshold evident when normalizing to protein concentration to obtain “C5a per IgG”.

**Supplementary Figure 27.**
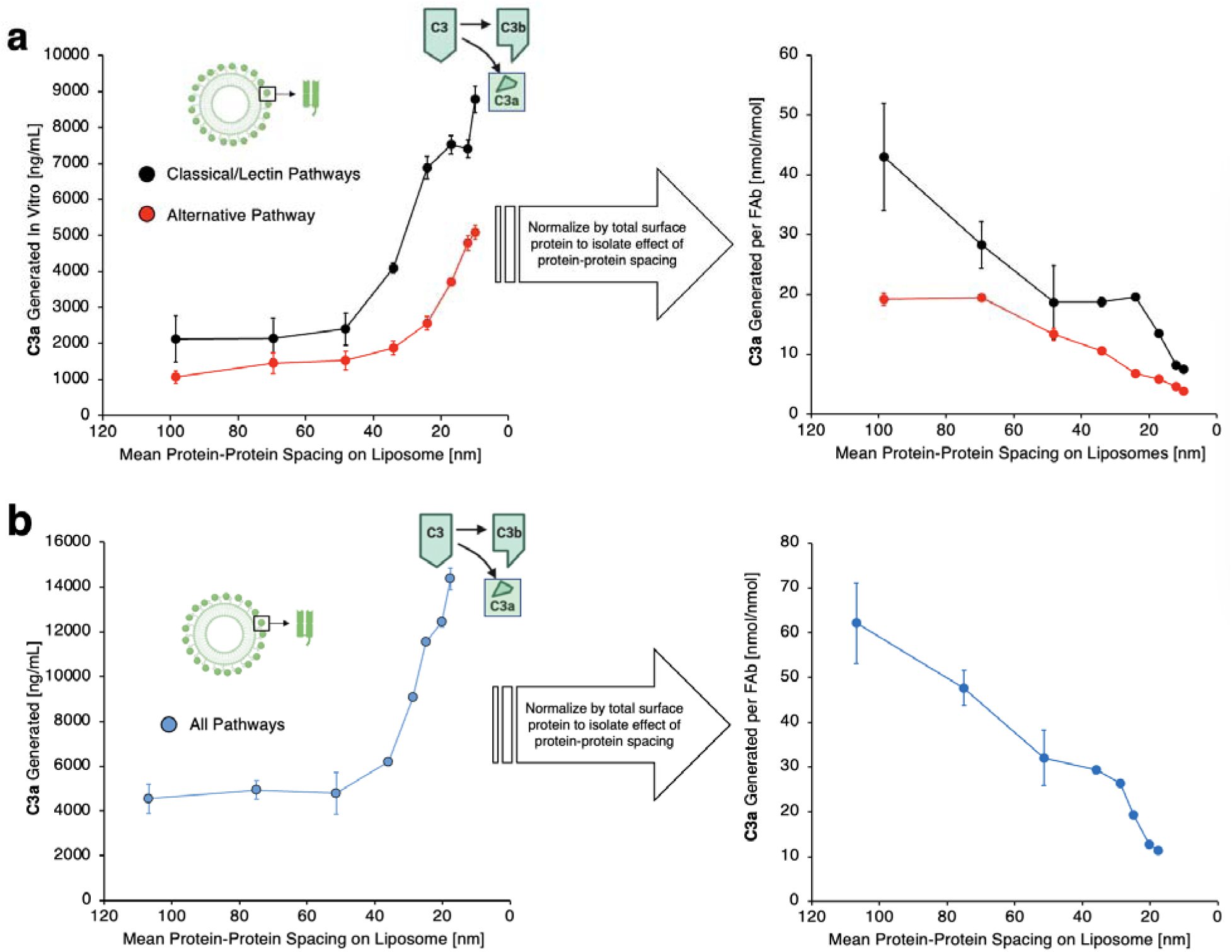
C3a response to FAb-liposomes, showing steeply increasing C3a production for FAb-FAb spacing on liposomes < 36 nm. Data in (a) parallel data in Figure 3d-e in the main text and data in (b) parallel data in Supplementary Figure 26a, but show findings for liposomes conjugated to FAb fragments prepared from IgG, rather than to whole IgG. For the individual complement pathways (a) and the total complement response (b), the C3a generation vs. protein-protein spacing curves for FAb-liposomes resemble those for IgG-liposomes, indicating that the pattern in the data does not depend on the Fc component of the surface-conjugated protein.

**Supplementary Figure 28.**
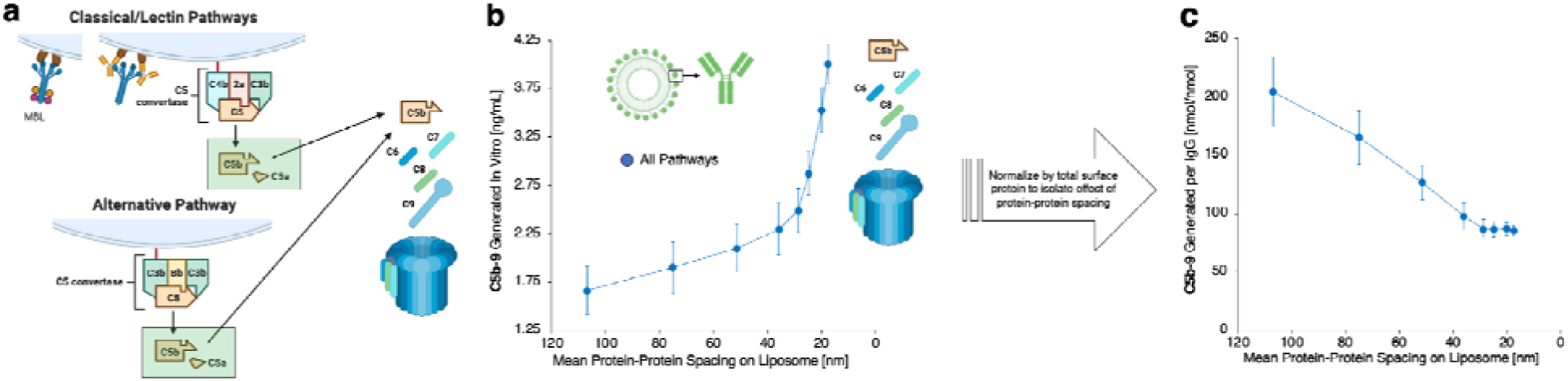
(a) Schematic of C5 convertase and membrane attack complex (MAC) assembly for the different complement pathways. (b) Generation of C5b-9 complex, a soluble marker of MAC formation, by IgG-liposomes in serum as a function of IgG-IgG spacing on the liposomes, showing a steep increase in C5b-9 generation for surface protein spacing < 30 nm. (c) The steep increase in C5b-9 production is not retained for “C5b-9 per IgG,” obtained by normalizing C5b-9 production to total protein on the liposomes.

**Supplementary Figure 29.**
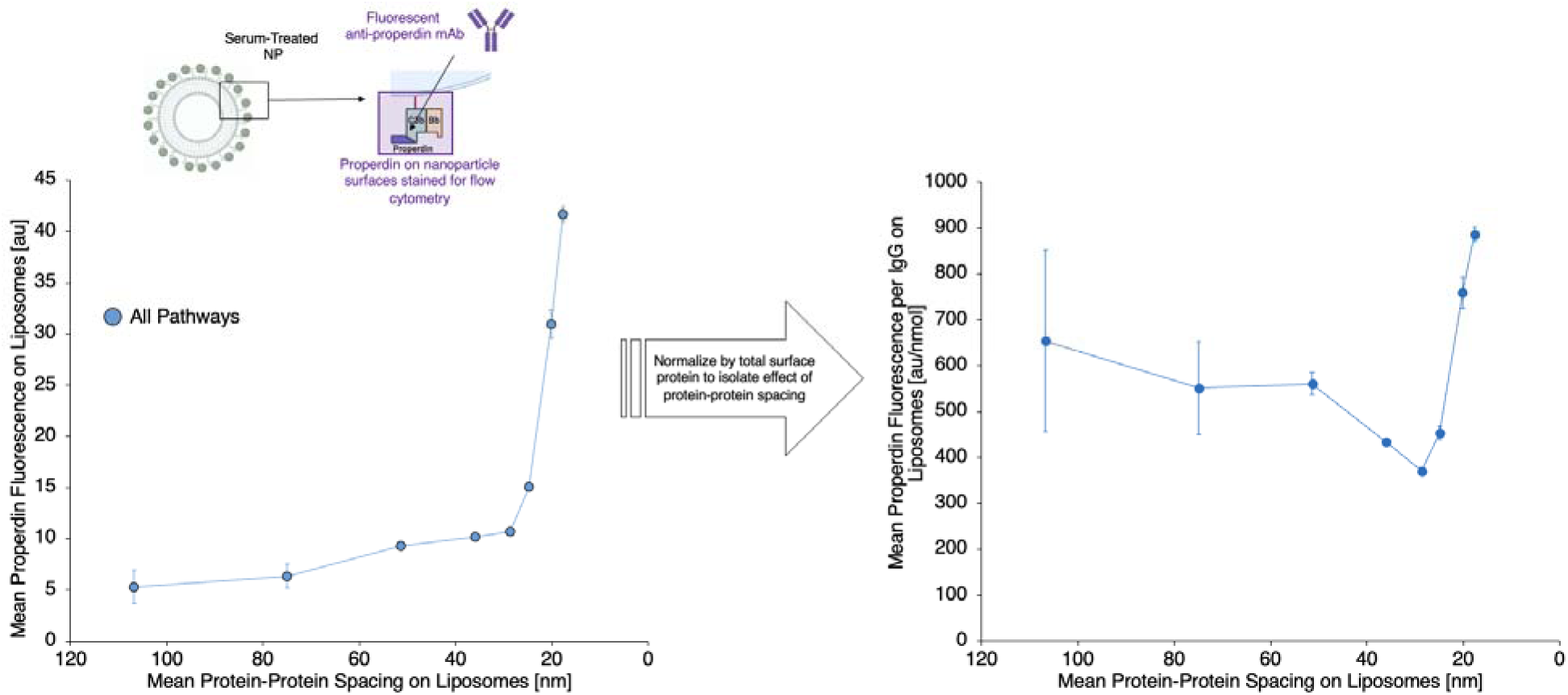
Flow cytometry data summarizing properdin staining on IgG-liposomes with different IgG-IgG intermolecular spacing after liposome-serum incubation. Data match that in Figure 3j-k in the main text but show cumulative properdin deposition on nanoparticles with all complement pathways active, while Figure 3j-k in the main text shows data for the alternative pathway or the classical+lectin pathways in isolation. The data show a threshold closely resembling the data for the alternative pathway alone, since properdin is selectively associated with the alternative pathway. The threshold is unchanged by normalizing properdin deposition to the total number of surface proteins on the liposomes.

**Supplementary Figure 30.**
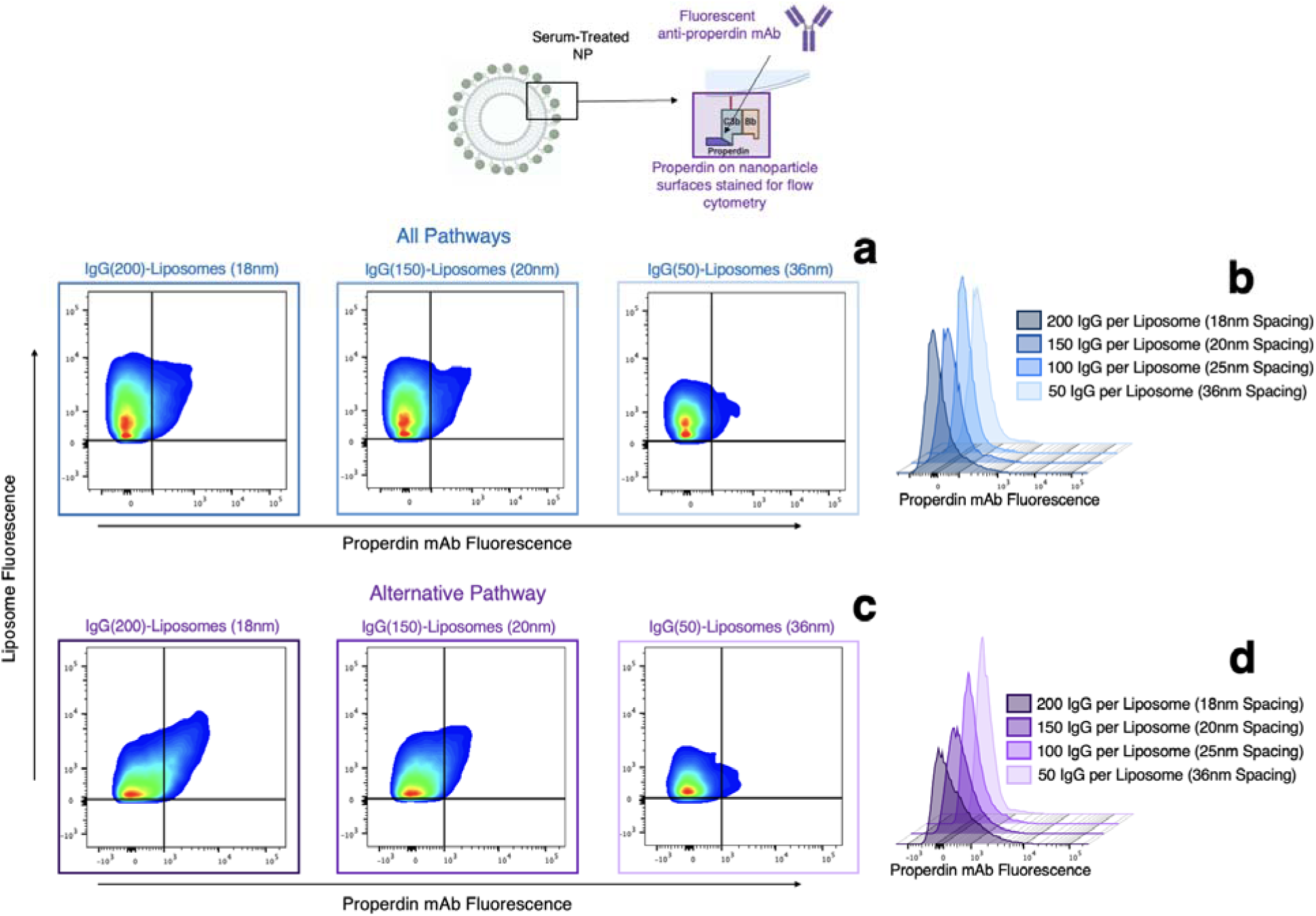
Flow cytometry data describing properdin staining on IgG-liposomes with different surface protein-surface protein intermolecular spacing after incubation with serum. Data show two-dimensional dot plots and histograms indicating the properdin fluorescence distributions that determined the plots in Supplementary Figure 29 and main text Figure 3j-k. Data in (a) and (b) show properdin deposition on liposomes by all complement pathways, as in Supplementary Figure 29. (a) Dot density plots of liposome fluorescence vs. anti-properdin stain fluorescence. (b) Histograms of anti-properdin stain fluorescence on IgG-liposomes with different surface protein-surface protein spacings. Alternative pathway properdin deposition (c-d) closely resembles properdin deposition due to all complement pathways (alternative+classical+lectin), since properdin deposition is uniquely associated with the alternative pathway.

**Supplementary Figure 31.**
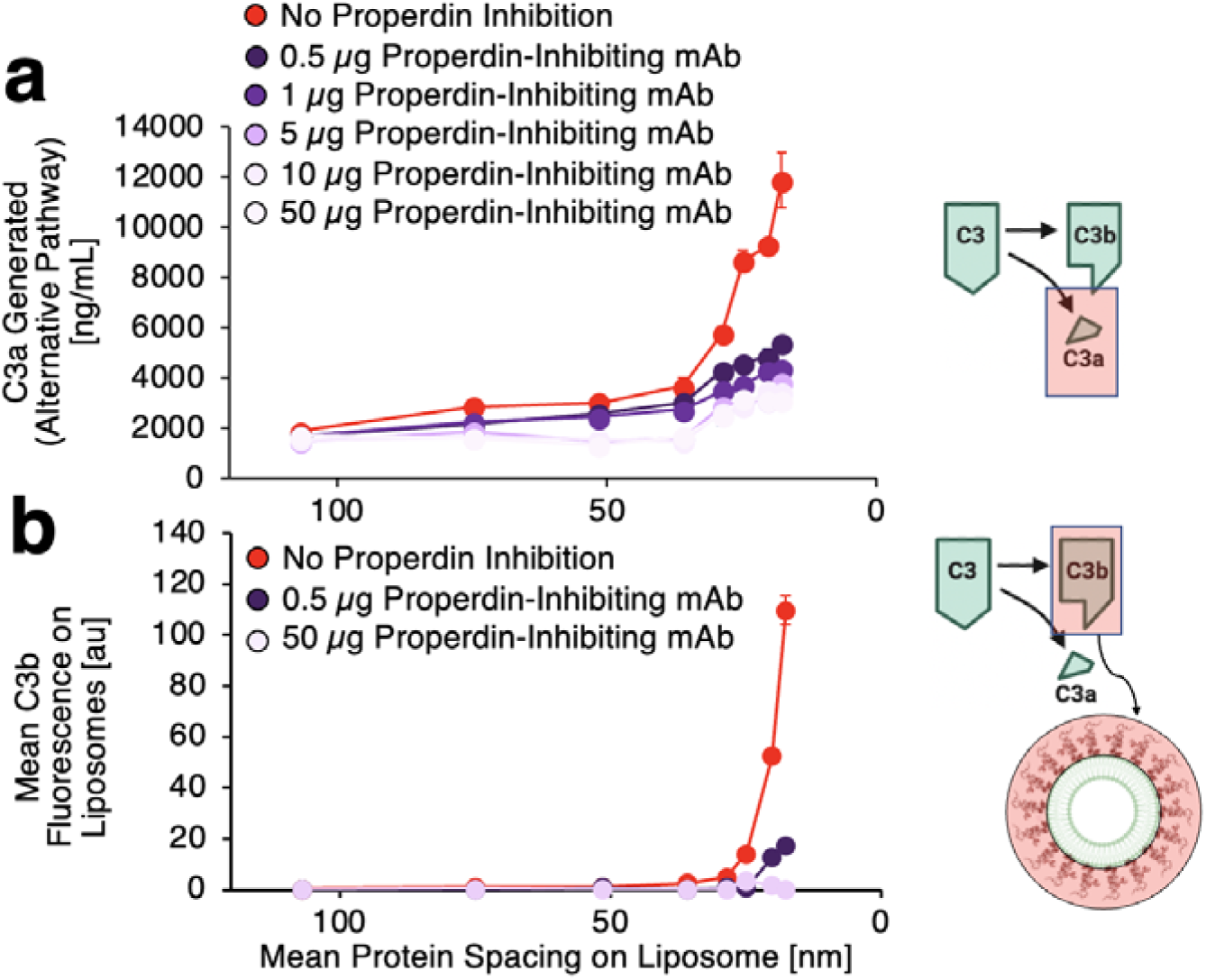
(a) Alternative pathway C3a production in serum+protein nanoparticle samples vs. protein-protein spacing on the nanoparticles, with inhibition of properdin eliminating the spacing-dependent changes in C3a production. (b) Alternative pathway C3b fluorescence on protein-nanoparticles vs. protein-protein spacing on the nanoparticles, as in Figure 2b in the main text, with inhibition of properdin eliminating the spacing-dependent changes in C3b deposition.

**Supplementary Figure 32.**
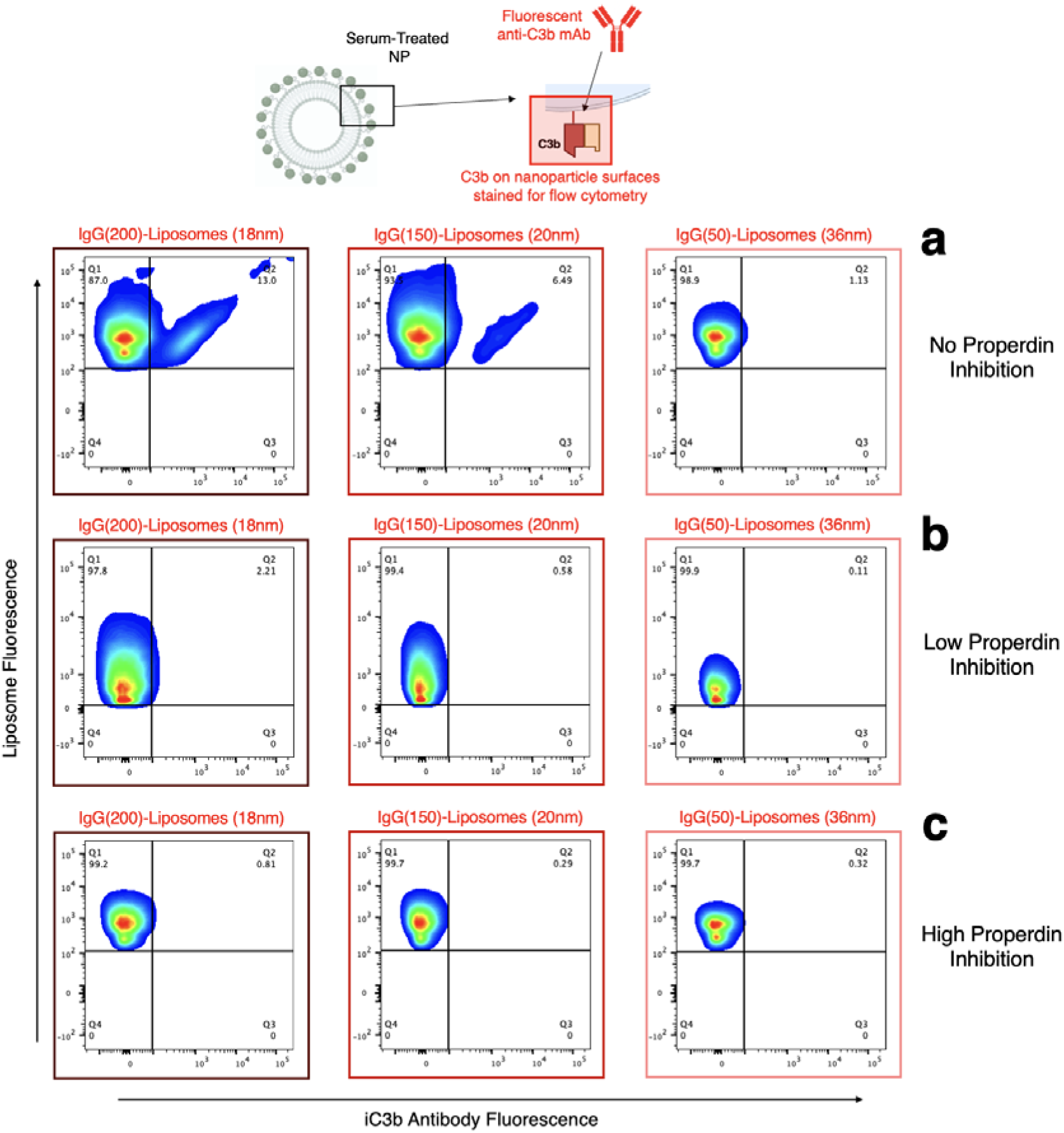
Flow cytometry data indicating fluorescent C3b stain on IgG-liposomes with different surface protein-protein intermolecular spacing after incubation with serum pre-treated with different quantities of properdin-inhibiting antibody. (a) Data for liposomes in serum without properdin-inhibiting antibody, as also depicted in Supplementary Figure 17c and Figure 2c in the main text. (b) Data for liposomes in serum treated with 0.5 µg of properdin-inhibiting antibody, showing that properdin inhibition causes loss of C3b-high populations, even for liposomes with the smallest protein-protein spacing. (c) Data as in (b), but with serum treated with 50 µg of properdin-inhibiting antibody. As with (b), C3b-high populations are missing. All data reflect serum treated with gelatin veronal buffer, isolating the alternative pathway of complement. Findings from these experiments are summarized in Supplementary Figure 31 above.

**Supplementary Figure 33.**
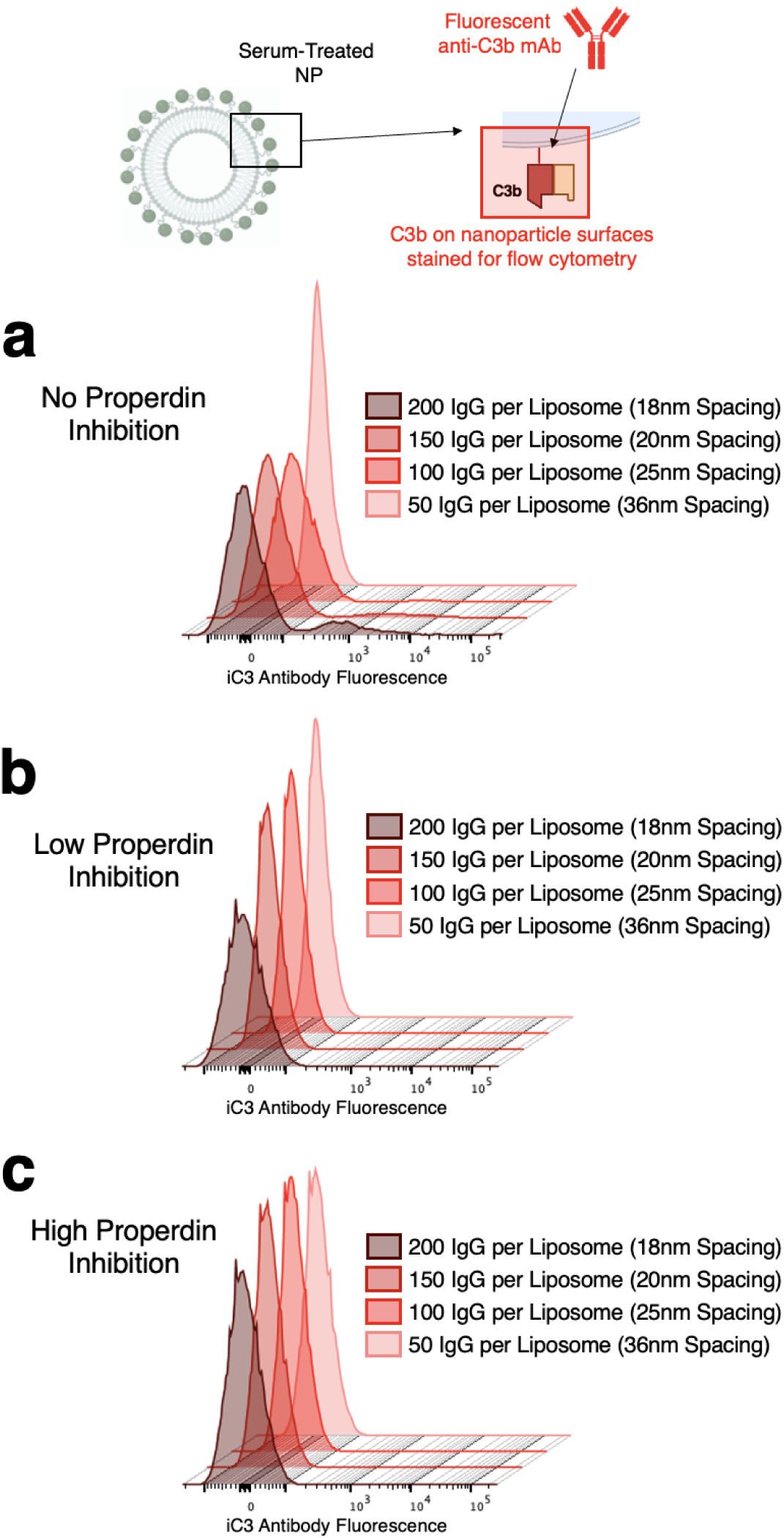
An additional representation of data from Supplementary Figures 31-32, showing histograms of anti-C3b fluorescence on IgG-liposomes after incubation with serum as a function of IgG-IgG intermolecular spacing on the liposomes. As with data in Supplementary Figure 32, high-C3b populations are evident without properdin-inhibiting antibody (a) but are not evident with either 0.5 µg properdin-inhibiting antibody (b), or 50 µg properdin-inhibiting antibody (c). Findings from these experiments are summarized in Figure 3h in the main text.

**Supplementary Figure 34.**
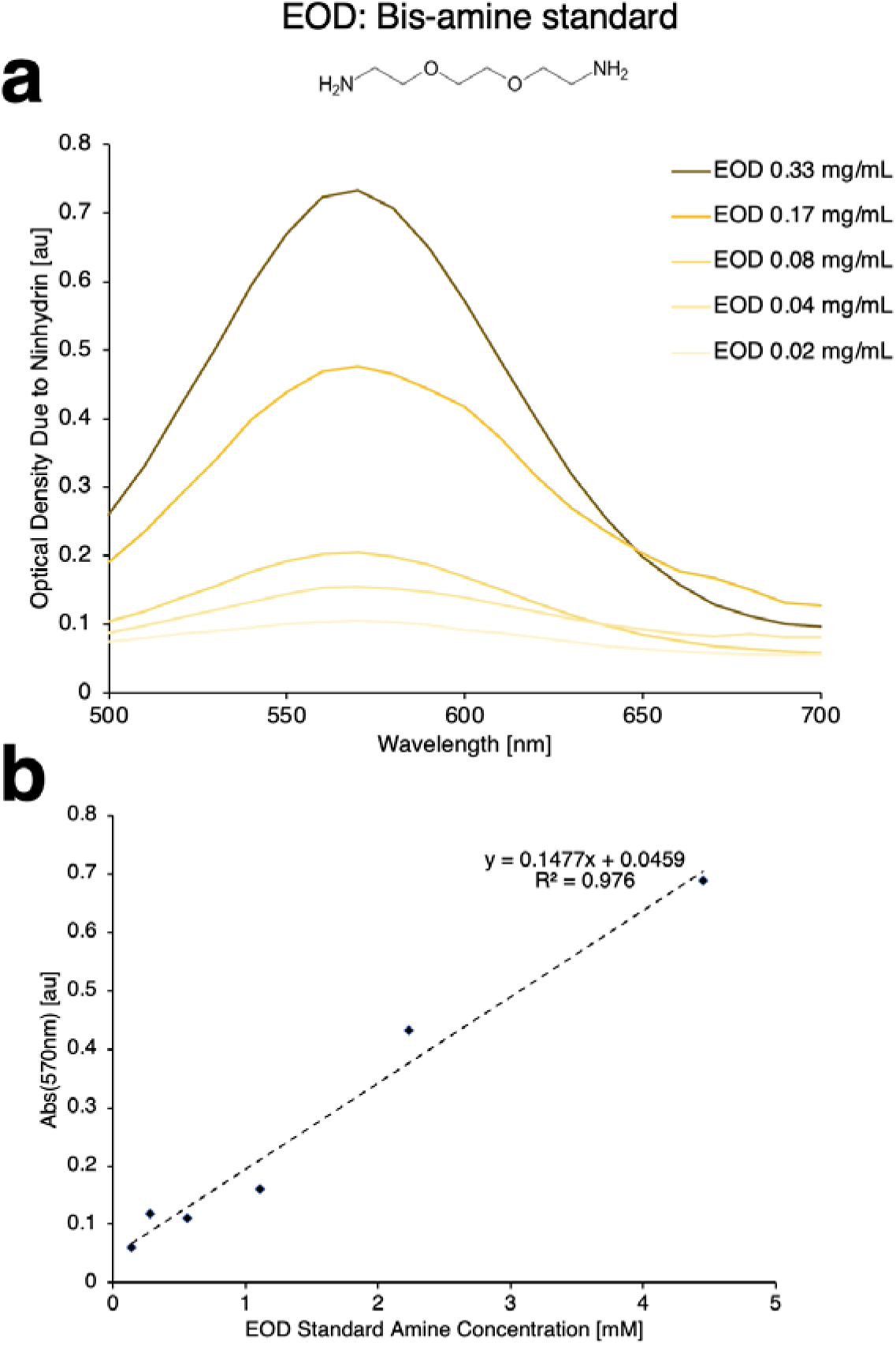
UV/visible spectrophotometry data showing results of preparing bis-amine standards (2-2’-(ethylenedioxy)bis(ethylamine), EOD) with ninhydrin solutions. (a) Optical absorbance spectra showing peak in absorbance at ∼570 nm, corresponding to Ruhmann’s purple, as a function of quantity of bis-amine standard added to ninhydrin preparations. (b) Magnitude of optical absorbance at 570 nm as a function of quantity of bis-amine standard added to ninhydrin preparations, demonstrating linear range of the assay. For each repetition of the assay with samples containing unknown quantities of amines, a new EOD bis-amine standard curve was prepared in parallel to identify linear range and quantify amine content in samples.

**Supplementary Figure 35.**
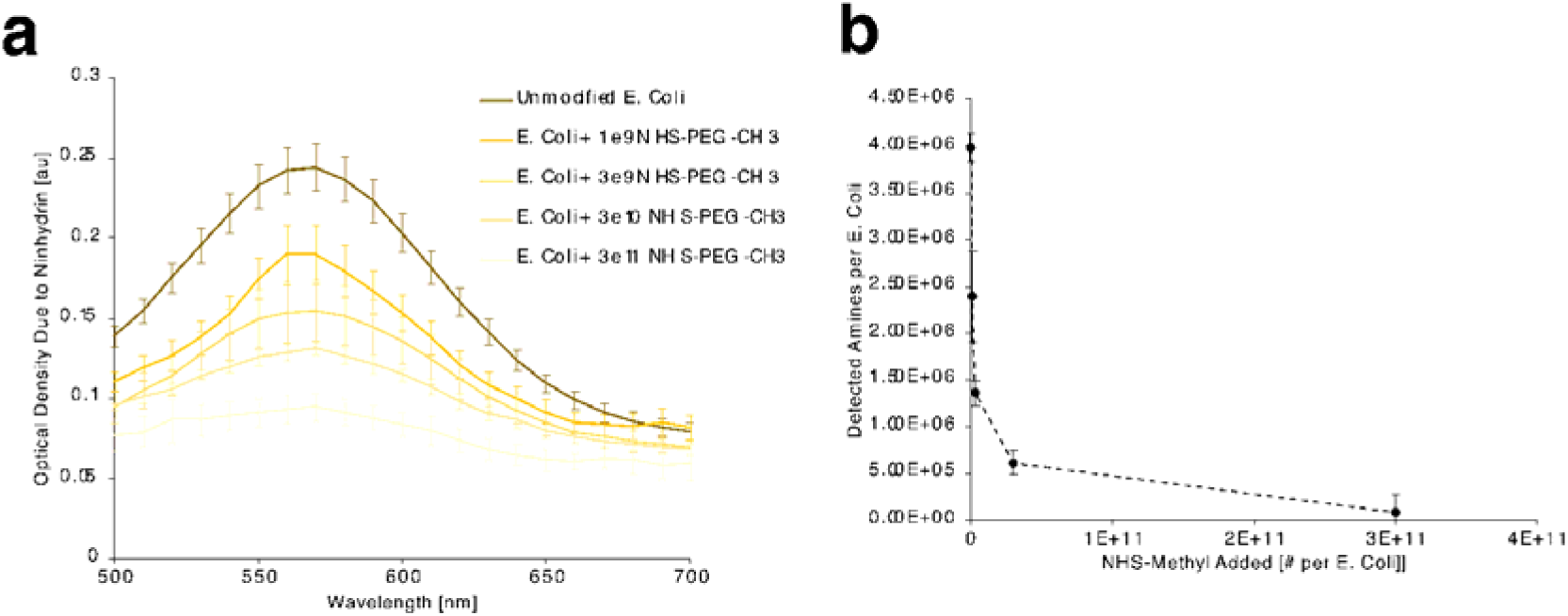
UV/visible spectrophotometry data as in Supplementary Figure 31, but showing results of ninhydrin assays with methyl-PEG-NHS treated E. coli. Unmodified E. coli were compared to E. coli treated with different quantities of methyl-PEG-NHS for generation of Ruhmann’s purple in ninhydrin/ethanol solutions, as identified by optical absorbance peaks at 570 nm (a). 570 nm absorbance values were compared to standard curves as in Supplementary Figure 34b to determine the concentration of amines in the prepared samples. Concentrations of E. coli in the suspensions were determined prior to the assay by measuring optical density at 600 nm, allowing conversion of amine concentrations measured in the ninhydrin assay to values indicating number of amines per E. coli vs. methyl-PEG-NHS added per E. coli (b).

**Supplementary Figure 36.**
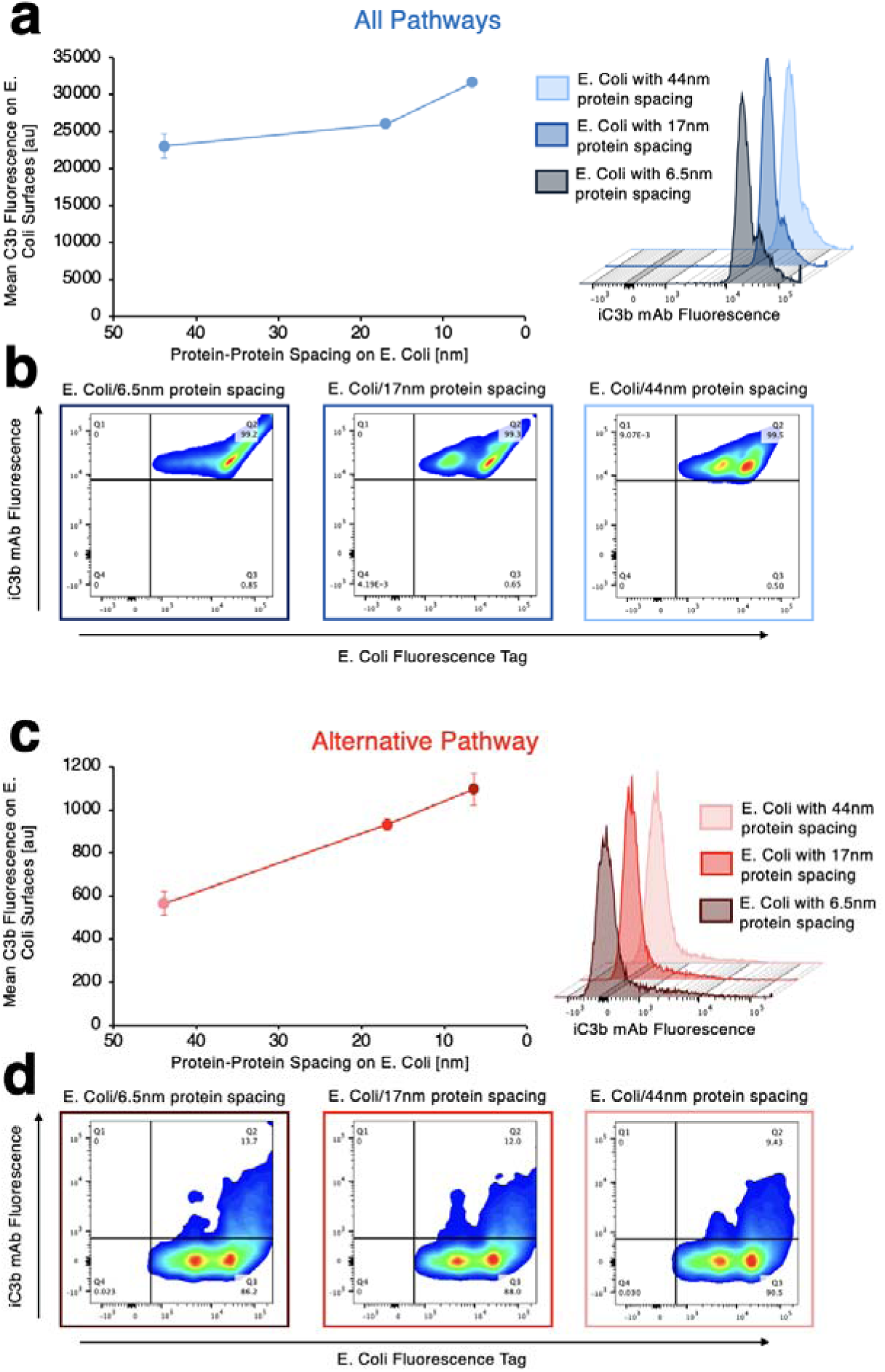
Flow cytometry data showing C3b staining on E. coli after incubation with serum, either with all complement pathways active (a-b) or with the alternative pathway isolated by inclusion of gelatin veronal buffer with the serum (c-d). (a) and (c) depict mean fluorescent intensity of C3b stain (leftmost panels) as a function of effective protein-protein spacing on the E. coli, as modified by pre-treatment of the E. coli with methyl-PEG-NHS and quantified by ninhydrin assay, depicted in Supplementary Figures 34-35. (a) and (c) also show histograms of C3b fluorescence on E. coli for treatment with different quantities of methyl-PEG-NHS. (b) and (d) show two-dimensional dot plots of C3b fluorescence vs. fluorescence from an Alexafluor 488 tag applied ubiquitously to identify the E. coli, showing the gating strategy for identification of C3b signal on E. coli. The isolated alternative pathway yields over an order of magnitude less C3b adhesion to the E. coli than the combined alternative+classical+lectin pathways, likely due to the outsized role of the lectin pathway in responses to bacteria. Nonetheless, in all results, C3b deposition increases as effective protein-protein spacing decreases.

**Supplementary Figure 37.**
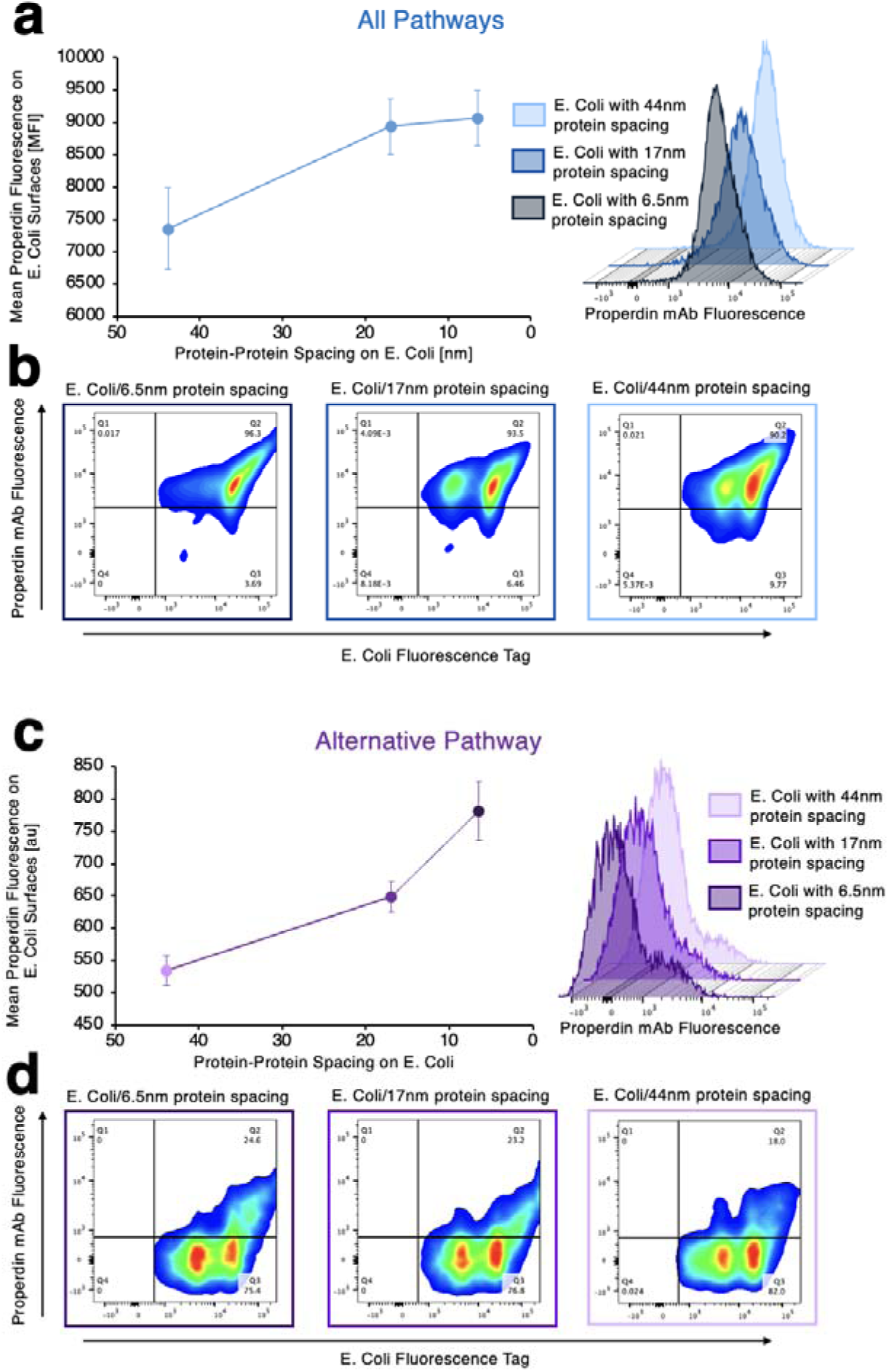
Flow cytometry data showing properdin staining on E. coli after incubation with serum, either with all complement pathways active (a-b) or with the alternative pathway isolated by inclusion of gelatin veronal buffer with the serum (c-d). Data are arranged as for C3b staining data in Supplementary Figure 36, with (a) and (c) showing mean properdin fluorescence data and properdin fluorescence histograms and (b) and (d) showing two dimensional plots of properdin fluorescence vs. ubiquitous tag fluorescence on E. coli. The patterns in fluorescence with respect to protein-protein spacing resemble those for C3b staining in Supplementary Figure 36.

**Supplementary Figure 38.**
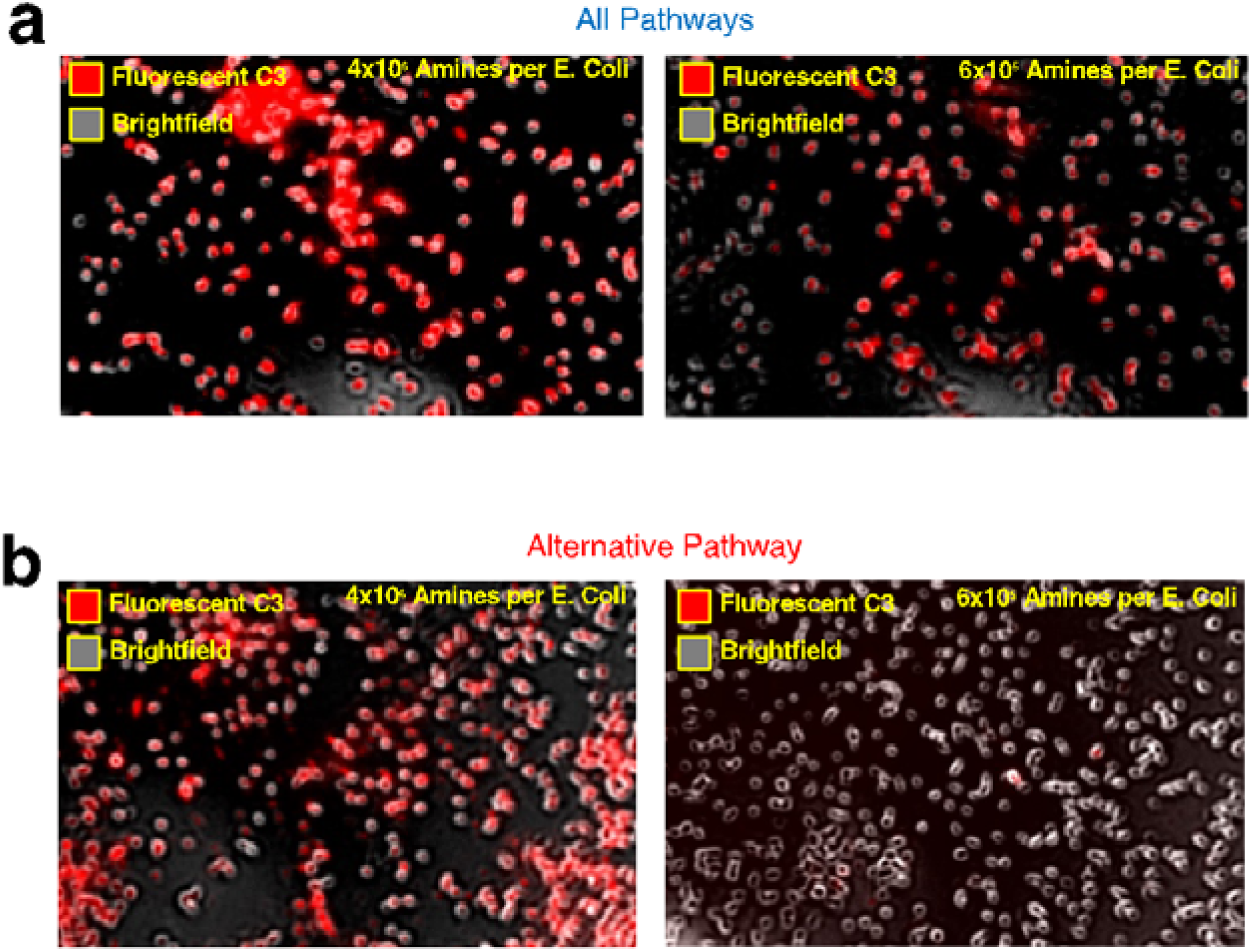
Micrographs showing fluorescent C3 accumulation on E. coli during incubation with serum, either with all complement pathways active (a) or with the alternative pathway isolated via gelatin veronal buffer (b). Leftmost panels depict C3 accumulation on E. coli with 4×10^6^ amines per bacterium (as detected by quantitative ninhydrin assay) and rightmost panels depict C3 accumulation on E. coli with 6×10^5^ amines.

**Supplementary Figure 39.**
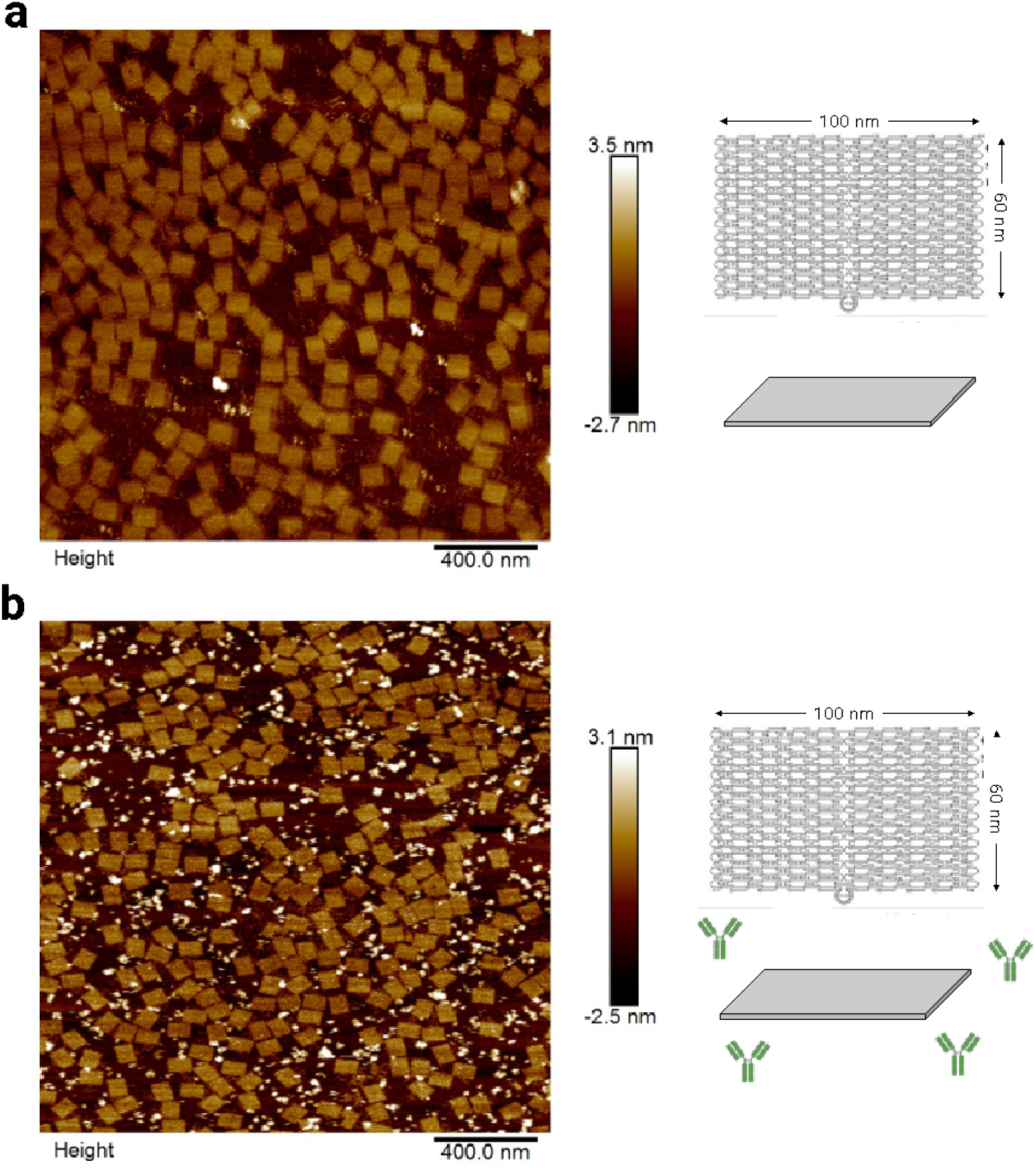
Atomic force micrographs of planar DNA origami (a) before and (b) after addition of IgG to the origami surfaces. Data depict DNA origami without azide-terminated anchor strands to react with strained alkyne (DBCO) conjugated IgG. A portion of panel (b) is depicted in Figure 5a in the main text. (b) shows that IgG does not non-specifically adhere to the DNA origami without an azide anchor strand.

**Supplementary Figure 40.**
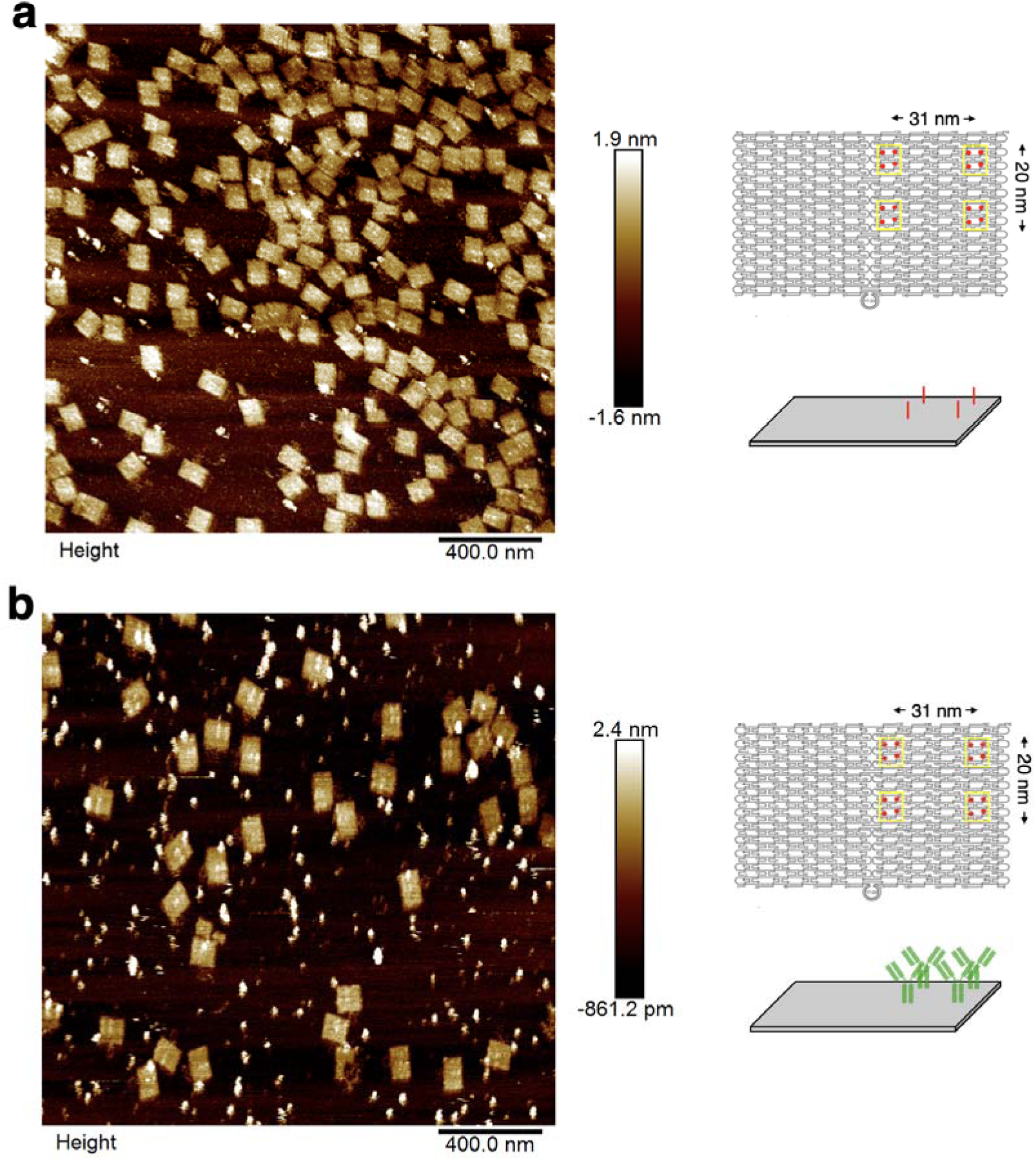
Atomic force micrographs of planar DNA origami (a) before and (b) after addition of IgG to the origami surfaces. Data depict DNA origami with four azide-terminated anchor strands arranged in a 31 nm × 20 nm quadrangle. A portion of panel (b) is depicted in Figure 5b in the main text. (b) shows that IgG adheres at the positions of the azide-terminated anchor strands on the origami, as depicted in (a).

**Supplementary Figure 41.**
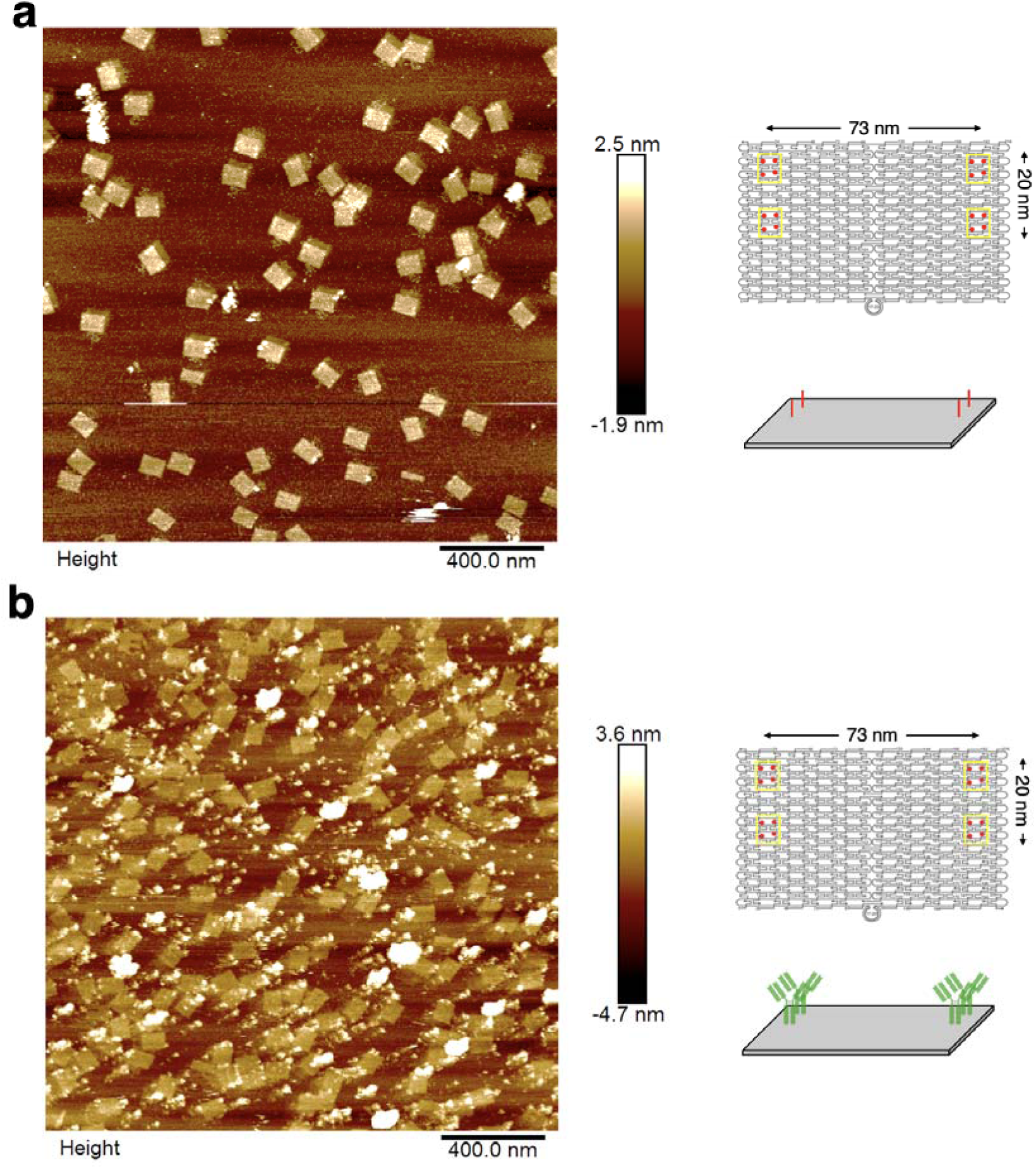
Atomic force micrographs of planar DNA origami (a) before and (b) after addition of IgG to the origami surfaces. Data depict DNA origami with four azide-terminated anchor strands arranged in a 73 nm × 20 nm quadrangle. A portion of panel (b) is depicted in Figure 5c in the main text. (b) shows that IgG adheres at the positions of the azide-terminated anchor strands on the origami, as depicted in (a).

**Supplementary Figure 42.**
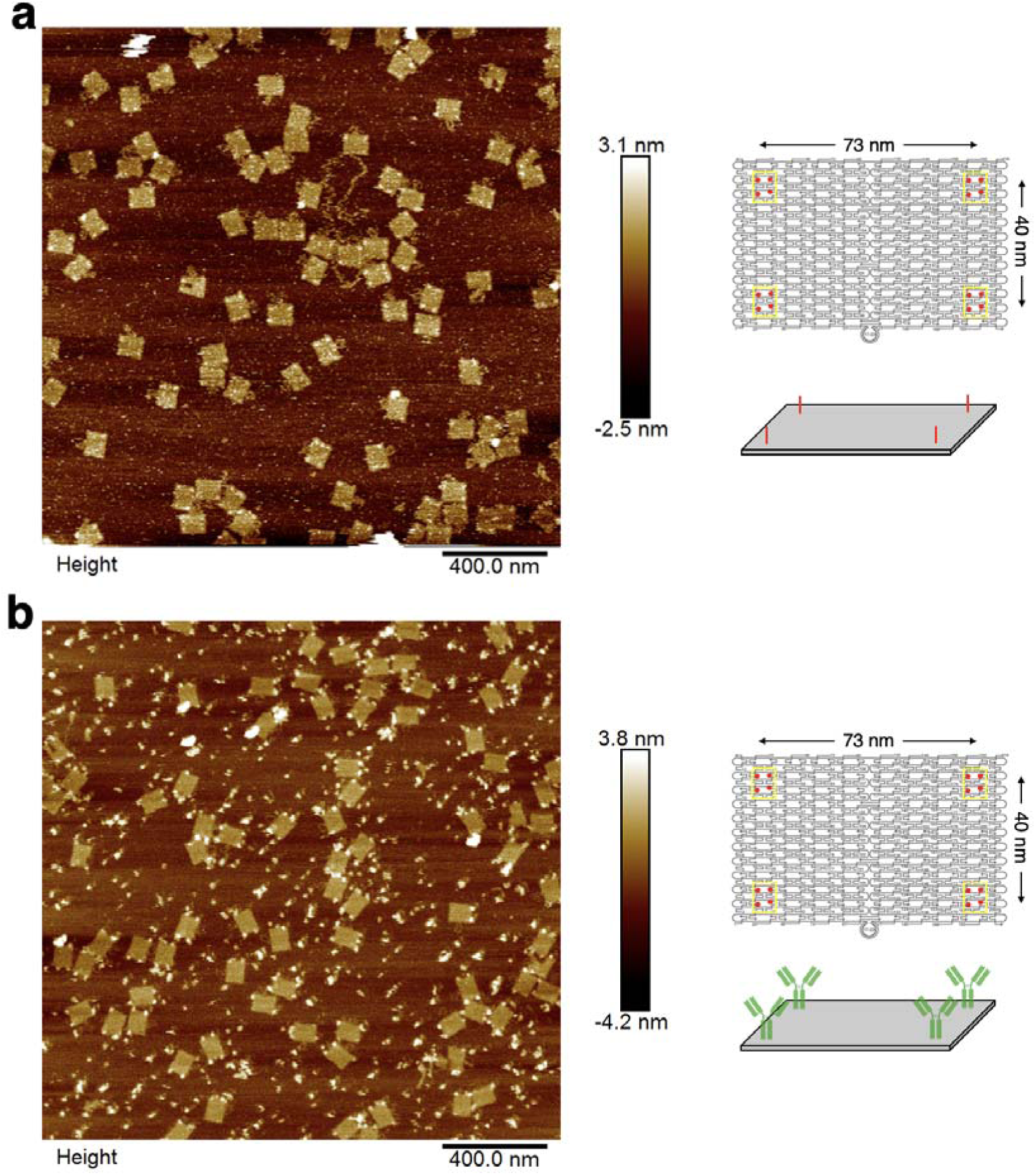
Atomic force micrographs of planar DNA origami (a) before and (b) after addition of IgG to the origami surfaces. Data depict DNA origami with four azide-terminated anchor strands arranged in a 73 nm × 40 nm quadrangle. A portion of panel (b) is depicted in Figure 5d in the main text. (b) shows that IgG adheres at the positions of the azide-terminated anchor strands on the origami, as depicted in (a).

**Supplementary Figure 43.**
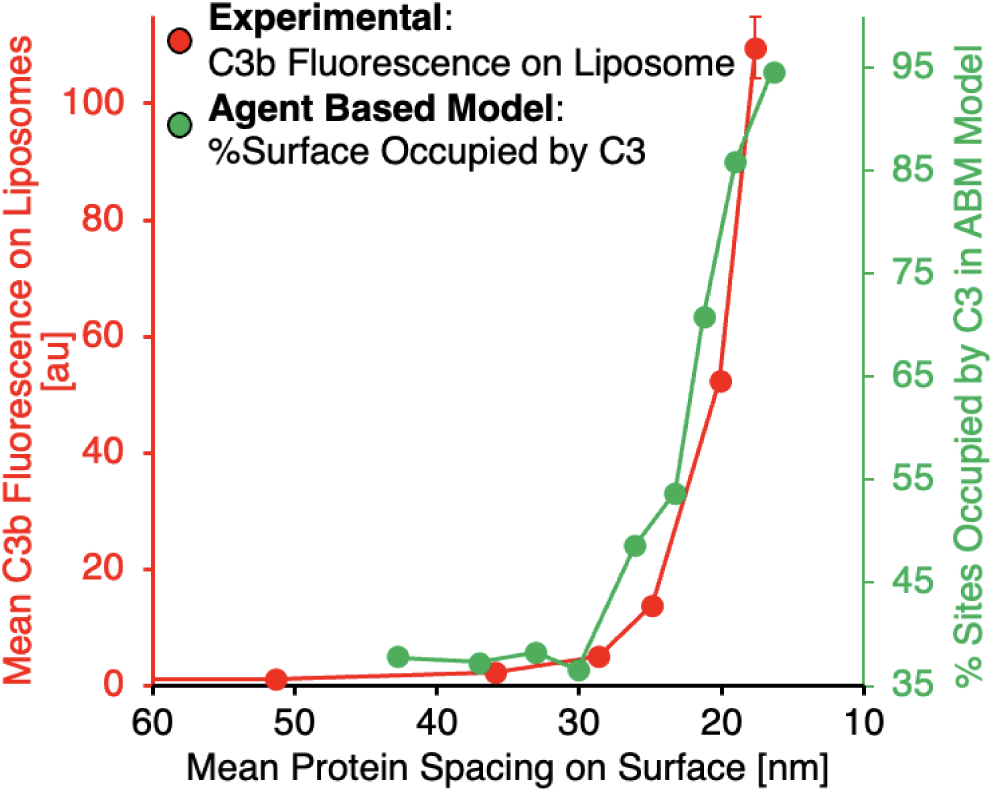
Alternative depiction of the data in Figure 6f in the main text. The main text presents agent based model outcomes as number of C3b adhered per nanoparticle, while the data here converts that finding to percentage of available C3b-nucleophile binding sites occupied in the model outcome. The normalized data presented here maintains the curve shape found in the experimental data, preserving the C3 adhesion vs. binding site spacing threshold noted in other data throughout the text and supplement.

### SUPPLEMENTARY SECTION B: Descriptions of Supplementary Movies

**Supplementary Movie 1.** Scattering nanoparticle tracking analysis recording for liposomes in buffer

**Supplementary Movie 2.** Fluorescence nanoparticle tracking analysis recording for liposomes in buffer

**Supplementary Movie 3.** Scattering nanoparticle tracking analysis recording for plasma with fluorescent C3

**Supplementary Movie 4.** Fluorescence nanoparticle tracking analysis recording for plasma with fluorescent C3

**Supplementary Movie 5.** Fluorescence nanoparticle tracking analysis recording for plasma with fluorescent C3 and protein-liposomes with surface protein spacing = 18 nm

**Supplementary Movie 6.** Fluorescence nanoparticle tracking analysis recording for plasma with fluorescent C3 and protein-liposomes with surface protein spacing = 25 nm

**Supplementary Movie 7.** Fluorescence nanoparticle tracking analysis recording for plasma with fluorescent C3 and protein-liposomes with surface protein spacing = 29 nm

**Supplementary Movie 8.** Fluorescence nanoparticle tracking analysis recording for plasma with fluorescent C3 and protein-liposomes with surface protein spacing = 36 nm

**Supplementary Movie 9.** Fluorescence nanoparticle tracking analysis recording for plasma/gelatin veronal buffer with fluorescent C3 and protein-liposomes with surface protein spacing = 18 nm

**Supplementary Movie 10.** Fluorescence nanoparticle tracking analysis recording for plasma/gelatin veronal buffer with fluorescent C3 and protein-liposomes with surface protein spacing = 25 nm

**Supplementary Movie 11.** Fluorescence nanoparticle tracking analysis recording for plasma/gelatin veronal buffer with fluorescent C3 and protein-liposomes with surface protein spacing = 29 nm

**Supplementary Movie 12.** Fluorescence nanoparticle tracking analysis recording for plasma/gelatin veronal buffer with fluorescent C3 and protein-liposomes with surface protein spacing = 36 nm

**Supplementary Movie 13.** Agent-based model of complement deposition on a sphere with surface protein spacing = 17 nm

**Supplementary Movie 14.** Agent-based model of complement deposition on a sphere with surface protein spacing = 24 nm

**Supplementary Movie 15.** Agent-based model of complement deposition on a sphere with surface protein spacing = 34 nm

### SUPPLEMENTARY SECTION C: Details of Computational Modeling

Through experimental observations, we established that the complement system displays a threshold-like activation dependent on a single geometric parameter – the average distance between potential complement (C3) attachment sites on a given material surface. We turn to computational modeling to understand how this surprising threshold behavior manifests in a complex network of complement proteins. Here, we employ multiple, distinct computational models, including systems of coupled ordinary differential equations (ODEs) and agent-based models (ABMs). We found all models exhibiting the same surprising trend: the threshold behavior results from a percolation-type transition and is universally captured irrespective of model details or parameter values. Further, we identified a small subnetwork within the large complement cascade that accounts for this behavior. These models predicted additional trends, which we then observed experimentally, such as “runaway” spreading C3b spreading across a single nanoparticle, which only occurs below the critical threshold, producing a small population of nanoparticles that C3b completely coats, and a large population with low C3b coating.

Our computational models focused on the complement system’s alternative pathway (**AP**) since the experimental data indicated that the observed threshold does not require the other branches of the complement cascade (the classical and lectin pathways). Additionally, we modeled the AP’s interaction with the particles for which we could best experimentally control and measure nano-scale properties: liposomes conjugated to a model protein (here IgG, which models antibody-targeted nanoparticles). By focusing on the AP and these experimentally facile nanoparticles (**NPs**), we established an iterative scheme between experiments and computation in order to reveal the key underlying dynamics.

In Section S1 with an ODE model of the entire AP system interacting with nanoparticles in the bulk phase, (meaning we do not model individual nanoparticles, but rather we track the dynamics averaged over a population of particles). We show that this ODE model, which utilizes equations and parameters from the literature, actually recapitulates the threshold-like behavior seen experimentally.

We critically analyzed the agreement between the model predictions and the experimental results by focusing on one of the most relevant outputs: the threshold behavior of the complement species [C3a] or [C3b] versus the distance between attachment sites. Simple exponential curves did not produce good fits suggesting that the recruitment was not due to independent binding events. We next analyzed the dependence through fit to the Hill function, which showed reasonable fits for Hill coefficients > 1, suggesting a possible cooperative recruitment mechanism. However, as multiple mechanisms can underlie cooperativity (e.g., simple nucleation and growth resulting from preferential stabilization of one state versus critical phenomena resulting from a fluctuations-driven transition), we needed to perform statistics on the fluctuations as well as a careful analysis of system-size-scaling of the fluctuations during the transition process.

In Section S2, we employ a spatially resolved lattice-based ABM of a single nanoparticle, which models the mean and the fluctuations about the mean and we found that our ABM also recapitulates the threshold-like behavior. Further, the ABM results showed that below the same threshold distance between potential C3 attachment points, two additional phenomena arise: i) a sharp rise in the probability of a “runaway event,” in which a single initial C3b attachment to the nanoparticle leads to a chain reaction spread of C3b attachments across the entire nanoparticle; ii) a sharp change in the time course of active C3b-surface adduct concentration. Both of these phenomena arise from a sharp change in the probability distribution of fluctuations within the system. Such large changes in the fluctuations are reminiscent of a critical phenomenon (or a phenomenon associated with collective dynamics in the proximity of a critical or bifurcation point) associated with certain phase transitions, as described in statistical mechanics and network science. Therefore, in Section S2, we continued by investigating how various fluctuation quantities (standard deviations) in system output change near the threshold, and found they increased dramatically around the threshold, and increased as the physical size of the nanoparticle increased. These changes are most consistent with a critical phenomenon, rather than competing models, like nucleation-and-growth. To further understand the critical phenomena and characterize the nature of the critical point, in Section S3 we identified a “reduced ODE model,” which contains only a small subset of reactions of the AP, but recapitulates the threshold behavior observed experimentally. We show that the reduced model can be mapped to the well-studied model, SI, already known to display critical phenomena in the percolation universality class. Finally, in Section S4, we describe the power-law behavior associated with the critical threshold and the critical exponent calculation of the threshold phenomenon, showing that it self-consistently fits all our computational and experimental data with the same exponents, and thus clearly establishing that the complement system displays a percolation-type critical threshold.

#### S1. ODE model

Zewde et al. [1] developed a quantitative model for the complement system’s alternative pathway. This model is defined by 107 ODEs, corresponding to various proteins, protein fragments, and protein complexes of the AP, and 74 kinetic parameters. The values of these kinetic parameters and molecular concentrations were based on numerous direct experimental measurements and estimation techniques based upon structurally/functionally homologous proteins, association/dissociation rate constants, and calculations based on diffusion rates in blood, and optimization procedures [1].

##### S1.1 ODE Model Description

In this section, we describe the application of Zewde’s model to our system of NPs. The supplementary material of [1] describes the amplification loop of the AP on cell surfaces; we have modified this part of the model to incorporate complement amplification on the NP surface (see Fig SC1). To tailor the ODE model to our system, we parametrize a key initial condition to the model, namely the concentration of potential C3b binding sites on NP surfaces. Notably, because each surface protein (IgG) on a NP has a smaller hydrodynamic radius than the C3bBb complex (which catalyzes cleavage of additional C3s), we assume that each surface protein (here, IgG) can be conjugated to only one C3b. Consequently, each surface protein acts as a single potential attachment site for a single C3b, forming a single C3b-nucleophile adduct on the NP surface. The initial value for the concentration of potential C3b binding sites on NP surfaces matches that used in our paper’s experiments. It is calculated in the following way:

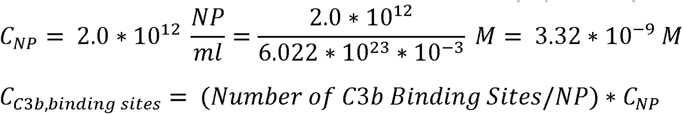

**Figure SC1:**
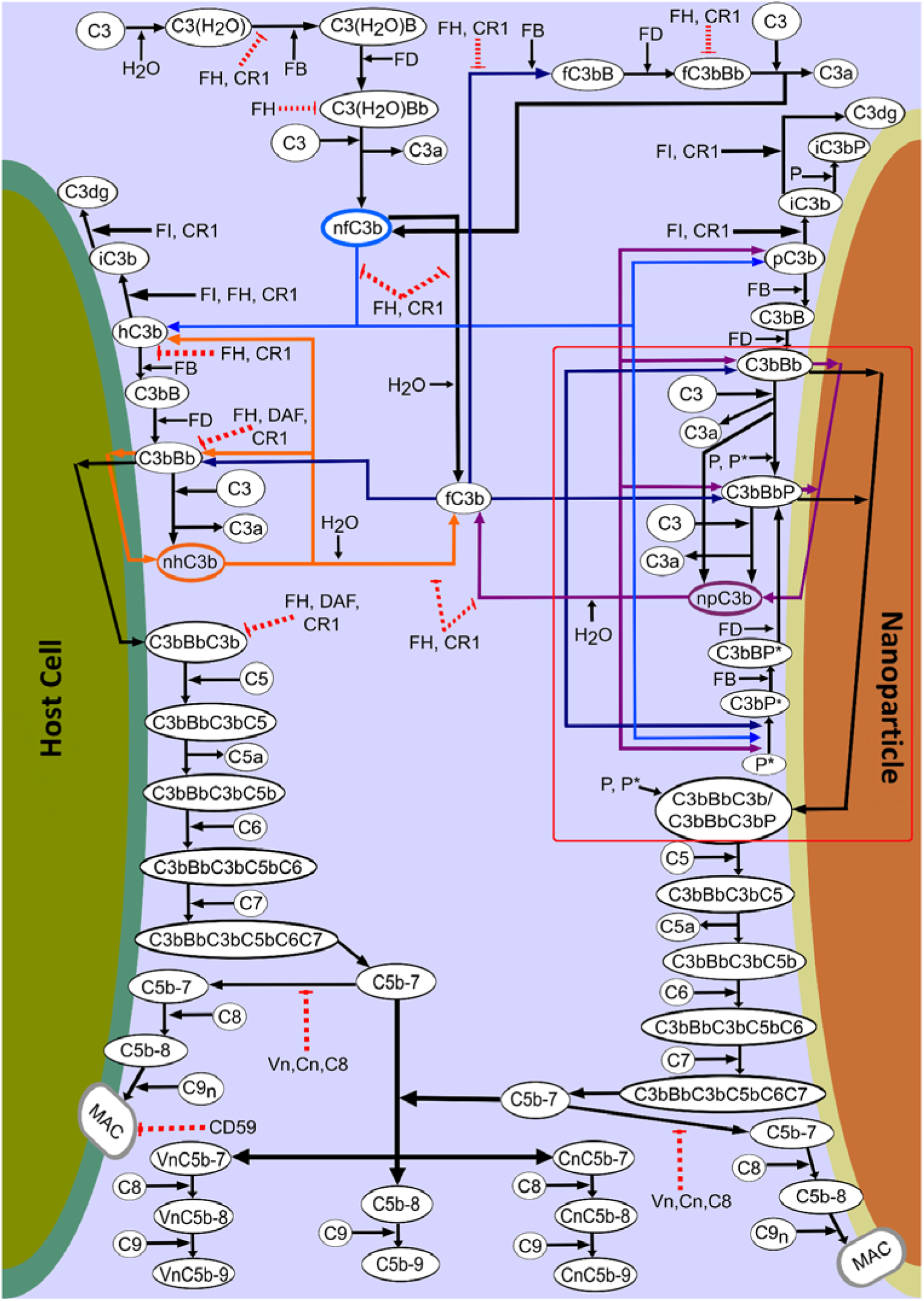
Complement proteins and protein complexes of alternative pathway (AP). Note that the amplification loop is boxed in red. The graphic was modified from [1], under Creative Commons License.

Here the “Number of C3b binding sites/NP” varies from 5 to 200 (equivalent to 5 - 200 IgGs we put on liposomes in experiments), and we obtain the steady-state value of various species by solving the system of ODEs.

##### S1.2 ODE Model Results

The 107 ordinary differential equations model was solved using Julia’s DifferentialEquations.jl [4] package. Supplementary material of [1] provides the initial concentrations of various complement proteins and the kinetic parameters. This section describes the calculation of the initial value of the concentration of C3b binding sites used to solve the initial value problem described by the system of ODEs.

To investigate what components of the complement AP drive the dynamics, we performed a global sensitivity analysis (**GSA**) [8], using the GlobalSensitivty.jl package in Julia. Specifically, our GSA examined the significant system parameters associated with the steady-state concentration of C3a. Figure SC2 shows the first and total order Sobol indices obtained from GSA. The GSA suggests that the kinetic parameters affecting the steady-state concentration of C3a are the attachment rate of C3b to various C3 convertases, which leads to the formation of C5 convertase and thus decay of catalytic activity to cleave other C3s, and the cleavage rate of C3 by C3 convertase. These two rates are the opposing forces to the amplification loop, the former inhibiting amplification and the latter supporting it.

Next we examined how the ODE model responds to changes in the distance between potential C3 attachment sites on a nanoparticle (e.g., modeling average distance between surface-conjugated IgGs on a liposome). Figure SC3 shows the steady-state concentration of C3a and the maximum value of surface-bound C3b as a function of the average distance between potential C3 attachment sites. This metric is equivalent to #binding sites/NP, and we calculate it by assuming that binding sites are distributed uniformly over the nanoparticle surface. As shown in Figure SC3, the ODE model displays a threshold phenomenon strikingly similar to that observed experimentally by changing the initial concentration of surface sites.

**Figure SC2:**
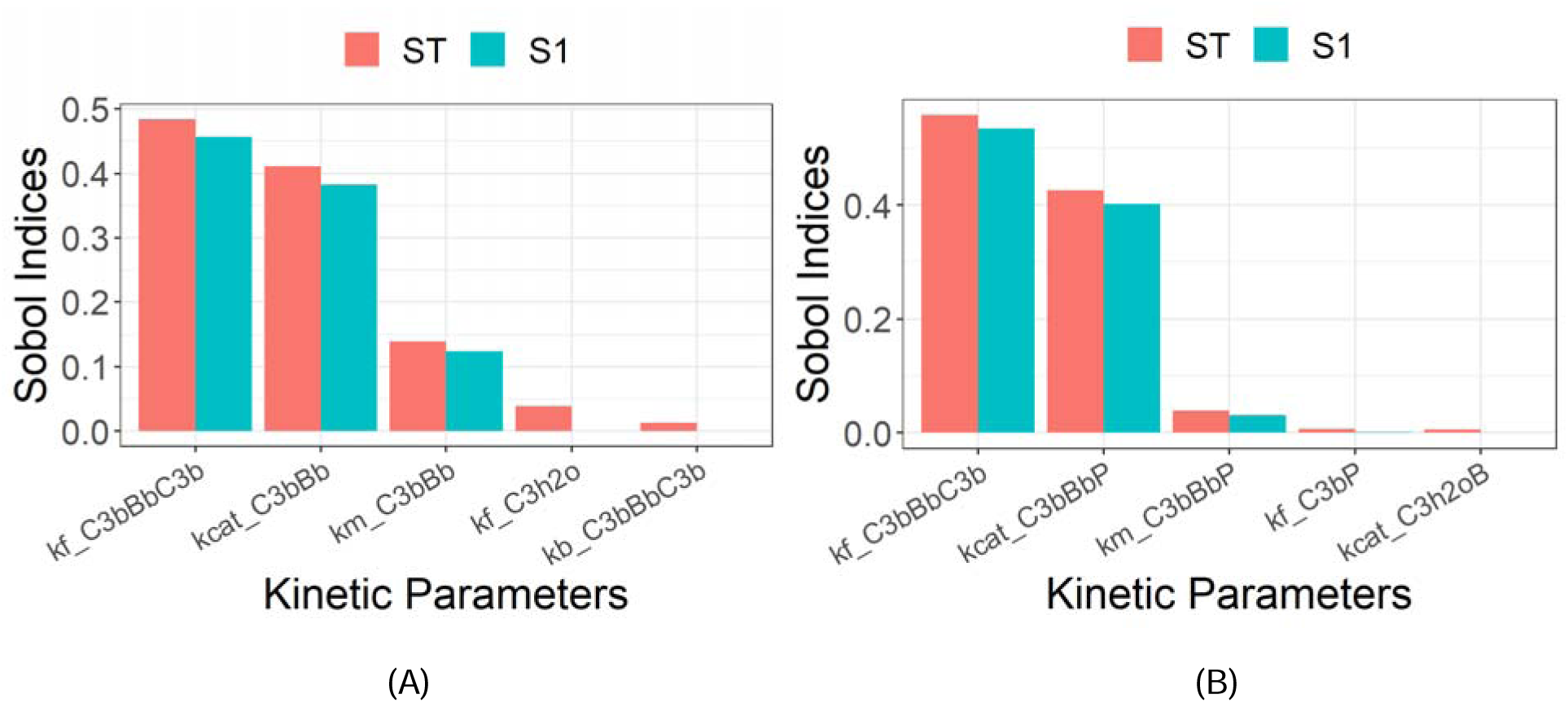
Global sensitivity analysis shows that amplification loop parameters have the greatest impact on complement activation. First order (S1) and Total Order (ST) Sobol Indices obtained from Global Sensitivity Analysis (GSA) of the ODE model. The GSA was performed in the absence (A) or presence (B) of properdin, since experimental data showed the very large impact of properdin on the threshold phenomenon. As shown above, the GSA suggests that parameters associated with the amplification loop strongly affect the steady-state value of C3a.

**Figure SC3:**
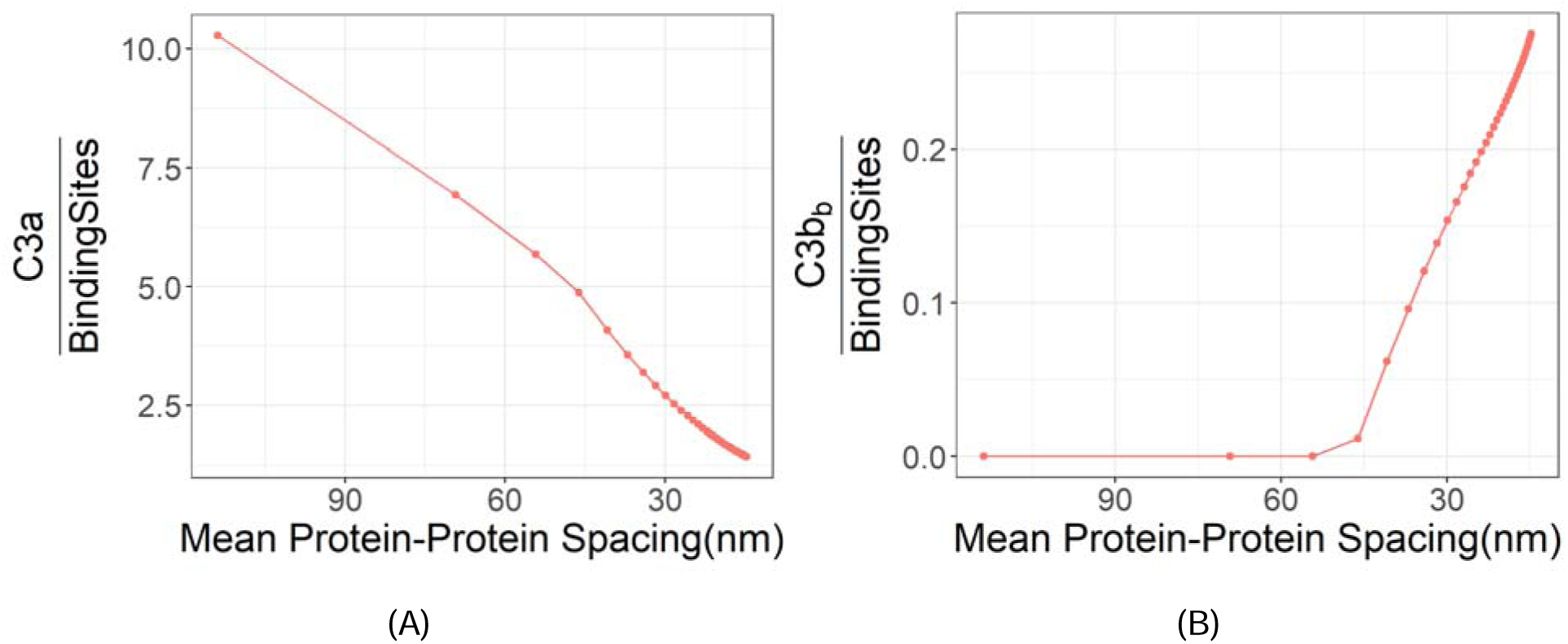
The C3b response per unit binding site displays threshold-like behavior. In the model, we varied the spacing between IgG molecules, modeling the assumption that each IgG can serve as the attachment point for a single C3b. The maximum value of surface-bound C3b (B) displays a threshold-like behavior.

#### S2 Agent-Based Model (ABM)

##### S2.1 ABM Description

We next wanted to approach computational modeling from a single nanoparticle point of view. We also wanted to explore complement dynamics to be spatially resolved on a discrete lattice, since lattices have been very fruitful for studying networks’ thresholds. Therefore, we constructed an agent-based model (**ABM**) to understand the spreading of C3b on the surface of NPs, allowing us to investigate whether an amplification loop will produce a sharp threshold in the relationship between complement activation and surface density of nucleophiles. The ABM is a quasi-2D simulation where the agents can diffuse over the 3D spherical surface (see Fig SC4). The agents refer to the immobile binding sites spread uniformly on a grid, and the C3b protein exists in two states, bound and unbound. Every agent has a set of properties, namely: id: a unique identifier; position: location of the agent; velocity: the speed at which the agent moves over the grid; type: agent type (binding site or C3b); engaged: an indicator for C3b-binding site engagement; state: dead/alive (meaning able to act or not in the relevant reactions; and age: lifespan of the agent, used to cause the decay. The simulation begins with randomly adding a C3b agent on a binding site. A bound C3b agent recruits more C3b agents to the surface, which move randomly over the grid and can either bind to an unoccupied site when in contact or decay because of aging. The recruitment process is also time-dependent, where the bound C3b agent eventually decays and can no longer recruit more C3bs to the surface. The bound C3b agent eventually decays and loses its catalytic activity. Fig SC4 shows how these rules work together.

**Figure SC4:**
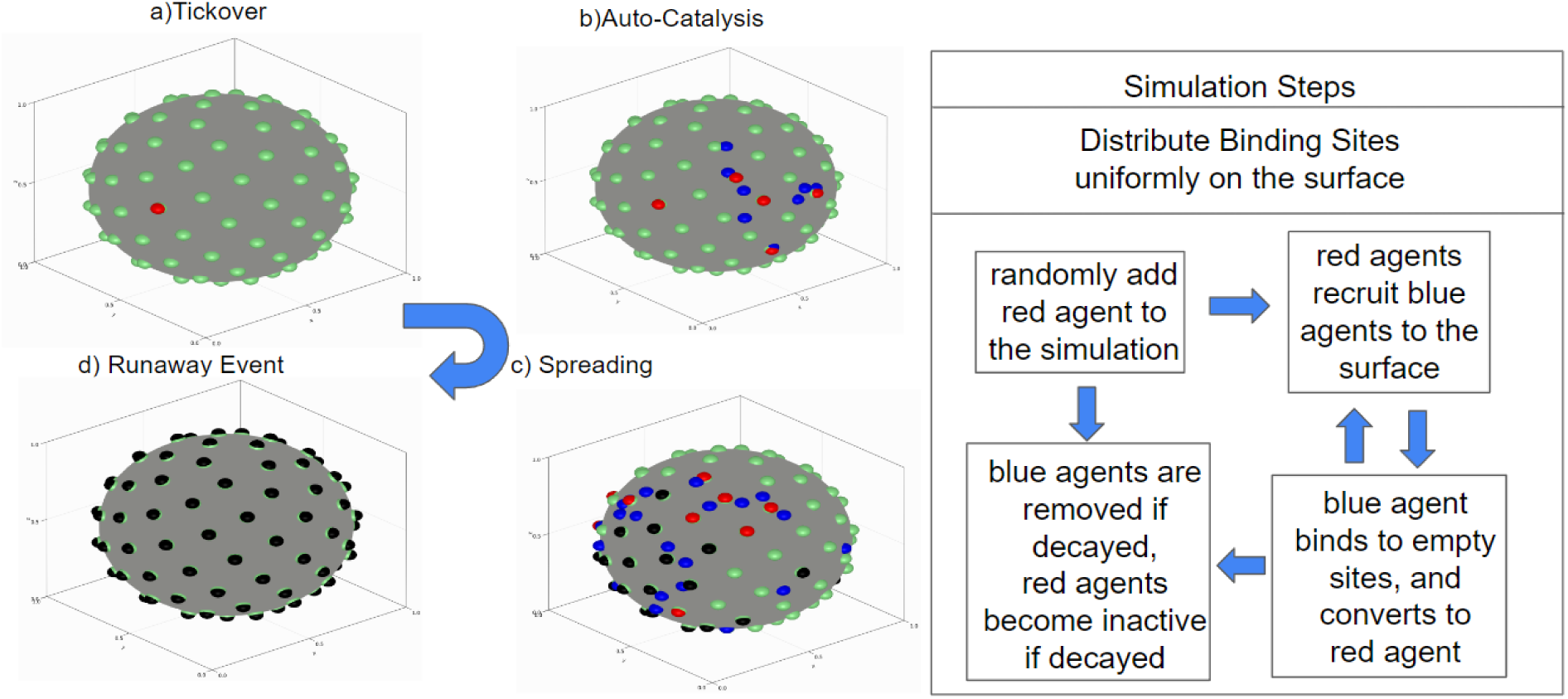
Agent-Based Model schema. We first define the agents: Green Agents: Binding Sites; Red Agents: Bound C3b / C3-Convertase; Blue Agents: Diffusing C3b; Black Agents: Decayed C3-Convertase. On the right above is a summary of the rules by which the agents interact with each other. On the left above is a set of 4 images from an ABM simulation run. In (c), we define “spreading” to mean spread of C3b-surface adducts across the nanoparticle’s surface, as a wave emanating from the initial C3b-surface adduct. Additionally, in (d), a “runaway event” refers to the situation where spreading continues such that all binding sites become occupied by C3b-surface adducts. Note that in (d) all agents have become black because the C3b-surface adducts eventually decay to a catalytically dead state (no longer able to catalyze cleavage of C3).

We tuned two parameters to map the ABM simulations to observed experimental time. First, we set the decay rate of blue agents (unbound C3b) equal to one-third of the red agents’ decay rate (bound C3b), while experimentally this ratio is much less than one-third [1]. Second, and more importantly, we set the blue agents’ recruitment rate to 15 times that of the tick-over rate, while experimentally this ratio is much larger [1]. This is justified for the following reasons. The process of autocatalysis (complement amplification) happens significantly faster (orders of magnitude) than tick-over (which is spontaneous activation of C3 to C3-H_2_O, which can attach to surface nucleophiles without first becoming C3b). We had to set a high tick-over rate in order to initialize the simulation. Similarly, in the ODE model, we also had to set the initial concentration of to a nonzero value. The smaller ratio between the decay rates and the larger tick-over rate can be justified because the ABM simulation aims to determine the critical spacing between binding sites above which runaway events can be observed. The tick-over does not play any role in this, as it is a random addition of a red agent. In other words, tick-over is simply an initiation event for the entire cascade, which requires tuning in simulations to make them proceed at a reasonable simulation time. Indeed, sensitivity analysis showed that the smaller ratio between the decay rates simply shifts the baseline of the fraction of occupied sites (around ∼20%; see Fig SC5).

##### S2.2 ABM Results

We simulated the agent-based model described above using Julia’s Agents.jl [5] package. Figure SC5 shows the fraction of sites occupied as a function of the spacing between binding sites and the histogram for a different number of binding sites per NP. Note that the fraction of occupied sites is a proxy for bound C3b concentration and is therefore compared directly with ODE models and experimental data. We can see a threshold around the 30-20 nm region, similar to experimental results.

###### S2.2.1 Runaway events and rapid transients in the ABM suggest that system fluctuations become larger near the threshold

To understand how system fluctuations impact the system behavior, we next examined the histograms (Fig SC5) of the number of binding sites occupied at the end of each of a series of 400 simulations. As we increase the number of binding sites per NP (i.e., decrease the average distance between binding sites), a clear transition occurs, from a right-tailed distribution to a left-tailed distribution. The shift in distributions indicates a sharp change in system behavior as the spacing between binding sites decreases: below the 30-20 nm region threshold, there suddenly emerges a nonzero probability of “runaway events,” in which a single C3b binding to the nanoparticle can catalytically spread a wave of C3bs across the entire nanoparticle, covering nearly every potential C3 attachment site.

Next we examined the time course of ABM simulations (Fig C6), using the same simulations as in Figure SC5 B-E. In Figure SC6, the y-axis represents the number of *active* agents (red and blue, representing catalytically active C3b-surface adducts and C3b in solution) over the spherical surface. In Figure SC6, we can see a sharp peak and rapid decay from the maxima in the plots in which the distance between C3 attachment sites is below our typical threshold (6C & D). On the contrary, there are no sharp peaks when the nanoparticle has a distance between attachment sites above threshold (Figure S6A, B).

Notably, these runaway events and a transient spike in active agents are reminiscent of behavior seen in the critical phenomena associated with phase transitions, as described in statistical mechanics and network science. In particular, both of these properties are seen in the networks described by percolation theory. For example, the transient rise in panels S6C & S6D can be understood with an analogy to forest fires, one of the commonly studied *in silico* models in percolation theory. The shorter the distance between the two nearest trees, the more effective the spread of forest fire—similarly, the number of our model’s active agents (like fires) multiplies when the binding sites are close (similar to nearby trees), producing a sharp rise to peak. In a similar analogy to forest fire, the number of active agents (like fires) decays as blue and red ones die out. This decay in our ABM is more rapid when there are more binding sites, as the entire nanoparticle gets covered rapidly and generates many blue agents. For comparison, decay rates for the four cases shown in Figure SC6 are in increasing order of A<B<C<D. These decay rates are inversely proportional to the number of available binding sites post-peak.

**Figure SC5:**
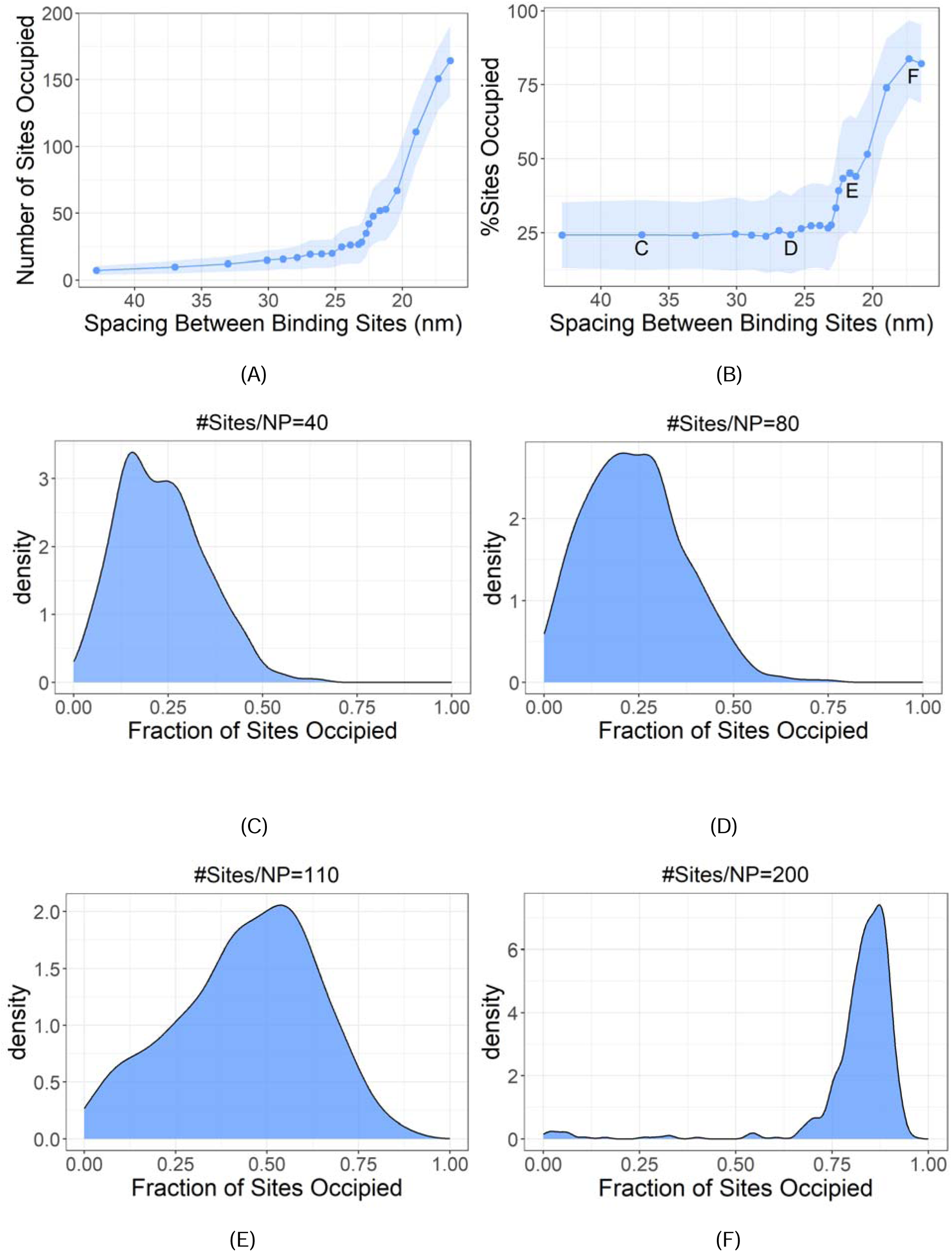
ABM simulations display threshold-like behavior, with runaway events rarely occurring if spacing between potential C3b attachment sites is below threshold. We used the ABM to simulate individual nanoparticles with different distances between their potential C3b attachment sites (modeling the average distance between, for example, surface-conjugated IgGs on nanoparticles).A) A plot of Number of sites occupied by C3b agents as a function of spacing between binding sites, B) A plot of the fraction of sites occupied by C3b agents as a function of spacing between binding sites, showing a threshold in the range of 20 - 30 nm, the error bar represents mean ± standard deviation of sites occupied at the end of 500 simulations, this metric isolates the effect of site spacing from the effect of just adding more sites to each particle. C - F) After 500 ABM simulations for a given number of potential surface attachment sites (e.g., IgG) per molecule, we plot the histograms of the number of occupied sites on a nanoparticle (e.g., # of C3-IgG surface adducts). The number of potential surface attachment sites (“Sites/NP) were for each panel: C = 40, D = 80, E = 110, F = 200. Note that “runaway events,” where nearly all sites on a nanoparticle become occupied by C3b, were not observed below the threshold from panel A (corresponding to B & C), but common well above threshold (E).

The peak tends to a plateau for case A, as the blue agents cannot spread effectively to occupy the binding sites. In contrast, the number of active agents in D decreases rapidly post-peak because most sites are already covered. Thus, at this point, we can say that by a heuristic analogy to the forest fire model of percolation theory, that the transition between the two behaviors of the complement AP system (comparing Figures SC6 A&B vs SC&D) appears to obey percolation-type critical threshold behavior. However, to more definitively determine if critical phenomena are occurring, we need to perform a system size scaling analysis, as we do in the next section.

**Figure SC6:**
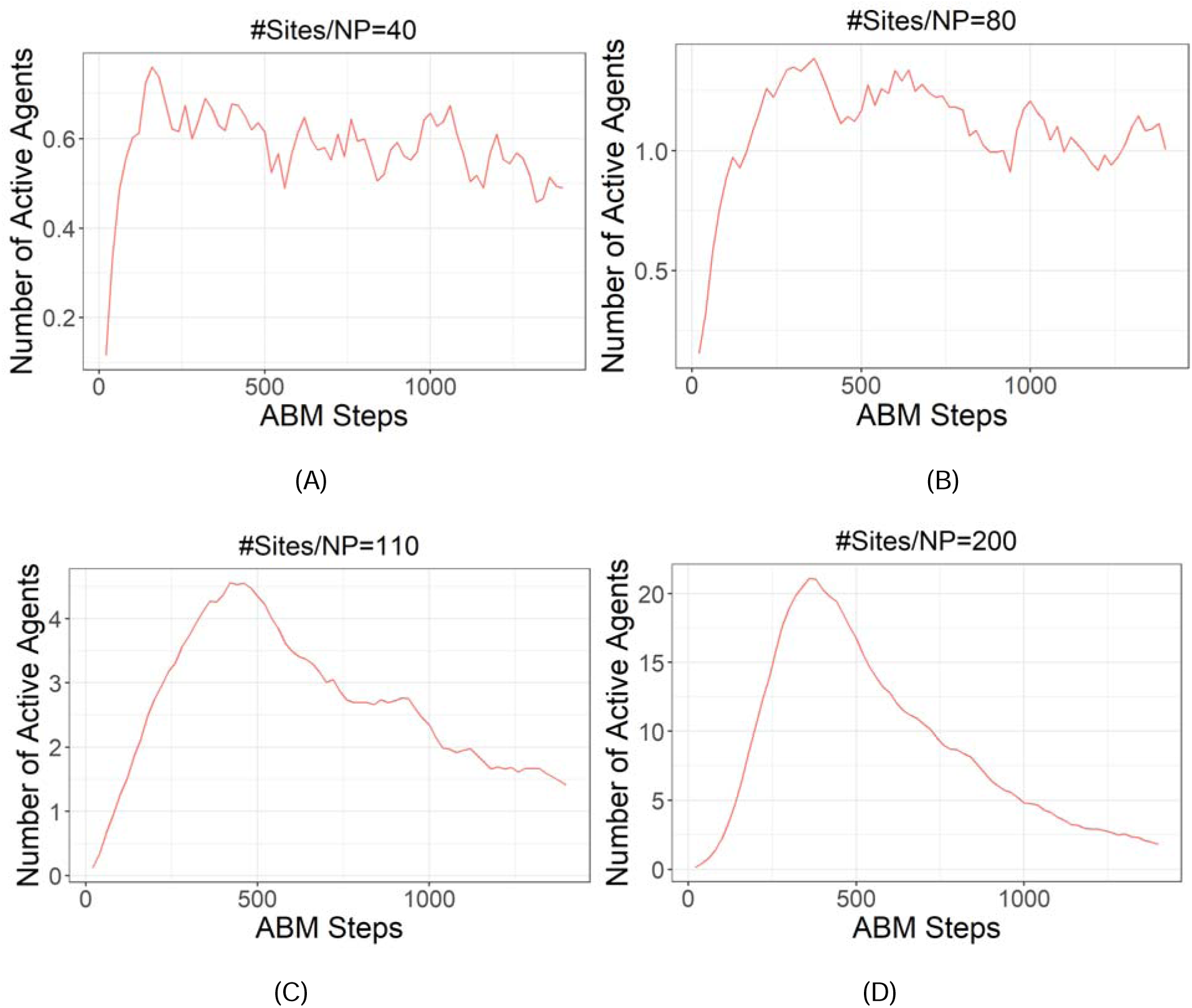
The ABM displays a sharp change in temporal dynamics when the spacing between potential C3b attachment sites dips below a threshold: Here we employed the same simulations as in Figure C5, but here we plot the average of the number of sites over time (here measured in steps in the simulation). A to D corresponds to decreasing spacing between binding sites on a NP. (A) and (B) show a sharp rise in the number of active agents followed by slow decay, while C) and D) show a sharp increase in the number of active agents followed by a quick decay, indicating runaway events have occured.

###### S2.2.2 System-Size Scaling

To establish that the observed threshold behavior is associated with a critical phenomenon, we next analyzed system behavior as we scaled the system size. As our base case, we used d=1 unit as the baseline diameter of the NP surface. This baseline diameter was mapped to the diameter of NPs used in experiments (150 nm). We explore the trend behavior for systems with 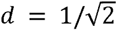 (half the area) and 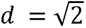 (double the area) in Fig SC7.A. In Figure SC7.A (left column), we can see that all three systems follow a similar trend of threshold-like behavior. Fig SC7.B (right column) highlights that the standard deviation of the fraction of sites occupied *peaks* in the region of criticality. Such a peak in standard deviation is characteristic of a critical system.

**Figure SC7:**
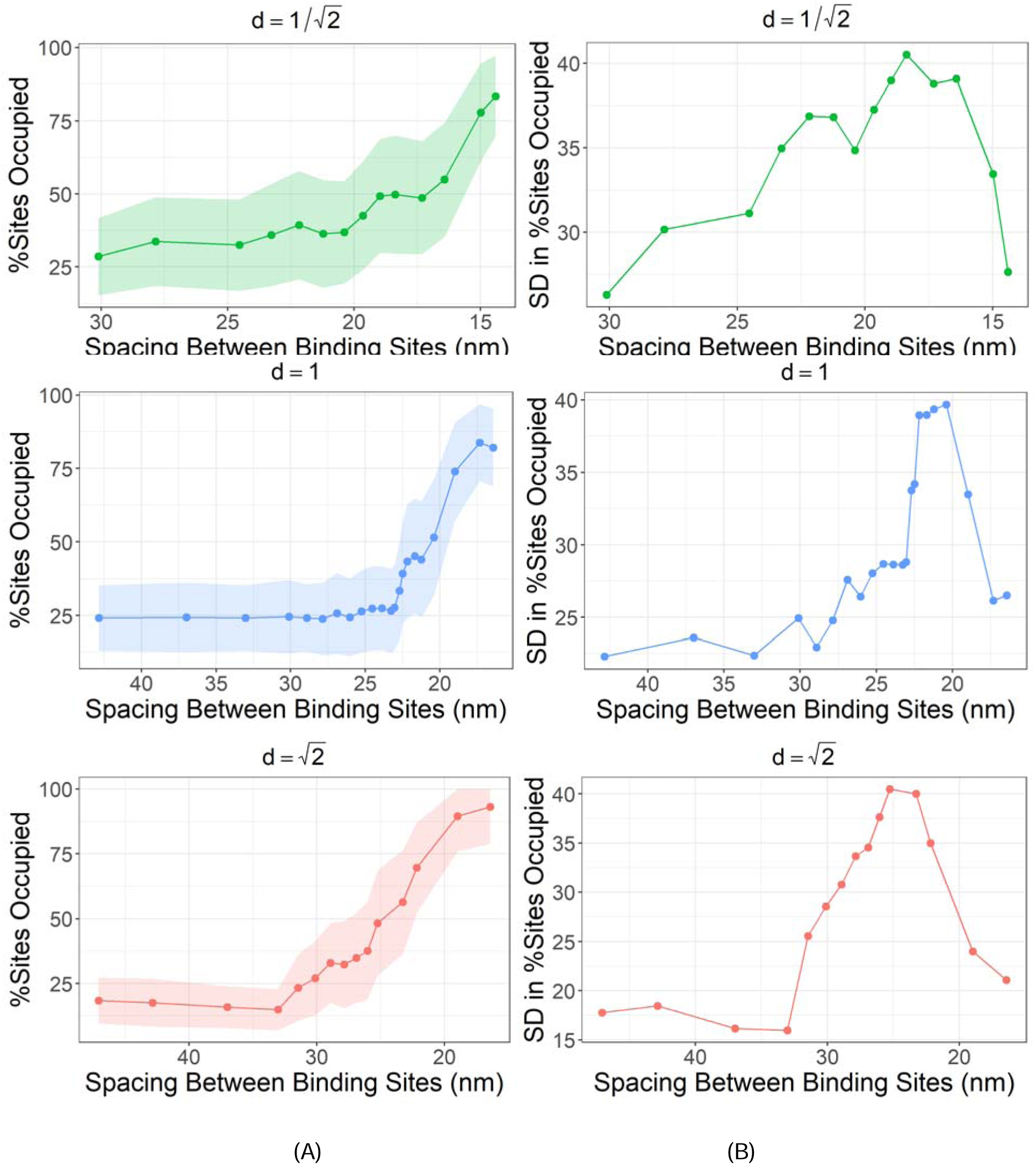
System size scaling demonstrates that the ABM of the complement AP displays critical phenomena: We conducted ABM simulations for systems with different NP diameters to compare the trend and standard deviation in the fraction of sites occupied. A) Fraction of sites occupied at the end of simulation as a function of the spacing between binding sites, B) Standard deviation (SD) in the fraction of sites occupied as a function of the spacing between binding sites, for three cases 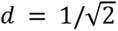, *d*= 1, 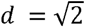 (top-bottom). Note the peak in SD at the threshold, which is characteristic of critical phenomena at their region of phase transition.

Keeping the #sites/NP/Area constant, we further explored the maximum number of active agents and standard deviation (of the maximum number of active agents) as a function of the system’s physical size for a particular value of spacing between binding sites. This corresponds to 200 sites/NP for d=1. Fig SC8A shows that, as the system increases in physical size, the maximum number of active agents grows, and the maximum number of agents refers to the maxima of Fig SC6D, for different system sizes. Fig SC8B shows that, as the system increases in physical size, the fluctuations in the maximum number of active agents also grow. The error bars in the maximum number of active agents and standard deviation in the maximum number of active agents are computed by binning the 500 simulations in 5 sets of 100 simulations, each set gives a value of mean and standard deviation in the maximum number of active agents for the 100 simulations in that set.

Collectively, the system size scaling results strongly indicate that ABM is displaying critical phenomena. If the system was instead governed by simple nucleation-and-growth, the fluctuations in the system (e.g., fluctuations in the number of active agents) would not increase as the system size increases, but would instead decrease (a simple result of the Central Limit Theorem). By contrast, near the critical point of phase transitions, fluctuations within a single system (Fig SC8.B) and variance between multiple instances of a system (Fig SC8.A) increase as the system size increases.

**Figure SC8:**
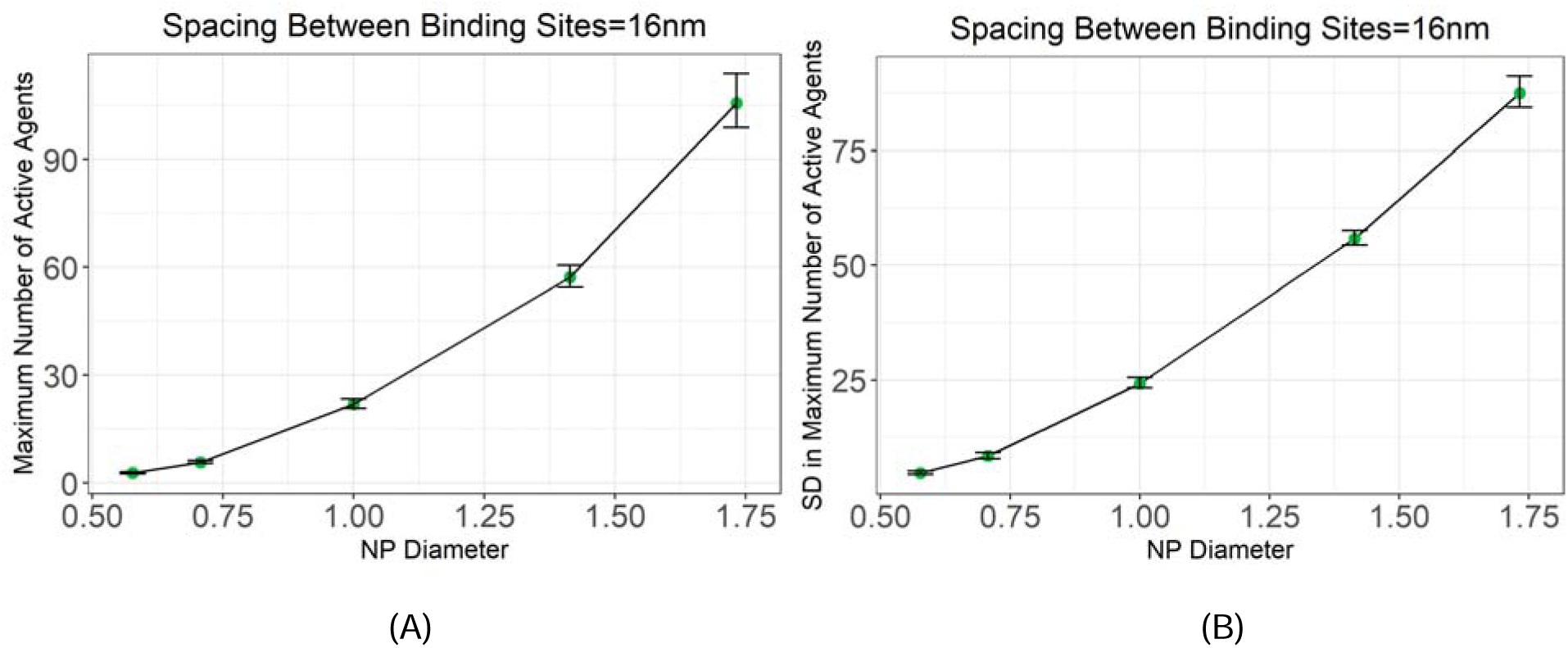
Fluctuations grow with system size, a property of criticality: We start with nanoparticles whose spacing between attachment sites was below the threshold observed previously. We fixed the spacing between the binding sites, but increased the physical size of the particles. We then ran 400 simulations of nanoparticles at each size. In (A) we plot the mean (and standard deviation around the mean) of the maximum number of active agents that occur during the course of each simulation run. Note the increase in the standard deviation with system size. For B, in each simulation run, we measured the standard deviation of the number of active agents (which fluctuates over time), and this measure shows fluctuations increase with system size.

#### S3.1 Reduced ODE model

Given that the ABM indicated that the complement cascade’s observed thresholds are a type of critical phenomena, we next wanted to characterize scaling behavior close to the critical point. Of course, the full ODE model is excessively complicated (107 ODEs, 70 parameters), due to the web of interacting proteins in the complement system. However, due to the universal nature of critical phenomena, we hypothesized that a reduced subset of this network could explain the threshold phenomena and the critical behavior.

Therefore, we compared the AP’s behavior to various networks displaying critical thresholds, such as the Ising model, percolation, and more. We noted that the ABM (Section S2) movies pictorially resembled the spread of infectious diseases within a population. Since the 1920s, spread of infectious agents has been commonly modeled by the **SI** (susceptible-infected) model of 2 coupled ODEs, which displays critical threshold behavior when projected onto a discrete lattice. We therefore attempted to model the AP as if it were an infectious disease spreading, using the terminology of the SI model. Here, potential C3b binding sites represent the hosts and C3b the infectious organism. We tested several variations of the SI model, such as the SEIR and **TIV** models, and found that the TIV models’ 3 coupled ODEs fit precisely with the dynamics of the AP (see equations below for mapping). This allowed mapping the full ODE model to the TIV-like model [2] to explain the threshold phenomena. We further show that this TIV-like model collapses into a SI-like model [3] under precisely stated simplifying assumptions. Given that the SI model is known to be in the percolation universality class, pur mapping provides further evidence that the complement AP’s threshold-like behavior is due to percolation-type criticality, which we explicitly demonstrate below.

#### S3.2 TIV model and comparison with our reduced model

***TIV model:*** The TIV variant of the SI model has been used to study spread of viruses (V) among infected cells.

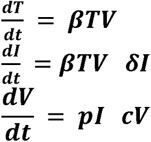

**Reduced ODE model of the complement AP:**

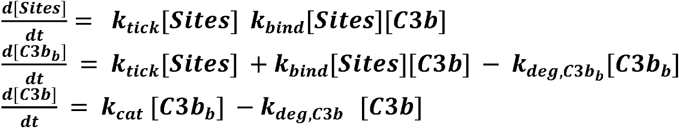

In the reduced ODE model of the complement AP, the variables and parameters are:

[***Sites***]: Concentration of surface binding sites
[***C3b_b_***]: Concentration of bound C3b/C3b protein complex
[***C3b***] : Concentration of Free/unbound C3b

***k_tick_***: Slow tick-over/binding rate of C3b with surface sites independent of amplification loop
***k_bind_***: Binding rate of C3b with surface sites due to amplification loop
***k_deg_***: Degradation/Decay rate of C3b or Surface bound C3b
***k_cat_*** : Autocatalysis rate of cleavage of C3.

Importantly, if we assume ***k_tick_***= 0, then the two models (TIV vs reduced ODE model of the AP) are equivalent in the following sense: they have identical form of their ODEs, only changing the names of the variables and parameters. The assumption of ***k_tick_***= 0 is valid because the rate of C3 cleavage by tickover is orders of magnitude lower than C3 cleavage produced by the amplification loop. To put this assumption into practice, in the reduced model, we set ***k_tick_***= 0 and set the initial value of surface-bound C3b to a very small non-zero value to start the amplification loop. Figure C9 shows the surface-bound C3b using the TIV representation. As with our Full ODE model of the complement AP, this TIV-like, “reduced” ODE representation of the complement AP also shows threshold-like behavior, with a similar numerical value for the threshold.

##### S3.2.1 Relation between TIV-like model parameters and Full model parameters

The TIV model parameters are defined in the following.

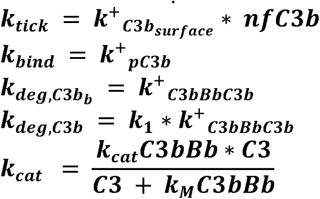

***nfC3b*** : Nascent fluid phase C3b
***C3*** : Concentration of protein C3
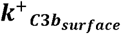 : Attachment of nfC3b to NP
***k^+^_pC3b_*** : Attachment of C3b to NP
***k^+^_C3bBbC3b_*** : Attachment of C3b to C3bBb
***k_cat_C3bBb, k_M_C3bBb*** : Cleavage of C3 by C3 convertase, C3bBb
***k*_1_ :** Proportionality constant

The values of above-defined parameters are taken from the supplementary material of [1]. The values of ***nfC3b*** is set to be constant ∼1e-6, which acts as baseline complement activity in the blood. The protein concentration C3 is set to be constant and equal to the experimentally measured value of 6 *µM.* The degradation rate constants have been set proportional to ***k^+^_C3bBbC3b_*** based on the global sensitivity analysis. The surface-bound convertase responsible for the cleavage of C3 converts into a C5 convertase, which slows down the amplification loop. This further cascades down to the production of membrane attack complex (MAC) [1]. However, the reduced model does not include the AP’s production of the MAC, as the MAC does not intercalate into nanoparticles but rather into larger cells, which are not modeled experimentally. Thus, the TIV-like reduced ODE model of the complement AP directly maps onto the Full ODE model, with realistic approximations and assumptions.

##### S3.2.2 Connection between TIV-like model and ABM

The ABM described above closely resembles the TIV representation in the following way:

Green Agents > [***Sites***]
Red Agents > [***C3b_b_***]
Blue Agents > [***C3b***]
Black Agents > Decayed [***C3b_b_***]

Here, we report the ABM statistics regarding the number of red, blue and black agents. The maximum value of [***C3b_b_***] in the TIV-like model corresponds to the number of black agents on the surface at the end of the simulation. Thus, for the complement AP, a TIV-like model not only has direct mappings to our Full ODE model, but also to our ABM.

#### S3.3 SI model and comparison with our reduced model

As the TIV model is much less studied than its close relative SI, we wanted to understand if there was a further mapping from not just our Full Model to the TIV-like model but from the TIV-like model to SI. TIV and SI have previously been linked via a quasi-steady-state assumption (QSSA) [6]. Therefore, we sought to determine if a similar QSSA could map our TIV-like model to the SI-like model and understand the physical meaning of the QSSA in the complement AP system. To do that, we first examined the time course of unbound C3b in the TIV-like model. We use a quasi-steady-state approximation for C3b, noting that the value of C3b is close to zero (due to its short half-life), with a sharp transient peak at the beginning of the time series (see Fig SC9. B). We can therefore set the derivative of C3b to zero. Further, if we assume 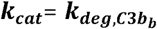, we essentially have a SI-like model.

Using the approximations as mentioned above, we set,

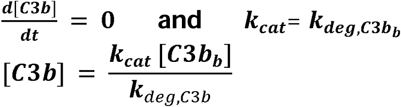

therefore eliminating [*C3b*] from the first two differential equations and setting the tickover rate to zero (***k_tick_*= 0**) we get,

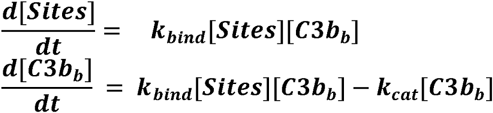

Where

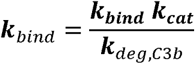

This is equivalent (simply by changing variable names) to the classic SI model of:

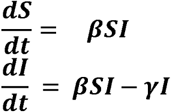

Figure SC10 shows the maximum value of bound C3b during the time-course and using the SI representation. Once again, we observe a threshold.

##### S3.4 Reduced Model Results

The reduced model defined in the subsections above consists of four ODEs in the TIV-like representation and three ODEs in the SI-like representation. These differential equations are solved in Julia and show a similar threshold in the relationship between complement activity and surface site density. Fig SC9 and SC10 show the steady-state concentration of surface-bound C3b as a function of average spacing between potential C3b attachment points obtained from the TIV-like and SI-like models, respectively.

There are two key takeaways from these reduced models. First, even though the complement AP requires 107 ODEs and 74 kinetic parameters to model every reaction, the surprising threshold-like behavior observed experimentally is nearly entirely explained by a very reduced set of reactions, modeled by just 3 chemical species and 2 reaction rates. Thus, consistent with the nature of critical phenomena being universal (i.e., not dependent on the detailed nature of the interactions, or specific parameter values, but rather applicable more broadly and across a wide range of systems), a seemingly enigmatic emergent phenomenon (thresholding) emerges not from the complicated complement AP, but from a tiny subset of the reactions. Second, the complement AP’s reduced ODE models map directly to the SI / TIV ODE models, simply changing variable names. Further, these reduced models map directly to the ABM. Thus, we can understand the complement AP in terms of the universality class into which SI fits, namely percolation models. This result fits with those of Section 2, in which we showed a phenomenon found in other percolation-class models: runaway events and increase in fluctuation size as the system size scales up.

**Figure SC9:**
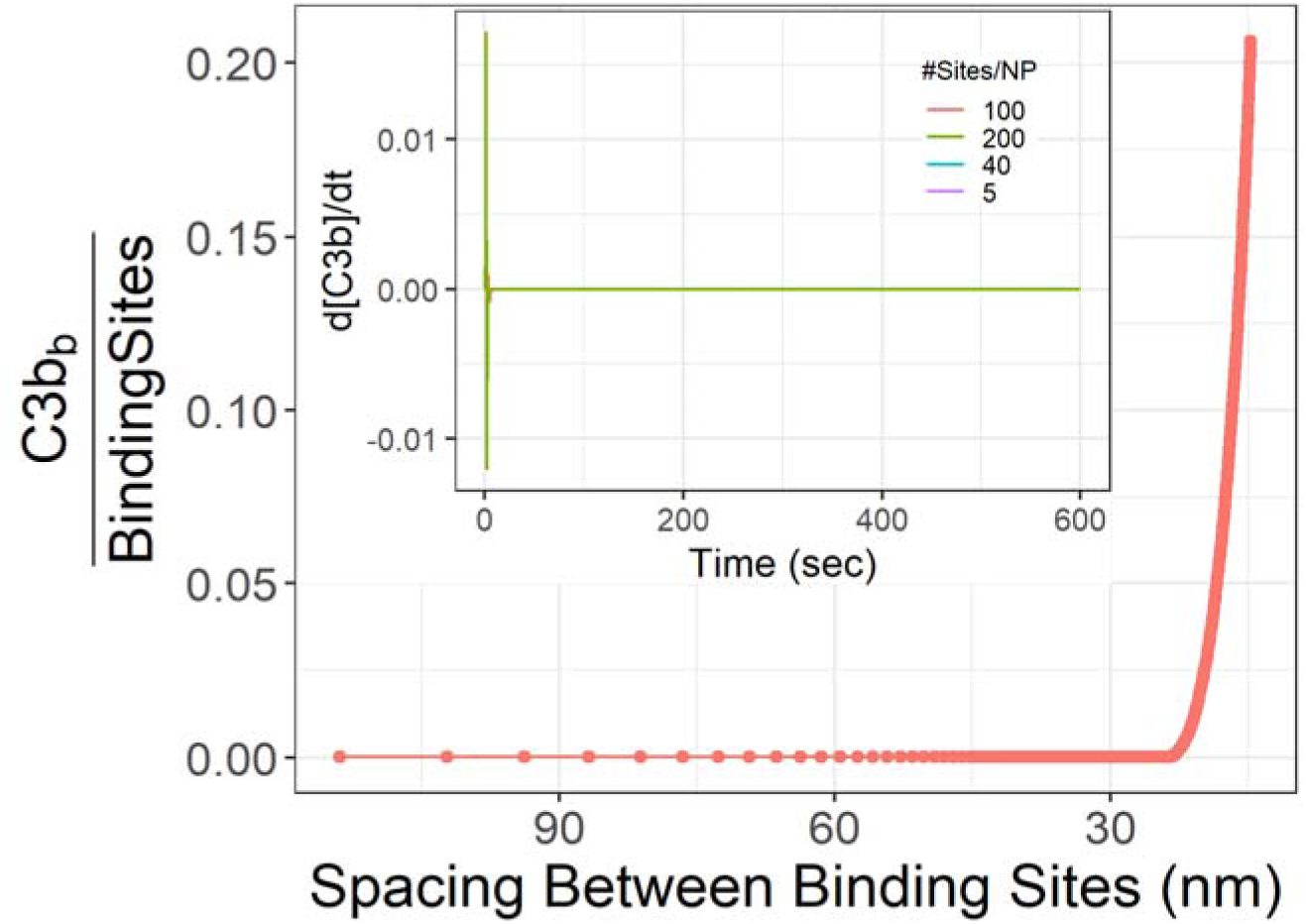
The TIV-like model of the complement AP displays threshold-like behavior. As in our prior models, we varied the spacing between potential C3b-attachment points on a particle (listed here as Average IgG-IgG Spacing, as one IgG may attach to one C3b). The maximum value of surface-bound C3b displays a threshold-like behavior in the region of 40-20 nm between IgGs, similar to the full ODE model. The subplot indicates that the concentration of unbound C3b does not vary with time.

**Figure SC10:**
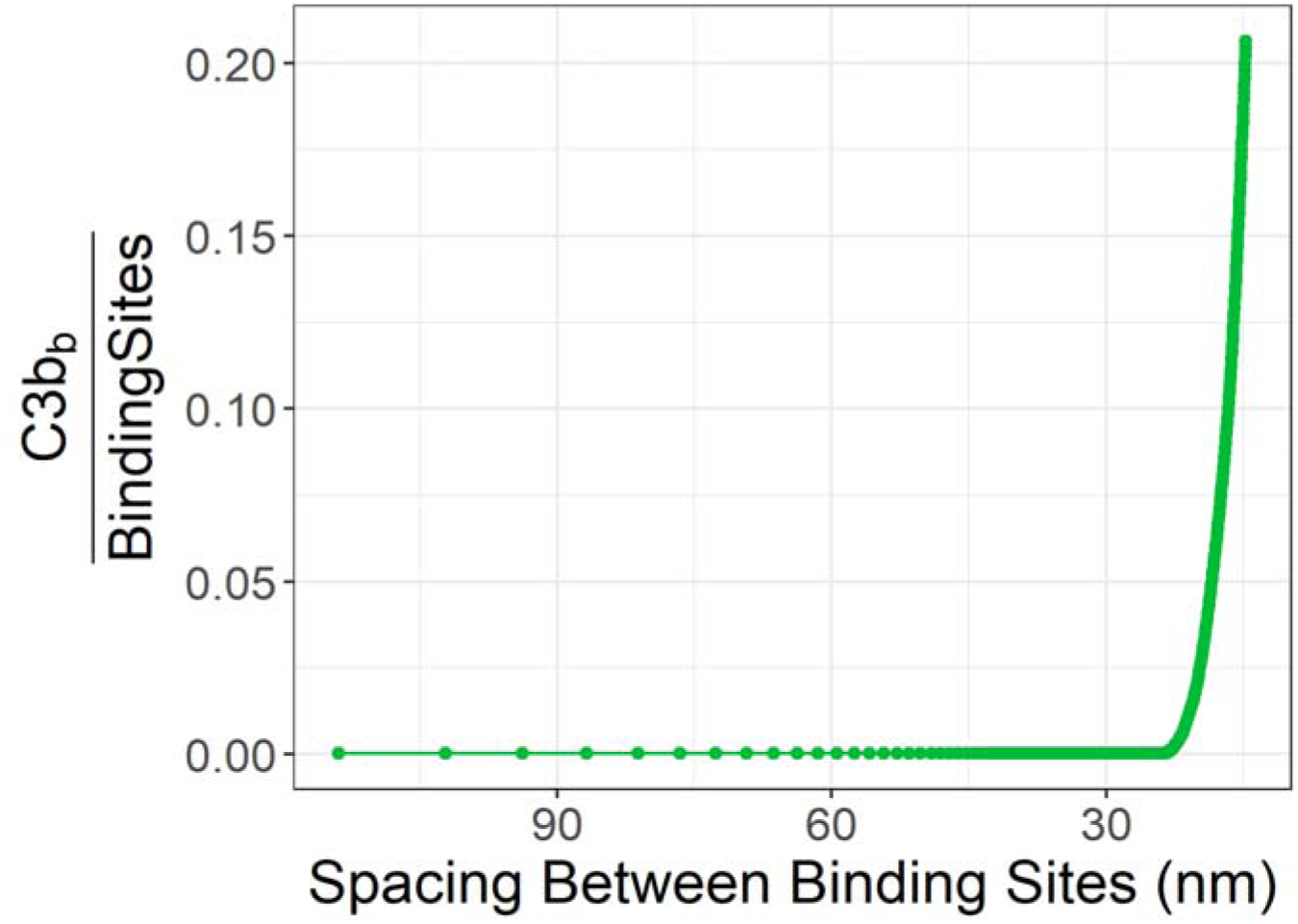
The SI-like model of the complement AP displays threshold-like behavior. The maximum value of surface-bound C3b displays a threshold-like behavior in the region of 40-20 nm between IgGs, similar to the full ODE model and TIV-like model.

#### S4.1 Critical Exponent Calculation To Explain Threshold Phenomenon

Sections S2 & S3 strongly indicated that the complement AP displays criticality and fits into the percolation universality class of networks. To finalize this classification, measuring the key descriptive parameters of a critical phenomenon, namely the critical exponents, is necessary. Therefore, we fit a critical point exponent model to the ODE simulation data. We begin by fitting critical exponents to the TIV-like and SI-like models. Each such fitting gives a locus of possible values of the critical exponent, with the locus not being a single point due to a free parameter in each model. As explained in [7], the model equation in the region near the critical point is:

[*C3b*] = *A*|*d_r_*|*^λ^ exp* (*Bd_r_*)

Where

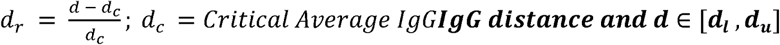

For this model we perform nonlinear least squares regression in the domain [***d_l_, d_u_***] and corresponding normalized [*C3b*] values to obtain the value of *λ, d_c_*, the leading coefficient A, and B in the exponential term. Although the critical exponent model is applicable in the 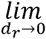, we apply the model in the vicinity of *d_c_*, such that |*d_r_*| < 1, the lower bound ***d_l_*** and the upper bound ***d_u_*** is manually chosen to satisfy this condition.

The critical exponent is given by [7]:

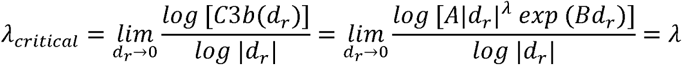

Figures SC11 and SC12 demonstrate the application of the critical exponent model to the maximum concentration of bound C3b during the time course for TIV-like and SI-like models, respectively. For these curves, we estimated the curve fit only using simulated data in the x-axis subregion of [35 to 25 nm] distance between potential C3b attachment sites. However, the estimated values of the critical exponents might depend on the size of the subregion of the x-axis; e.g., if we expand to fit the curve to simulated data over a range of 40 to 15 nm. Therefore, we next sought to estimate the locus of critical exponent estimates that fit the simulation data when we varied two elements: 1) the domain of spacing between attachment points that are included in each curve fitting (e.g., do we fit data over the range of 35 to 25 nm or 40 to 15 nm); and 2) varying the SI and TIV model parameter (***k_bind_***, see section S3.2 and S3.3). The resulting locus plots of the critical exponent (including the error bars) and critical distance for TIV-like and SI-like models are presented in Figures SC13 and SC14, respectively. The error bars in these plots represent the range of fitted values (of critical exponent or critical distance) obtained when we vary the domain of spacing (***d*** ∊ [***d_l_, d_u_***]) between potential C3b attachment sites, at a fixed value of the SI or TIV model parameters. The error bars show that the critical exponent is sensitive to the domain of spacing used in the curve fitting, while the critical distance is relatively unaffected by the spacing range. Having obtained a locus of estimated critical exponents (and critical distances) from each of the TIV- and SI-like models, we next wanted to know if our experimental and ABM data produce estimated critical exponents that fall into this locus. Therefore, we fit a critical exponent model to experimental data for [C3b], and separately fit the model to the ABM data, measuring the fraction of occupied sites at the end of the simulation, as shown in Figure SC15. The resulting estimated critical exponents and critical distances for both datasets fall within the critical exponent locus obtained from TIV-like and SI-like models. Therefore, we conclude that SI and TIV-like models effectively explain the threshold phenomenon.

This analysis unequivocally supports our conclusion that the complement AP’s threshold phenomena is explained by a small subnetwork of reactions, and that subnetwork (and the whole network) maps directly to the universality class that includes SI and percolation. Thus, the initially enigmatic emergent behavior of the very complicated network of complement proteins can be directly explained by the elegant phenomena associated with criticality.

**Figure SC11:**
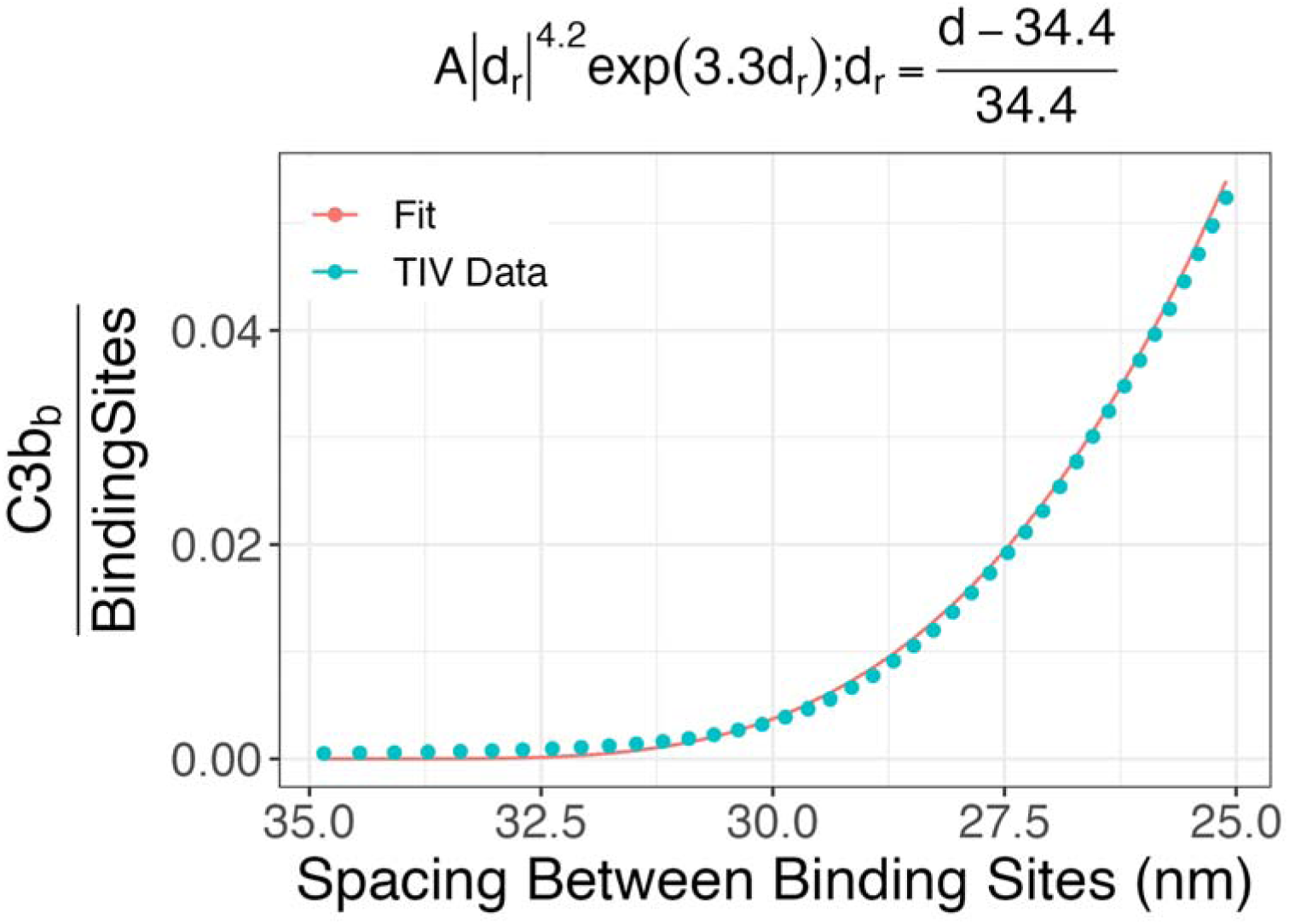
Critical Exponents explain threshold-like phenomena in TIV-like model: Here we plot the TIV-like model’s output (Surface Bound C3b) in the small physical distance around the theshold (35-25 nm between potential C3b attachment sites). We also plot a curve fit to that data, with the curve of the form listed above.

**Figure SC12:**
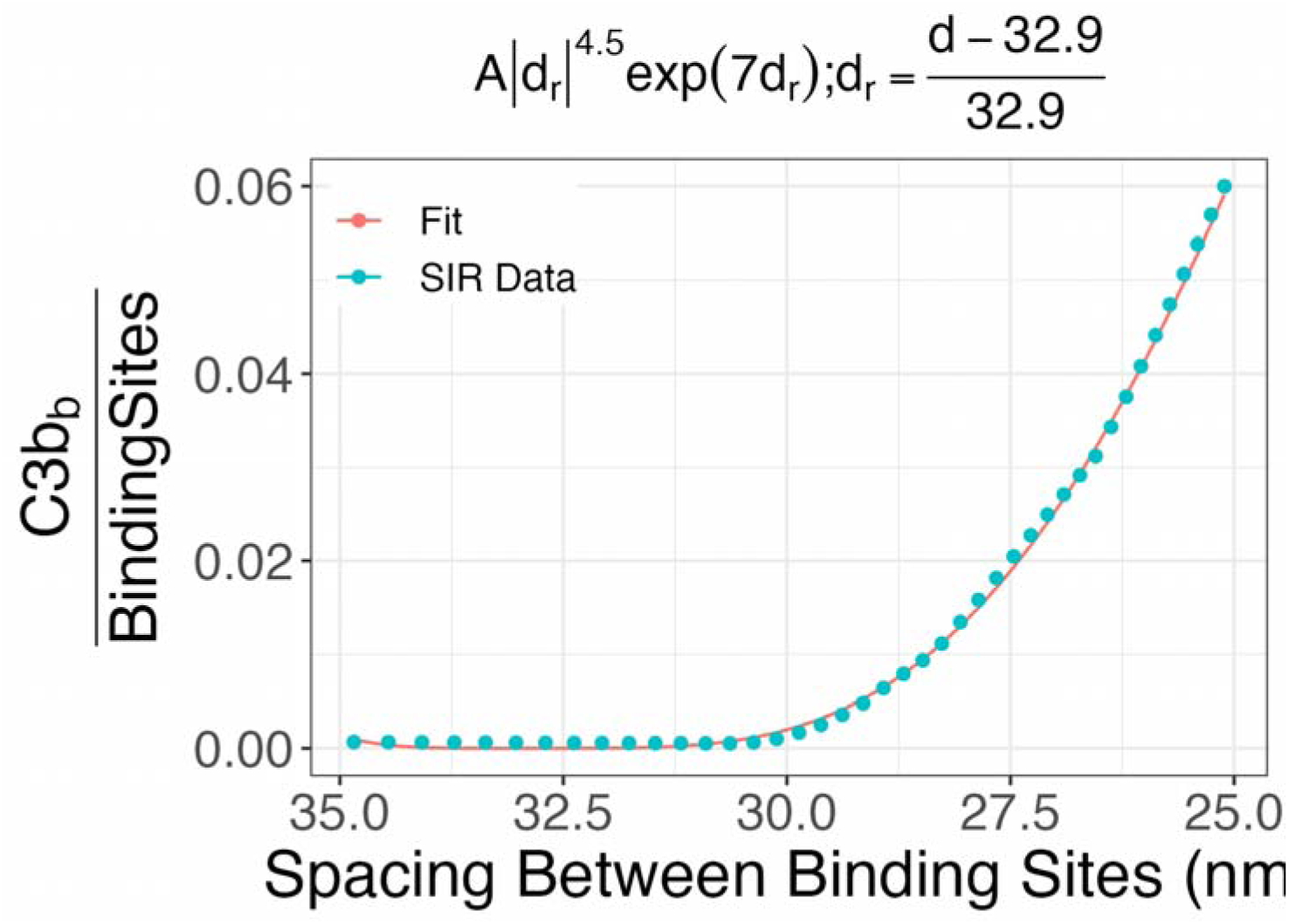
Critical Exponents explain threshold-like phenomena in SI-like model: fit for Surface Bound C3b.

**Figure SC13:**
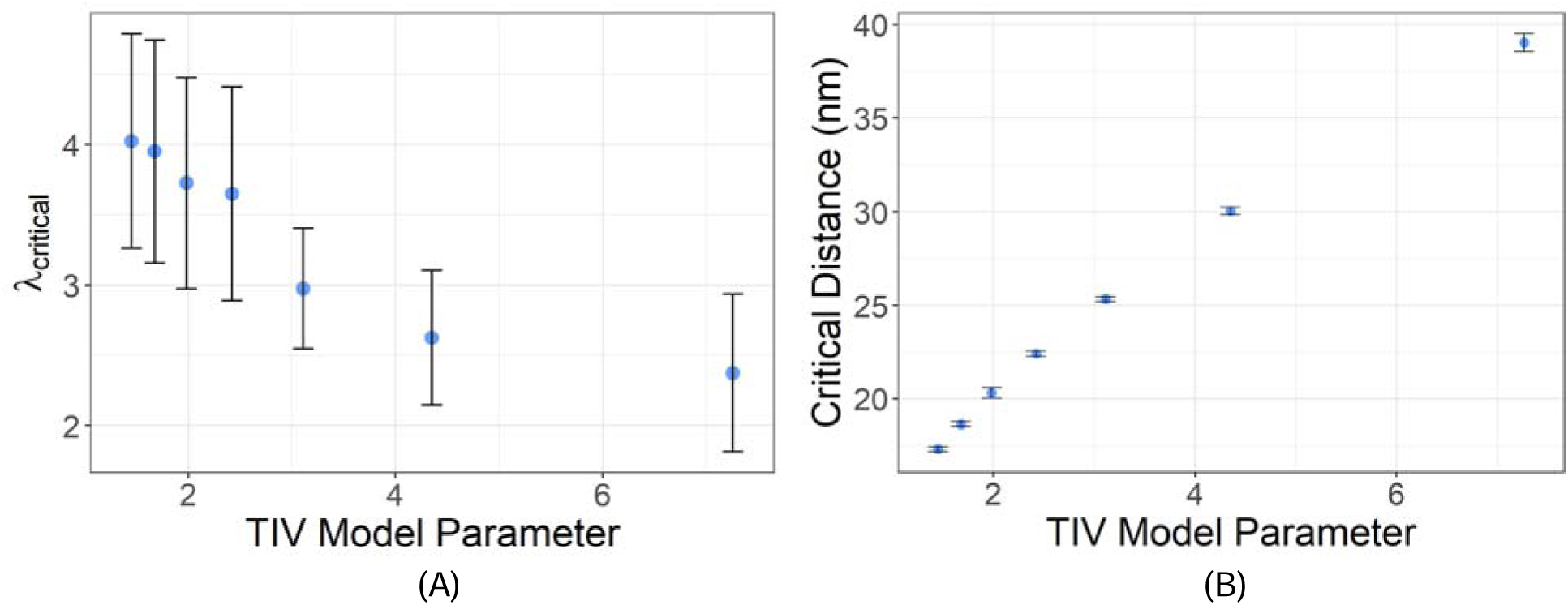
Varying TIV-like model parameter gives locus plots for (A) Critical exponent and (B) Critical Distance. From the error bars we can see that the critical exponents are sensitive to the domain of spacing (***d*** ∊ [***d_l_, d_u_***]) between binding sites used for fitting the critical exponent model. The model fitting was done using the maximum value of bound C3b concentration during the time course.

**Figure SC14:**
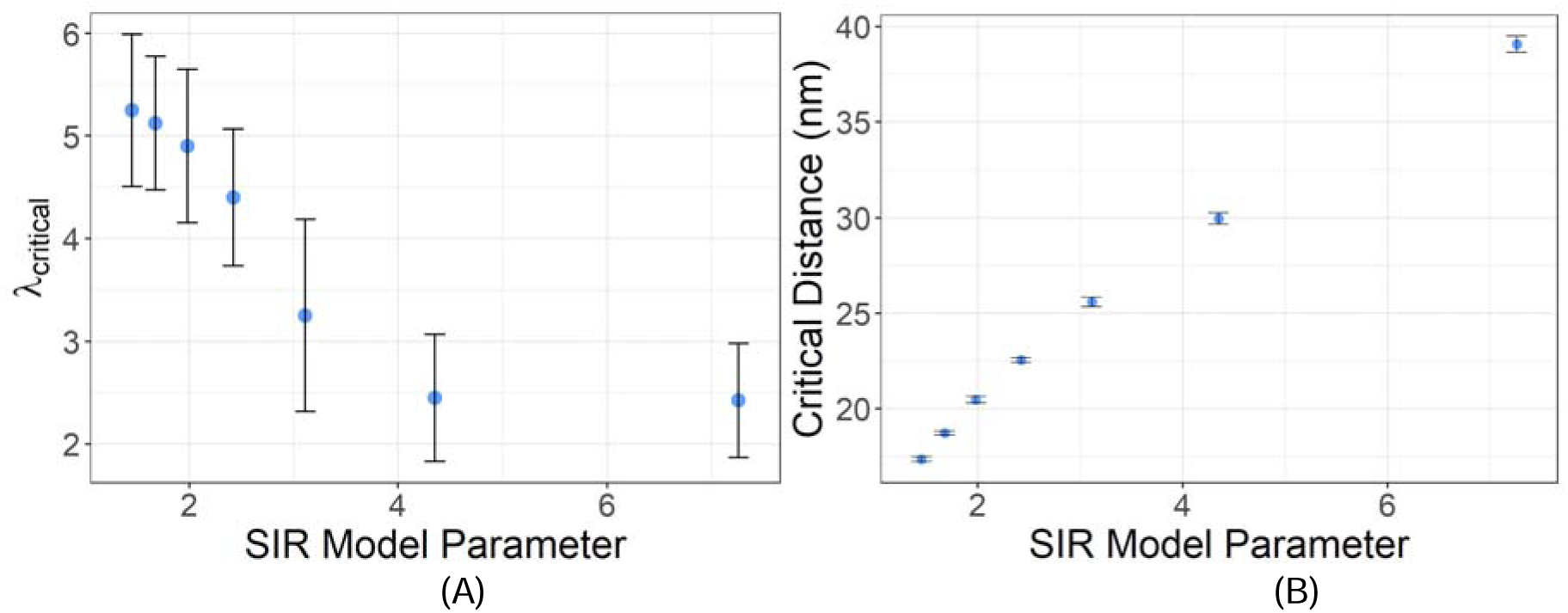
Varying SI-like model parameter gives locus plots for (A) Critical exponent and (B) Critical Distance. Once again, from the error bars we can see that critical exponents are sensitive to the range of spacing between binding sites used for fitting the critical exponent model. The values exponents and critical distance are in close proximity to the TIV-like model. The model fitting was done with the maximum value of bound C3b concentration during the time-course.

**Figure SC15:**
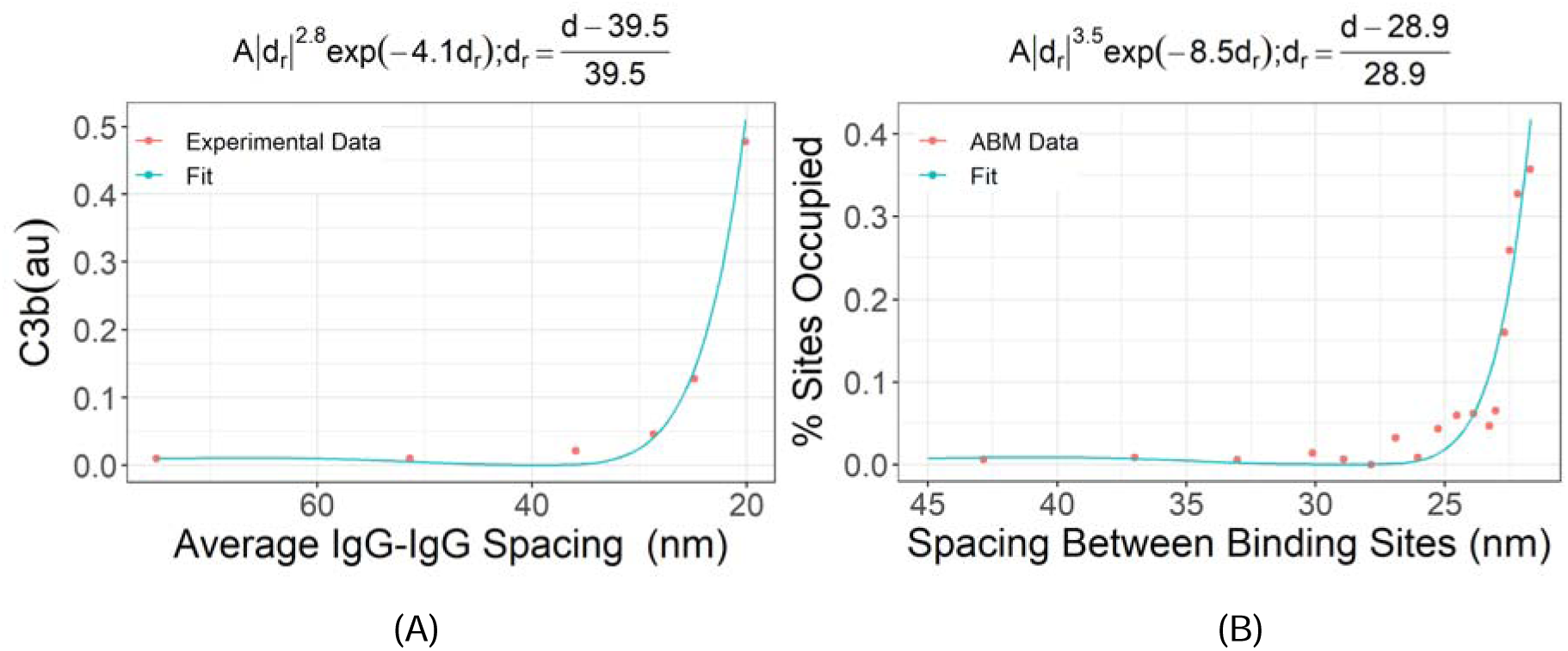
Critical Exponents explain threshold-like phenomena in. A) steady-state C3b data obtained from experiments and B)% of Sites occupied in ABM simulation. Both datasets have been normalized to be between 0 and 1.

## Notes

### Competing Interest Statement

The authors have declared no competing interest.

